# Transcription factors operate on a limited vocabulary of binding motifs in *Arabidopsis thaliana*

**DOI:** 10.1101/2023.08.28.555073

**Authors:** Sanja Zenker, Donat Wulf, Anja Meierhenrich, Sarah Becker, Marion Eisenhut, Ralf Stracke, Bernd Weisshaar, Andrea Bräutigam

**Affiliations:** Computational Biology, Faculty of Biology, Bielefeld University; Center of Biotechnology (CeBiTec), Bielefeld University; Genetics and Genomics of Plants, Faculty of Biology, Bielefeld University

**Author notes:** to whom correspondence should be addressed Contributions: SZ analyzed the data and co-wrote the manuscript, DW conceived of the study, AM produced and analyzed ampDAP-Seq data in *Marchantia polymorpha*, SB re-analyzed HY5 experiments, ME edited the manuscript, RS edited the manuscript, BW edited the manuscript, AB suggested analyses, interpreted data, and co-wrote the manuscript.

**Keywords:** transcription factor, binding motif, gene regulation, *Arabidopsis thaliana*, DAP-Seq

## Abstract

Predicting gene expression from promoter sequence requires understanding of the different signal integration points within a promoter. Sequence-specific transcription factors (TFs) binding to their cognate TF binding motifs control gene expression in eukaryotes by activating and repressing transcription. Their interplay generates complex expression patterns in reaction to environmental conditions and developmental cues.

We hypothesized that signals are not only integrated by different TFs binding various positions in a promoter, but also by single TF binding motifs onto which multiple TFs can bind. Analyzing 2,190 binding motifs, we identified only 76 core TF binding motifs in plants. Twenty-one TF protein families act highly specific and bind a single conserved motif. Four TF families are classified as semi-conserved as they bind up to four motifs within a family, with divisions along phylogenetic groups. Five TF families bind diverse motifs. Expression analyses revealed high competition within TF families for the same binding motif. The results show that singular binding motifs act as signal integrators in plants where a combination of binding affinity and TF abundance likely determine the output.

## Introduction

Transcription factors (TFs) govern transcriptional regulation in plants by controlling responses to a broad range of abiotic, biotic, and internal stimuli from temperature (Koini et al., 2009; Chung et al., 2020) over bacterial infection (Asai et al., 2002) to phytohormones (Matsuzaki et al., 2010; Powers et al., 2019). We frequently perceive TFs as on/off switches of gene expression as we are influenced, for example, by the pioneering work of Jacob and Monod with the *lac*-operon (Jacob and Monod, 1961) or the early ABC model of floral development (Haughn and Somerville, 1988; Coen and Meyerowitz, 1991).

Transcriptional regulation is also mediated by multiple general TFs, which form the transcription preinitiation complex with the RNA polymerase II (Orphanides et al., 1996; Roeder, 1998; Kadonaga, 2012). The regulatory region of a eukaryotic gene generally consists of the core promoter and multiple proximal or distal regulatory regions, e.g., enhancers, silencers, and insulators (Levine and Tjian, 2003). Besides the elements recognized by the general TFs at the core promoter, these regions also contain TF binding motifs (TFBMs), which are DNA sequences that are bound by sequence-specific TFs. The type and position of TFBMs on the DNA in regulatory regions spell out a code, which is read out by the TFs (Seeman et al., 1976; Rohs et al., 2009; O’Malley et al., 2016). If the readout critically depends on positional information and on the completeness of an array of TFBMs, the system acts like an enhanceosome (Arnosti and Kulkarni, 2005). If elements of the enhancer can be read out individually, the system acts like a billboard (Arnosti and Kulkarni, 2005). In both cases, the regulatory regions consisting of multiple TFBMs integrate signals to provide a quantitative output. Deep learning has been used to predict TFBMs (Li et al., 2018) and to predict gene expression from DNA sequence features (Avsec et al., 2021b).

The TFBMs of TFs are hypothesized to be a main component to binding specificity and to guide the regulatory activity of TFs (Weirauch et al., 2014). TFBMs can be characterized *in vivo* and *in vitro*. For *in vivo* characterization, chromatin immunoprecipitation sequencing (ChIP-Seq) is most frequently used. This method is influenced by additional *in vivo* factors, such as chromatin structure and partner proteins (Gordân et al., 2009; Li et al., 2011). In contrast, *in vitro* methods, such as protein binding microarrays, high-throughput *in vitro* selection, and DNA affinity purification sequencing (DAP-Seq) allow derivation of TFBMs from pure DNA binding events (Berger et al., 2006; Jolma et al., 2013; O’Malley et al., 2016). Some studies have demonstrated that similar TFs have similar binding specificities (Rushton et al., 1995; Berger et al., 2008; Weirauch et al., 2014; Galli et al., 2018) while other studies show that small changes in TF amino acid sequence lead to changes in binding specificities (Cook et al., 1994; Noyes et al., 2008; Aggarwal et al., 2010). TFs are grouped into families based on their shared DNA binding domains (DBDs) and additional domains for e.g., protein interaction (Wilhelmsson et al., 2017). Very large-scale cross-kingdom analyses have suggested that TF binding specificity to particular TFBMs is predicted by DBD amino acid sequence similarity and that extensive similarity in binding can be detected (Weirauch et al., 2014). In *Saccharomyces cerevisiae*, however, 60% of TFs have evolved differential binding preferences due to variations mainly outside the DBD (Gera et al., 2022). In contrast, analyses of TFBMs in *Drosophila melanogaster* and *Homo sapiens* have shown striking conservation between structurally similar TFs despite expansion and divergence of families over the span of 600 million years (Nitta et al., 2015). Overall, in animals, extensive conservation is detected in some TF families but not all. The TF family with C2H2 as the DBD, for example, shows much higher variation in TFBMs despite high protein sequence conservation (Lambert et al., 2019). In multicellular organisms, TFs often comprise a larger proportion of the proteome than in unicellular organisms accounting for more than 2% of proteins; seed plants generally reach above 4% (Lang et al., 2010; De Mendoza et al., 2013). In different origins of multicellularity and therefore in different origins of complex development, different families of TFs expanded and still expand, but the evolutionary source of the TF family frequently predates the expansion event (De Mendoza et al., 2013). Expansion correlates with phylogeny as phylogenetic branches share specific family expansion patterns (Lang et al., 2010; De Mendoza et al., 2013). TFs identified in model organisms range from about 750 to 1,600 in number (Riechmann et al., 2000; Gray et al., 2004; Reece-Hoyes et al., 2005; Adryan and Teichmann, 2006; Lambert et al., 2018). The expansion of TF families is caused by whole genome duplications (WGD), local tandem duplications, and more distant duplication events. Duplication allows for sequence and function divergence under relaxed selective pressure (Ohno, 1970; Zhang, 2003). WGD evidently leads to higher retention rates for duplicated genes by balancing detrimental gene dosage effects potentially created by single copy duplications (Edger and Pires, 2009; Schmitz et al., 2016). Although most duplicated copies of genes become non-functional within a short evolutionary time by accumulating deleterious mutations leading to pseudogenization (Lynch and Conery, 2000), some gene copies neofunctionalize and adopt completely new functions compared to their paralog. Alternatively, both copies subfunctionalize to cover parts of the function of the original version (Ohno, 1970; Zhang, 2003). In contrast to animals, most plant genera are tolerant to autopolyploidy and allopolyploidy, which may provide an opportunity for different TF family expansions and for extensive neofunctionalization.

In plants, there are several well described examples of TF families that have expanded. The MYB superfamily, for example, can be found across all Eukaryota, but has increased by 9-fold in member number from *Chlamydomonas reinhardtii* to *Arabidopsis thaliana* (Feller et al., 2011; De Mendoza et al., 2013). MYB TFs are defined by their DNA-binding MYB-domain, which consists of a variable number of (imperfect) MYB repeats, each forming three α-helices, with the second and third forming a helix-turn-helix structure to interact with the DNA binding site. The MYB family can be divided into subfamilies based on the number of MYB repeats (Stracke et al., 2001). The most abundant family of MYB TFs in the plant kingdom are the R2R3-MYBs. Some R2R3-MYB proteins like MYB66/WEREWOLF have been shown to bind the TFBM AACNGC (Wang et al., 2019), while other MYB proteins recognize ACC(T/A)G or TAAC (O’Malley et al., 2016), making this family an example for partially conserved binding specificity. The WRKY TF family has also expanded within the Embryophyta clade. Members of this family are defined by the presence of at least one WRKY domain consisting of approximately 60 amino acids including zinc finger motifs and the conserved WRKYGQK region, which directly contacts the W-box as its TFBM (Eulgem et al., 2000; Rinerson et al., 2015). Although many WRKY TFs are involved in different responses to abiotic and biotic stresses, strong conservation of TFBMs has been reported (Eulgem et al., 2000) making this family an example for a large TF family with a highly conserved TFBM. Many additional TF families and subfamilies, such as basic helix-loop-helix (bHLH) factors have evolved before the water-to-land transition (Catarino et al., 2016; Jin et al., 2017) and expanded in land plants. Fewer TF families, such as ARID, E2F/DP, or GRAS have more or equal members in mosses compared to angiosperms (Feller et al., 2011; De Mendoza et al., 2013; Wilhelmsson et al., 2017). The general increase in TF numbers can be explained by more frequent occurrence of WGD in plant lineages compared to animals and higher rates of parallel expansion in common TF families (Shiu et al., 2005). Results from large-scale efforts to characterize TFBMs (Franco-Zorrilla et al., 2014; O’Malley et al., 2016) indicate that there is extensive conservation of TFBMs within TF families.

Here, we explored *in vitro* TFBM data from plants and showed that experimentally verified binding of multiple TFs on identical sites is detectable. We hypothesized that common binding motifs are a product of TF family expansion and tested binding preferences along evolutionary trajectories including dating of diversions with TFs from the bryophyte *Marchantia polymorpha*. We show that the vocabulary of TFBMs is comparatively small and hypothesize that, if expression patterns overlap, there must be considerable competition for binding at a given relevant TFBM. Indeed, the data demonstrates that expression similarity of TFs with the same TFBM contribute to the potential for signal integration at single TFBMs. Taken together, the results suggest that in addition to the well-known signal integration mediated by TF arrays on a plant promoter, the binding preferences and expression patterns of TFs provide ample substrate for signal integration at single TFBMs to control promoter output.

## Results

To gain an overview of experimentally validated TF binding events in the genome of *A. thaliana, we* mapped these onto individual promoters of genes, here defined as the 1kb region upstream of the ATG using narrowPeak DAP-Seq data from O’Malley et al. (2016) (Figure 1A). Visualizations were amended with accessible chromatin regions from five publications (Lu et al., 2017; Maher et al., 2018; Sijacic et al., 2018; Sullivan et al., 2019; Lu et al., 2019) to visualize accessibility and overlaid with conservation of sequence information based on the 1001 genome project (Alonso-Blanco et al., 2016). It is immediately apparent that peak stacks can be observed over defined promoter regions (Figure 1A) showing substantial overlap in TFs binding to the same TFBM. Many of these TFBMs are indeed accessible in three different tissues (Figure 1A). Sequence conservation of these TFBMs in the *A. thaliana* pangenome is high, as validated by low sequence variation detected by the 1001 genome project (Figure 1B) in regions with utilizable data (Figure 1C). Analyzing 27,206 promoters of nuclear protein coding genes in *A. thaliana*, there is experimental evidence for 0 to 219 (one outlier with 786) binding events with median number of 41 binding events and on average 43 binding events per promoter (Figure 1C, https://doi.org/10.4119/unibi/2982196). This analysis of individual promoters qualitatively shows that there is indeed large positional overlap in TF binding and therefore likely large overlap in TFBMs.

**Figure 1.**
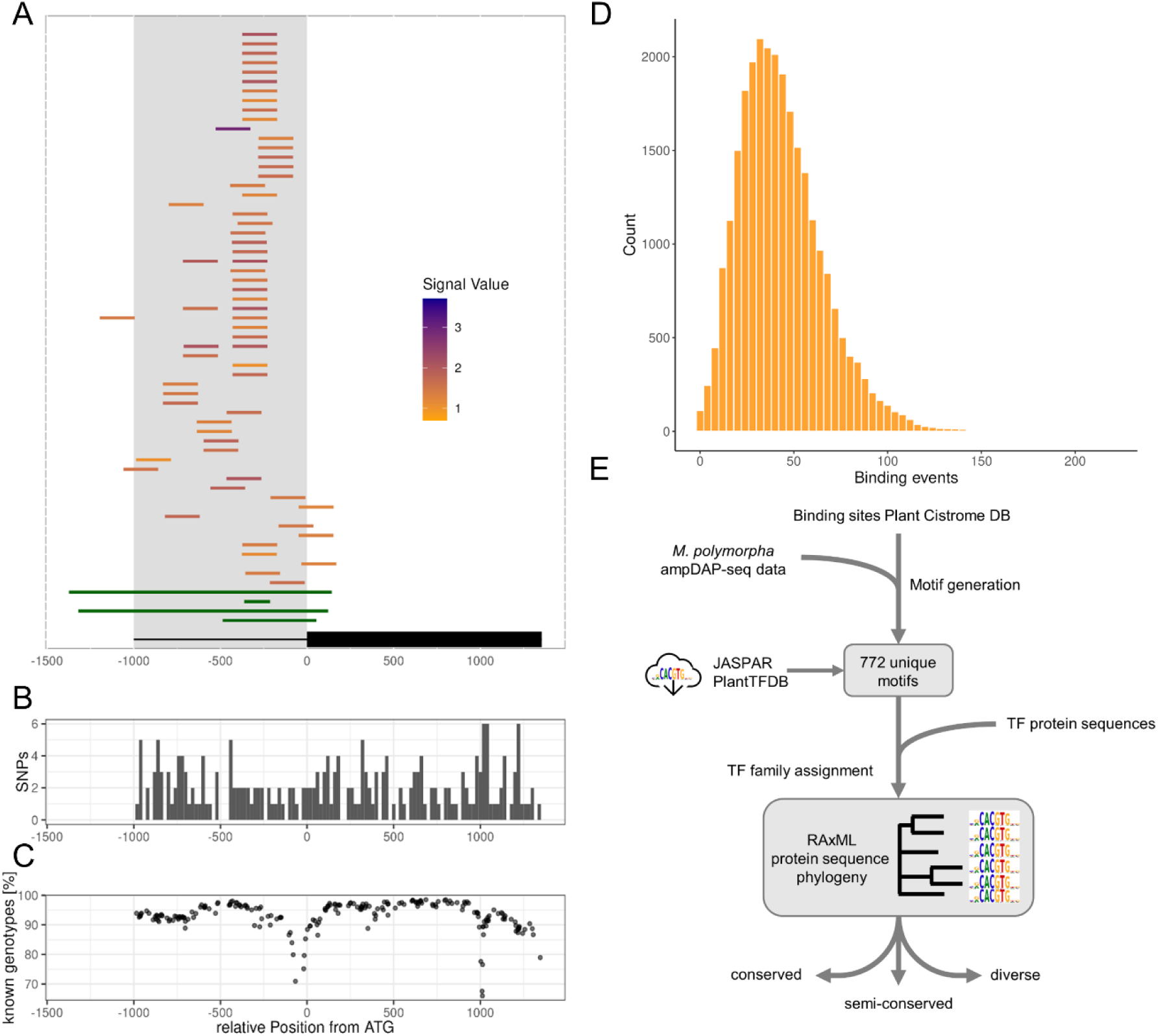
T**F**BM **conservation analysis. A:** Representative visualization of DAP-Seq binding events, colored from orange to purple by log10-tranformed peak height and open chromatin regions (green) on the *AT2G46310* promoter (all promoter figures as interactive visualizations deposited under https://doi.org/10.4119/unibi/2982196). **B:** Conservation of the *AT2G46310* promoter nucleotide sequence in *A. thaliana* pangenome visualized by position dependent SNPs. **C:** Pangenome data density visualized by percentage of successfully called genotypes at the SNP positions of the *AT2G46310* promoter. **D:** Histogram showing the number of experimentally determined TF binding events on nuclear protein coding gene promoters (one outlier with 786 binding events not shown). **E:** Workflow for TFBM conservation analysis.

To quantify TFBM identity and similarity across TF families in plants, we retrieved all available TFBMs in databases (Jin et al., 2017; Castro-Mondragon et al., 2022) and to reduce bias, we determined one representative TFBM for each TF through a common pipeline, preferring ampDAP-Seq based TFBMs due to the coverage of all genomic binding sites without methylation influence (Figure 1E). All peaks from the Plant Cistrome database (O’Malley et al., 2016) and, to analyze evolutionary trajectories, peaks from 14 *M. polymorpha* TFs were obtained and TFBMs generated using MEME-ChIP (Machanick and Bailey, 2011). All TFs with a TFBM were grouped into families based on their DBDs according to TAPscan TF family definitions for *A. thaliana* (Wilhelmsson et al., 2017), aligned and phylogenetically clustered. We then evaluated each TF family based on how many different TFBMs exist within this family and classified the families into either conserved, semi-conserved, or diverse (see methods for details, see Table S1).

Database queries yielded a total of 2,190 redundant data points from the Plant Cistrome (O’Malley et al., 2016), JASPAR (Castro-Mondragon et al., 2022) and PlantTF (Jin et al., 2017) databases (Table 1). The majority of known plant TFBMs are present in multiple databases, leading to extensive redundancy. After removal of redundant entries, 772 different TFs from 14 different plant species and 50 TF families remained. Data density for plants other than *A. thaliana* is quite low. Of the 1,725 DNA-binding TFs in *A. thaliana*, 680 have a known TFBM (Table 1). These 680 represent 50 of 71 annotated families in the TAPscan database (Wilhelmsson et al., 2017). Within these 50 families, we know on average 40.6% of all annotated TFBMs in *A. thaliana*. The databases also contain 18 TFBMs, which could not be assigned to a TAPscan TF family. We tested for overall TFBM identity and similarity and found that the 772 TFBMs represent 76 different core TFBMs, which are between 5 and 21 bp in length with an average length of 8.9 bp (Table 1). Some TFBMs like the WRKY W-box TTGAC TFBM are limited to one specific family, while the E-/G-box (C)ACGTG can be found as a TFBM for BES1, bHLH, bZIP family members, and one Trihelix family member (see Table S2). Taken together, the currently known promoter vocabulary of TFBMs consists of 76 distinct TFBMs in plants with most of the syntax (positioning, interaction, affinity, etc.) unknown.

**Table 1.** Overview of general TFBM statistics.

| Feature | Number |
| --- | --- |
| Total TFBMs JASPAR | 757 |
| Total TFBMs Plant Cistrome DB | 814 |
| Total TFBMs PlantTFDB | 619 |
| Total TFs (*A. thaliana*) | 1725 |
| TFs with TFBM (*A. thaliana*) | 680 |
| TFs with TFBM (other species) | 92 |
| Distinct core TFBMs | 76 |
| Average TFBM length [bp] | 8.9 |
| TF families (*A. thaliana*) | 71 |
| TF families with TFBM (*A. thaliana*) | 50 |
| TFBMs not assigned to TAPscan family | 18 |
| Average TFBMs known per family (*A. thaliana*) [%] | 40.6 |

### Many families show high levels of TFBM conservation

The majority of TFBMs are available for *A. thaliana* (680), followed by *Zea mays* (37). The TF families with the most known TFBMs are the AP2/EREBP family (107) and the MYBs of R2R3– and 3R-MYB families (67). Phylogenetic analyses were performed on amino acid sequences for 32 families, which contained at least four TFs with TFBMs to assess the relationship between amino acid sequence and TFBM similarity. To date the age of a phylogenetic subgroup, we assessed the presence of *M. polymorpha* and *C. reinhardtii* orthologs determined by Orthofinder2 (Emms and Kelly, 2019) which, if present, indicate an age of at least 500 or 950 million years, respectively, for the branch (Hedges et al., 2004; Harris et al., 2022).

Based on the low number of distinct TFBMs (Table 1) and the literature (Ciolkowski et al., 2008), we hypothesized that some TF families retain conserved TFBMs despite extensive family expansion dating back at least 500 million years. A phylogeny of the WRKY TFs resolved six subclades of which all except clade II-a contain at least one *M. polymorpha* ortholog (Figure 2A). We confirmed the known TFBM conservation in the WRKY family, with TFBMs available for 45 out of 73 members in *A. thaliana,* all binding the TFBM TTGAC across all clades with minimal variation in the flanking sequences (Figure 2A). One WRKY TF ortholog from *C. reinhardtii* with a known TFBM was identified belonging to subclade I, which has been proposed to be the oldest group, from which all others potentially evolved (Rinerson et al., 2015). Subclade II-d has an additional annotated Zn cluster domain, which does not appear to influence the recognized TFBM (Figure 2A). In principle, all WRKY TFs compete for the same TFBM (Figure 2A). To test whether competition is allowed via similar expression patterns, we compiled expression data from 6,033 RNA-Seq experiments and tested for expression similarity by Pearson correlation. Of the 45 family members with a TFBM in *A. thaliana*, 15 share expression with at least one other family member based on a correlation coefficient of >0.7. 33 have similar expression with at least one other family member based on a correlation coefficient of at least >0.5 (Figure 2C).

**Figure 2.**
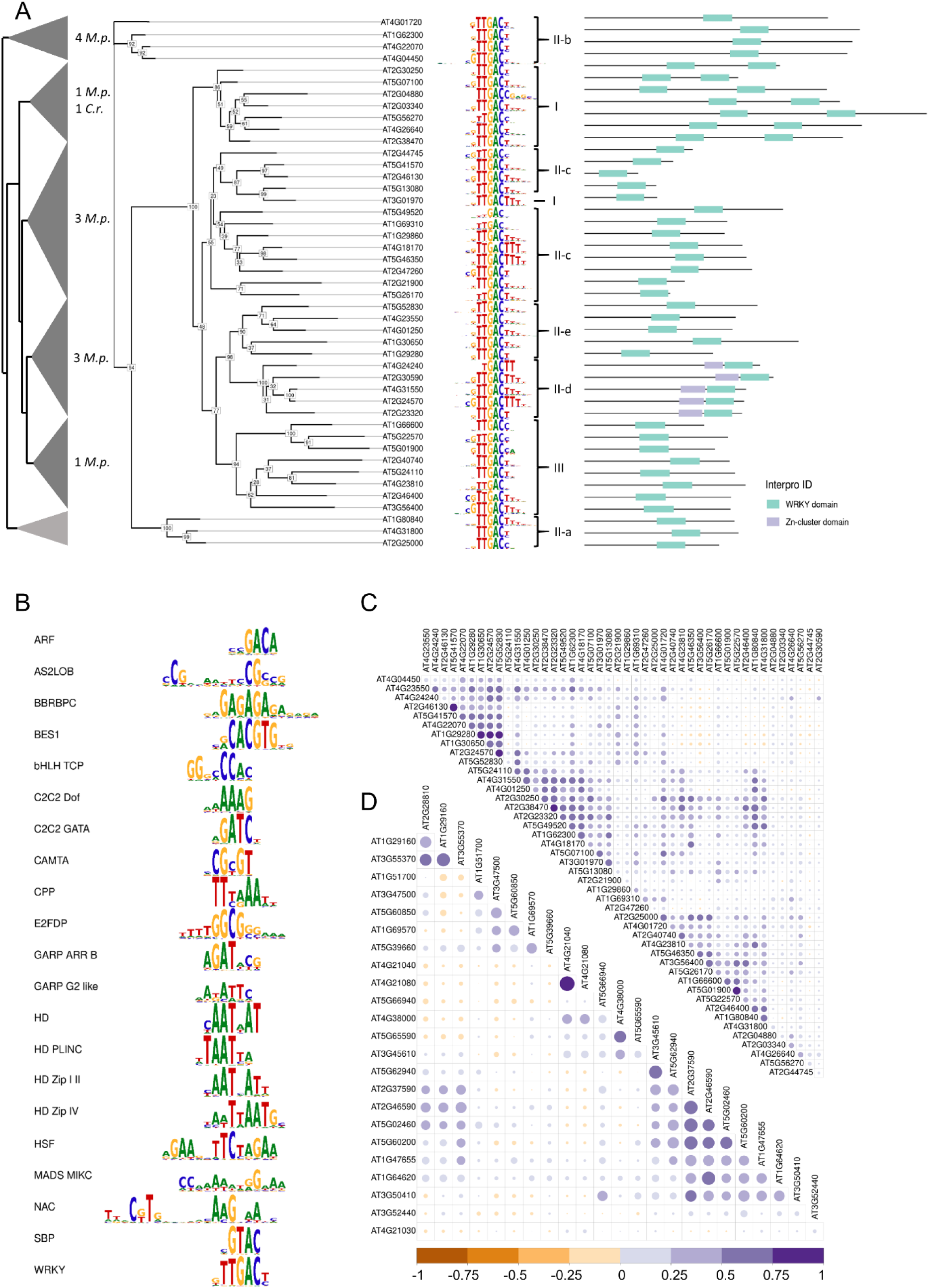
A**n**alysis **of TF families with high motif conservation. A:** Unrooted phylogenetic tree of the WRKY family TFs with known TFBMs. Support values at the nodes are based on 1,000 bootstrap iterations. Clade annotations are from Eulgem et al. (2000) and Interpro domain annotations. Collapsed phylogenetic tree are shown with indication of orthologues genes from *M. polymorpha* or *C. reinhardtii* in each subgroup. **B:** All TF families with consensus TFBMs of all families are considered as conserved. Base height corresponds to information content. **C:** Expression correlation of 45 WRKY family members in *A. thaliana*. The correlation coefficient is indicated by color and dot size. **D:** Expression correlation of 24 C2C2 Dof family members in *A. thaliana*.

Examination of the other 31 TF families (see Figures 3A, 5, S1-S29) with at least four characterized TFBMs revealed that 20 additional TF families show extensive TFBM conservation within the TF family (Figure 2B). For example, TFs of the C2C2 Dof zinc finger family have been identified in *A. thaliana*, *Z. mays,* and *Physcomitrium patens* and unanimously bind the consensus TFBM AAAG with little variation in the adjacent bases (see Figure S10). One *A. thaliana* TFBM stands out, as the AAAG is followed by the reverse TFBM CTTT, indicating a potential binding by a TF dimer, which was not picked up by MEME-ChIP for other members of the family (see Figure S10). One ortholog was found in *M. polymorpha* and one in *C. reinhardtii*, suggesting conservation of this TFBM since approximately 950 mya (Hedges et al., 2004). Expression analyses in the C2C2 Dof family show that two family members share expression with at least one other family member based on a correlation coefficient of >0.7. Thirteen C2C2 Dof TF family members have similar expression with at least one other family member based on a correlation coefficient of at least >0.5 (Figure 2D), which indicates potential for competition at a shared TFBM.

**Figure 3.**
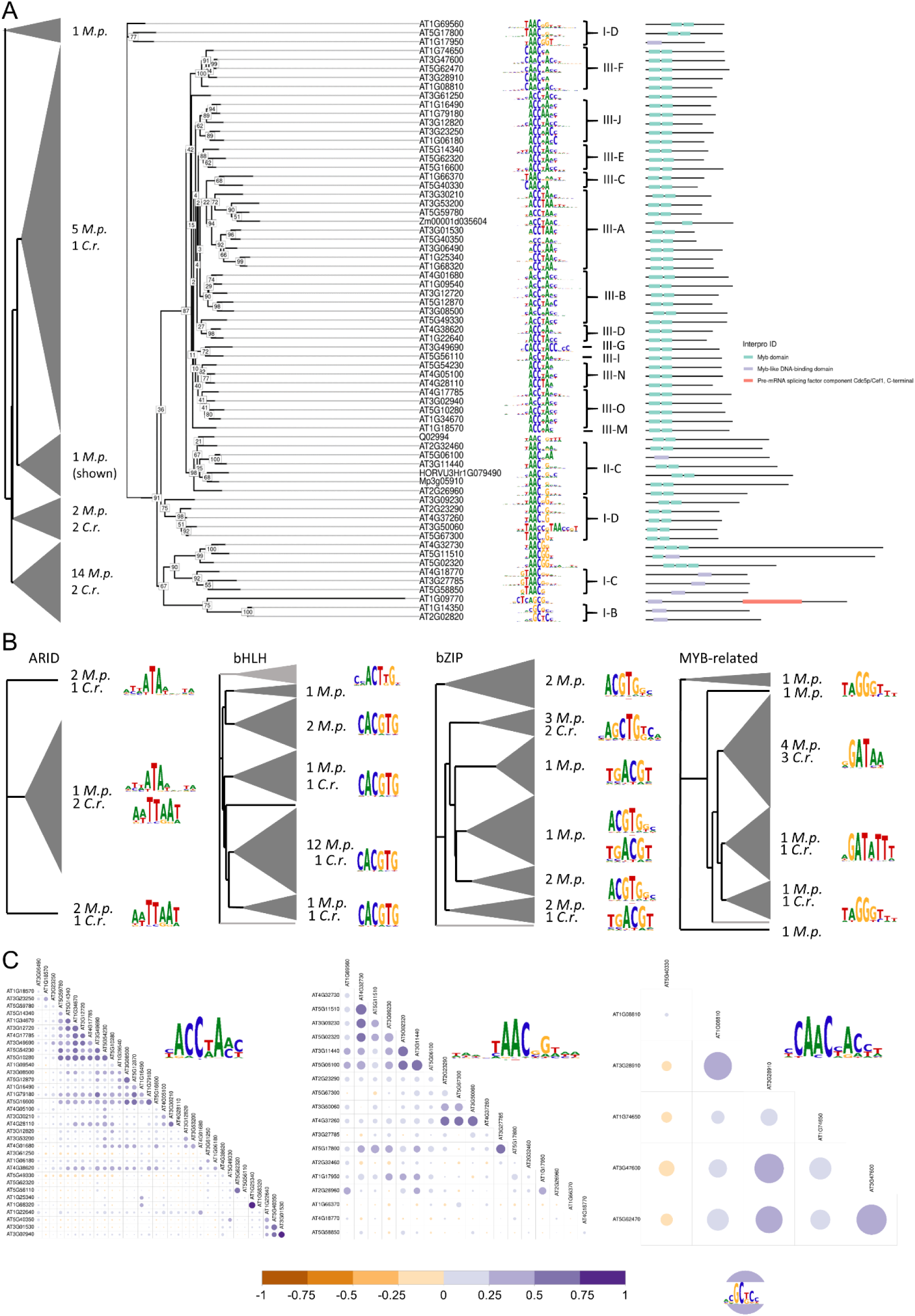
A**n**alysis **of TF families with semi-conserved TFBMs. A:** Phylogenetic tree of the MYB family with support values based on 1,000 bootstraps. Interpro domain annotations indicate structural similarities. Clade annotations are from (Chang et al., 2020) and domain annotations are from Interpro. Collapsed phylogenetic tree with indication of orthologues proteins from *M. polymorpha* or *C. reinhardtii* in each subgroup. **B:** Collapsed phylogenetic relations with orthologues proteins found. Light grey indicates orthologues were not found. TFBMs represent the consensus TFBM of the clade. **C:** Correlation of expression for the given TFBM subgroups.

Not all TF families with high conservation of TFBMs have existed since 500 mya and some can only be found in land plants. One example is the ARF family, which is defined by a B3 binding domain, an ARF domain, and in most members also an AUX/IAA domain (see Figure S3). All analyzed members share the conserved TFBM GACA, which is either preceded by a C or a G-stretch for many members (Figure 2B). These differences in surrounding bases have been postulated to arise from different genomic context of the TFBMs (Galli et al., 2018). Expression analysis in the ARF family shows that members do not share expression based on a correlation coefficient of >0.7. However, five out of six members in *A. thaliana* have similar expression with at least one other family member based on a correlation coefficient of >0.5 (see Figure S30) indicating that not all TF families contain members in intense competition for binding sites.

Among the 21 TF families, which represent the highest level of TFBM conservation, 21 of the 76 different possible TFBMs are represented (Figure 2B). The expression divergence of family members within each family (see Table S3), shows that for each of the 21 TFBMs, TFs are in some degree in competition for binding. In the analysis, we identified a total of 21 families with one conserved family TFBM (Figure 2B), pointing to a large constraint in the *de novo* evolution of TFBMs as a way for neofunctionalization of TFs in at least these families.

### Semi-conservation of TFBMs follows phylogenetic relationships

Not all TF families are under a similarly strict constraint for their member TFBMs. We expected that some families acquired more variation in the recognized TFBM as a potential way to neofunctionalize and regulate different pathways in more than 500 million years of evolution. Our analysis revealed a continuous transition between completely conserved families binding to one single TFBM (Figure 2B) and families that bind up to 14 different TFBMs across 35 analyzed members (C2H2) (see Figure S13). Therefore, we established the classification of semi-conservation to cover families with up to four different consensus TFBMs and no more than 15% outliers (TFBMs bound by only one member). This applied to the five families ARID, bHLH, bZIP, MYB, and MYB-related (see Table S1).

Many TFBMs of MYB TFs have been identified in *A. thaliana*, but there is also data available from *Hordeum vulgare*, *Z. mays*, and *Petunia hybrida* (Figure 3A) in openly accessible databases. We additionally determined the TFBM of one MYB TF in *M. polymorpha* using ampDAP-Seq. The phylogeny of the MYB TFs with known TFBM (Figure 3A) recaptures published groupings of the MYB family and breaks along protein domain lines (Figure 3A, Chang et al., 2020). When the TFBMs are overlaid over the phylogeny, they appear grouped along the phylogenetic tree corresponding to previously annotated clades (Chang et al., 2020), including non-*A. thaliana* TFBMs. The TFBMs thus reflect phylogeny, highlighting conservation of these TFBMs across selected plant species. We found a total of 15 *M. polymorpha* orthologues MYB TFs in subclade I, two additional ones in subclade II and five more in subclade III, demonstrating that these clades existed at least 500 mya. Four orthologs from *C. reinhardtii* are found in subclade I and, unexpectedly, one in subclade III, which was previously found to be unique to embryophyta (Chang et al., 2020). The MYB family has one known outlier, *AT1G09770 (CDC5)*, which is visibly more distant at the amino acid level and has a TFBM that is not found for other MYB TFs or MYB-related TFs. CDC5 has two MYB repeats with a 31% identity to the typical R2R3-MYB domains (Stracke et al., 2001). Taken together, we can conclude that different TFBMs in the MYB family have existed since at least 500 mya followed by expansion of the different subclades. Within each subclade, the MYB TFs potentially compete for their respective TFBM. Expression analysis in *A. thaliana* show that within the 35 members binding the consensus TFBM ACC(T/A)A, nine share expression with at least one other family member based on a correlation coefficient of >0.7. Nineteen TFs have similar expression patterns with at least one other family member based on a correlation coefficient of >0.5 (Figure 3C). Among the 19 members binding the consensus TFBM TAAC, 12 have similar expression with at least one other family member based on a correlation coefficient of at least >0.5 (Figure 3C). Among the 6 members binding the consensus TFBM CAACNAC, none have shared or similar expression patterns with at least one other family member (Figure 3C). The two members binding the TFBM GCTC have distinct expression patterns with a correlation coefficient <0.5 (Figure 3C). Although the higher level of divergence of the TFBMs reduces the potential for competition, we detected similar expression patterns between at least two members binding the same TFBM for 40 TFs, indicating that, also in the MYB family, TFs compete for the same TFBM.

The bHLH family members bind two different TFBMs that differ predominantly in two base positions. The initial C of the G-box TFBM CACGTG is missing in the second variant and the first G is replaced by a T, leaving a core of ACTTG, which was found to be bound by four bHLH TFs (Figure 3B). The bHLH TFs binding to ACTTG are more distant from all other members in the tree and the only subgroup with no orthologs in *M. polymorpha* or *C. reinhardtii* (Figure 3B). Again, expression analysis suggests potential competition of different bHLHs for the same TFBM. Within the 28 *A. thaliana* bHLH TFs binding the G-box CACGTG, four share expression patterns with at least one other bHLH family member based on a correlation coefficient of >0.7 and 13 have similar expression patterns with at least one other family member based on a correlation coefficient of at least >0.5 (Figure S50). The four members binding ACTTG have distinct expression patterns with a correlation coefficient <0.5 (Figure S50).

Grouping of TFBMs along the phylogenetic relationship is also observed in the bZIP family and the MYB-related family (Figure 3B), suggesting stability of TFBMs during clade-specific expansion. At the same time, expression analyses showed that there is extensive sharing of expression patterns within group members binding the same TFBM (see Table S3, Figures S49-S52). This group of five semi-conserved TF families covers 18 of the 76 core TFBMs including TFBMs that are only bound by one member in a family and were excluded from consensus TFBM generation. The level of competition for each TFBM is reduced compared to the TF families for which the TFBMs are highly conserved with a lower number of TFs who share expression or have a similar expression pattern. The analysis of phylogeny compared to binding specificity points to a reduced constraint of evolvability of binding specificity compared to the highly conserved TFs.

### Diverse TF families bind a variety of different TFBMs

We identified five TF families that have more than four different consensus TFBMs or more than 15% outliers and classified them as diverse. The diverse TF families are those that contain the DBDs of the type ABI3/VP1, AP2/EREBP, C2H2, C3H and Trihelix (see Table S1).

Trihelix TFs have only been identified in land plants, suggesting that the family evolved after the transition from water to land. Members of this family have one or two binding domains similar to the MYB domain and are therefore called MYB/SANT-like (Figure 4). The Trihelix family is divided into five clades based on amino acid sequence similarities and the phylogeny largely recapitulates the grouping (Kaplan-Levy et al., 2012) (Figure 4). As of now, TFBMs are available for 14 out of 26 members in *A. thaliana*, and we generated one TFBM using ampDAP-Seq in *M. polymorpha* of a member of the SIP1 clade. All previously defined clades of the Trihelix family have at least one known TFBM (Figure 4). The GT-2 clade has a conserved TFBM of TTTAC. Two TFs in the GT-1 clade, as well as one TF in the SIP1 clade bind to the known GT-element. The third TFBM from the GT-1 clade for TF GT-3a (AT5G01380) is known as the G-box (CACGTG) (Figure 5) (Ayadi et al., 2004). Despite the observed TFBM similarity within clades, members of the Trihelix family TFs bind to a wide range of TFBMs across and within clades.

**Figure 4.**
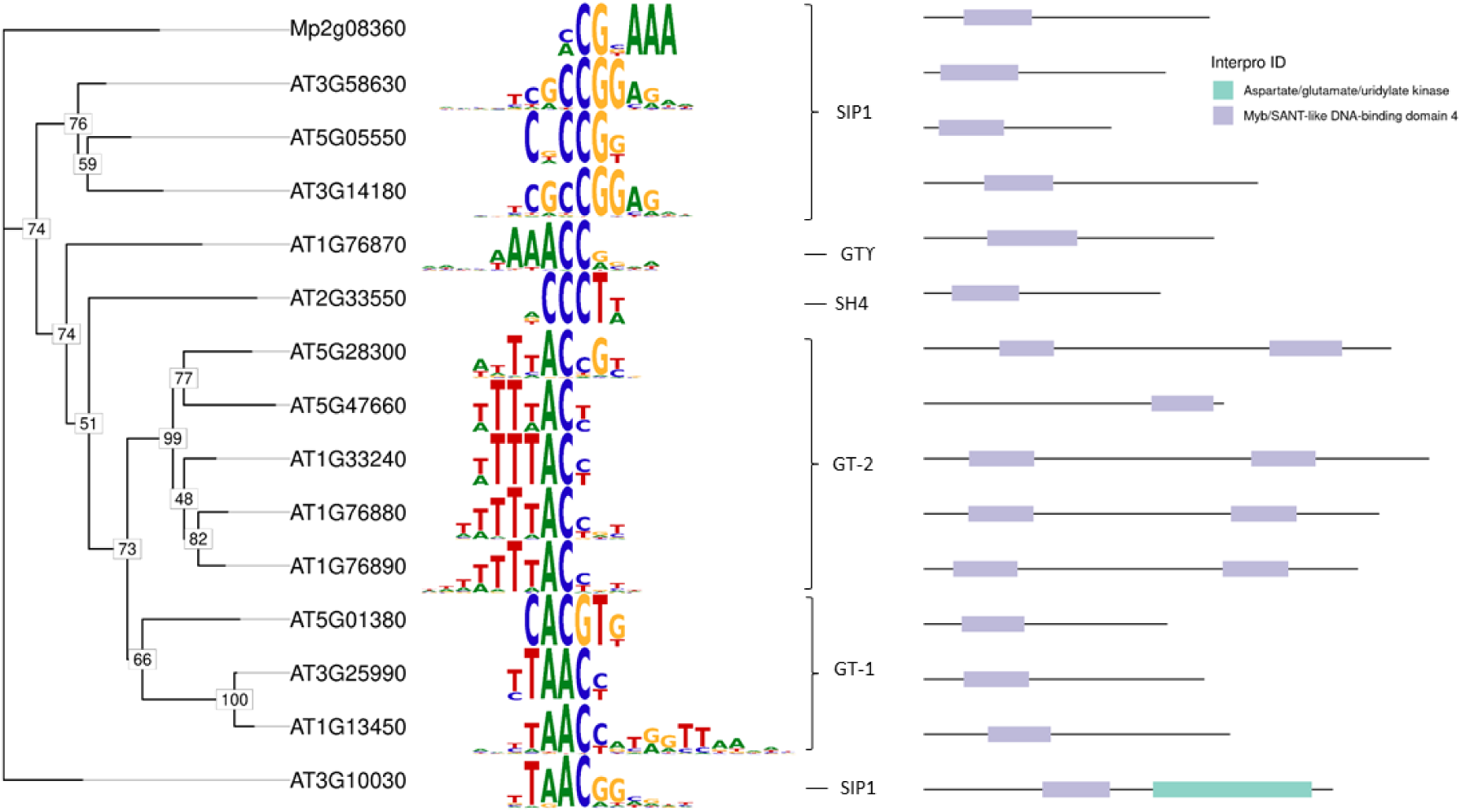
T**h**e **Trihelix family is an example of a TF family with diverse binding motifs.** Unrooted phylogenetic tree of the Trihelix TFs with known TFBMs. Support values at the nodes are based on 1,000 bootstrap iterations. Clade annotations from Kaplan-Levy et al. (2012) and domain annotations are from Interpro.

**Figure 5.**
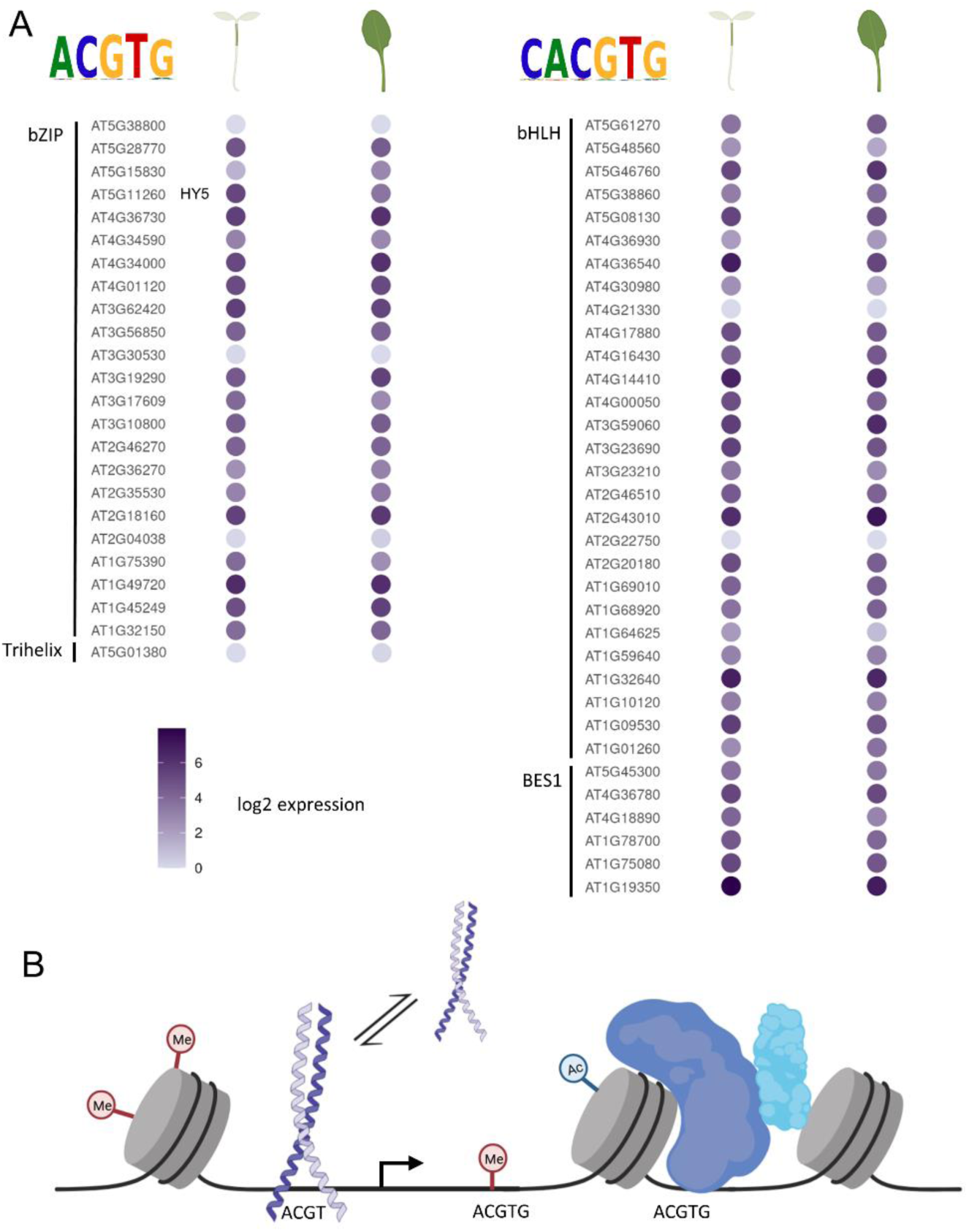
T**F**s **compete for the same binding motif. A:** Log2-scaled expression of TFs in the hypocotyl and leaf that bind the according consensus TFBM. HY5 (AT5G11260) is underlined. **B:** Graphical model of factors influencing HY5 binding at a given TFBM, including the dissociation constant (K_D_), histone modification, DNA methylation, chromatin structure, TFBM context and position.

In this analysis, we also identified a high diversity of TFBMs bound by the C2H2 TF family members in plants (see Figure S12), as it has been reported for fruit fly and human C2H2 TFs (Lambert et al., 2018; Lambert et al., 2019). The 188 members in the TF families with less TFBM conservation cover 39 of the 76 identified distinct TFBMs (Table 1). In the C2H2 family multiple different domains are annotated compared to the WRKY or Trihelix family, potentially correlating with the number of TFBMs within the family with a Pearson coefficient of 0.69 (see Figure S53). Diverse families have on average four domains annotated while semi-conserved families have 3.4 and conserved families have only about 1.7 domains per family (Figures 2, 3). More diversity of annotated protein domains and variation in repeat number of protein domains might explain part of the TFBM diversity in a TF family. The TF families identified as diverse readily evolve TFBMs *de novo* compared to conserved and semi-conserved families, thus creating possibility for neofunctionalization of TFs.

## Discussion

The analyses presented here indicate that despite having a large portfolio of thousands of sequence-specific TFs, plants like *A. thaliana* operate with a limited vocabulary of TFBMs. More than half of the TF families in *A. thaliana* show a high level of TFBM conservation. In cases of multiple groups of highly similar TFBMs within one family, these groups overlay with phylogenetic clades, suggesting that a difference in protein sequence goes along with changes in TFBMs. Similar findings have been reported in human (Lambert et al., 2018) and yeast (Nitta et al., 2015). As in animals, we observe a number of diverse TF families *in planta* that have evolved multiple TFBMs over time. The C2H2 family was reported as diverse in all studies (Nitta et al., 2015; Lambert et al., 2019; Han et al., 2020), proposing that some families are inherently more diverse across multiple kingdoms of life. Large scale eukaryotic analyses of TF binding characteristics have pointed to amino acid conservation in the protein DBD as a predictor of TFBM conservation (Weirauch et al., 2014). This study proposes a TF family argument for plant TFs: in 21 families, only a single core TFBM has been identified so far, irrespective of the degree of conservation on the protein sequence level (Figure 2). For these TFs, WGD and/or tandem duplications have not expanded the repertoire of TFBMs. Given the presence of *M. polymorpha* orthologs in the subclades of the phylogenetic tree, it can be inferred that the subclades are at least 500 million years old (Harris et al., 2022). Therefore, it can be assumed that members of the same TF family within *A. thaliana* and also within angiosperms likely bind the same TFBM. The TFBMs are shown to be stable on an evolutionary scale from bryophytes to dicotyledons. For additional four families, a high degree of conservation within subclades was observed (Figure 3). The *de novo* evolution of binding sites deep within the phylogenetic branches indicates that, in principle, TF families have the ability to evolve new binding specificity but this is a very rare occurrence. It is likely that uncharacterized TFs bind the known subclade TFBM within the families with high TFBM conservation (Figure 2) and semi-conservation (Figure 3). The third, diverse group of TF families demonstrates a high ability to evolve new and different TFBMs. To expand the currently known vocabulary of 76 TFBMs in *A. thaliana* and other plants, TF families with diverse TFBMs (i.e. Figure 4), especially those with large phylogenetic distance to TFs with known TFBMs, present attractive targets. The expression analyses throughout 6,033 *A. thaliana* wildtype RNA-Seq experiments demonstrate that TFs binding the same TFBM frequently have similar expression patterns (Figures 2C, 3C) indicating that these TFs compete for binding to the TFBM. Therefore, single TFBMs act as integrators of signals in plants where a combination of binding affinity (Jolma et al., 2013; Sielemann et al., 2021) and TF abundance determine the output.

Competition at TFBMs also provides an explanation for the high variation in complementation experiments for TFs on genomic sites, which are bound by many partners (Lee et al., 2007; Stracke et al., 2010; Gangappa and Botto, 2016). The bZIP TF ELONGATED HYPOCOTYL 5 (HY5) and its binding to the partial G-box motif ACGT TFBM is a prime example (Figure 5). This TFBM and its variations can be bound by at least 58 TFs (Figure 5, Table S1) (Ang et al., 1998; Chattopadhyay et al., 1998; Yadav et al., 2002). In all tissues at all times under all conditions where at least a subset of these competing TFs is expressed, they compete with HY5 for TFBM binding. For example, in leaf, 52 competing TFs are expressed with at least >1 transcript per kilobase million (tpm) compared to HY5 at 11 tpm and in the hypocotyl. The same 52 competing TFs are expressed with at least >1 tpm compared to HY5 at 39 tpm (Figure 5). As a consequence, one expects both an effect of HY5 dosage and a strong effect of condition on the outcome of a HY5 binding experiment and/or RNA-Seq experiment. In addition, potential binding at a TFBM will be affected by chromatin openness (Maher et al., 2018; Lu et al., 2019), potential histone modifications (Charron et al., 2010; Zhao et al., 2019), changes in DNA-shape due to binding of other proteins (Rohs et al., 2009) and TFBM flanking regions (Gordân et al., 2013). Indeed, detailed analyses of complementation experiments for HY5 with varying gene doses and varying tags on the protein show the expected large degree of variation in both binding sites and RNA-Seq results (Lee et al., 2007; Stracke et al., 2010; Gangappa and Botto, 2016). In the absence of a competing TF, i.e. in a knock-out of a TF, it is highly likely that at least some of the binding sites previously preferentially occupied by that TF are occupied by closely related TFs. If this occupancy is above a certain threshold it prevents an observable phenotype, and the TFs are called redundant (Finkelstein et al., 2005; Leivar and Monte, 2014). However, it is likely that RNA-Seq analyses of single mutants will uncover subtle shifts in transcript abundances, which do not necessarily lead to observable phenotypes i.e. single *phytochrome interacting factor (pif)* mutants (Leivar et al., 2012). If TFs operate in concert, with their affinities and their abundance also regulating promoter output, one would expect that TFs are preferentially retained after WGDs, even if they do not immediately sub– or neofunctionalize. Such retention has indeed been observed (Van de Peer et al., 2009; De Smet et al., 2013; Schmitz et al., 2016). The large degree of TFBM similarity and the co-occurrence of TFs binding the same TFBM extends the network argument previously made for preferential TF retention (Van de Peer et al., 2009; De Smet et al., 2013; Schmitz et al., 2016) to the output of individual promoters. Similar to the argument of the evolutionary ratchet that preserves interacting partners in a protein complex since mutation of a single one could expose hydrophobic surfaces and throw the system out of equilibrium (Hochberg et al., 2020), TFs integrating on the same binding site require equilibrium for stability after WGD.

The 76 distinct TFBMs present TF binding sites on which transcriptional regulation signals can be integrated. It has long been known that the promoter syntax, i.e. the positioning of binding sites relative to each other and relative to the transcription start site, carries critical information (Arnosti and Kulkarni, 2005). The current analyses show that not only TFs have the same core TFBM, but they also co-occur based on expression studies (Figure 2, 3). This suggests that binding affinity, perhaps modulated by other factors i.e. TF::TF interactions, DNA shape, or histone interactions, and spatial TF abundances likely play a major role in regulation (Figure 5). Indeed, binding affinity is modulated by DNA shape, which in turn is defined by the bases surrounding the TFBM (Rohs et al., 2009; Gordân et al., 2013; Sielemann et al., 2021). These DNA-shape differences explain why a TFBM is preferentially bound by some TF family members and not others (Sielemann et al., 2021). Signal integration at single TFBMs may provide one explanation for the still substantial gap in predictability of gene expression from sequence (Li et al., 2018; Avsec et al., 2021b; de Almeida et al., 2022). Finally, competition for binding at single sites may explain why TFs counterintuitively act as both activators and repressors (Mahendrawada et al., 2023). If a TF activates expression in an enhanceosome context, any competitive binding at its site will turn the different, competitive TF into a repressor for that particular gene irrespective of whether that TF is an activator on other genes. This may explain, why HY5 experiments with the TF carrying an activator or repressor domain still fail to clarify if native HY5 acts as a repressor or activator on its targets (Burko et al., 2020).

Modeling of transcriptional regulation will likely have to go beyond TF binding sites and promoter syntax (Avsec et al., 2021a; Reiter et al., 2023). Modeling transcriptional regulation and predicting gene expression from sequence (DNA and protein) will likely require kinetic parameters, such as K_D_ for TFs with respect to specific TFBMs and amounts of TFs to account for signal integration at TFBMs bound by multiple, frequently co-occuring TFs.

## Methods

### Amplified DNA affinity purification sequencing (ampDAP-Seq)

Plants were grown for six weeks on half-strength Gamborg’s medium (Gamborg B5; Duchefa Biochemie B.V., Netherlands) in petri dishes with a 16h/8h light/dark cycle at room temperature. DNA was extracted form male G2 generation *M. polymorpha* subsp. *ruderalis* BoGa (Busch et al., 2019) with the cetyltrimethylammonium bromide (CTAB) method (https://dx.doi.org/10.17504/protocols.io.bcvyiw7w).

DNA (5 µg) was fragmented by sonication to 200 bp with the M220 Focused-Ultrasonicator (Covaris, USA). End-repair, A-tailing and Y-adaptor ligation were performed following the protocol of (Bartlett et al., 2017): For the sample clean-up the DNA was purified using AMPure XP beads (Beckman Coulter, USA) instead of ethanol precipitation. To obtain an ampDAP library 15 ng of the DAP library was amplified with 11 cylces PCR. For the binding assay, genes were cloned in pFN19A (HaloTag®7) T7 SP6 Flexi® vector (Promega, USA) by Gibson assembly (Gibson et al., 2009). Halo-tagged TFs were expressed with TnT® Coupled Wheat Germ Extract System (Promega, USA) using 2 µg plasmid DNA. Halo-fusion proteins were purified with Magne® HaloTag® Beads (Promega, USA) and then incubated with 50 ng ampDAP library. DNA was recovered, amplified and indexed with 20 PCR cycles. Large fragments (200-400 bp) were extracted from a 1% agarose gel with QIAquick Gel Extraction Kit (Qiagen, Netherlands). The final library was sequenced as 85 bp long single-end reads on a NextSeq™ 550 (Illumina, USA).

### TFBM acquisition

TF families and their members in *A. thaliana* were retrieved from TAPscan (Lang et al., 2010; Wilhelmsson et al., 2017). DAP– and ampDAP-Seq data from O’Malley et al. (2016) was obtained from the Gene Expression Omnibus (GEO) database (Barrett et al., 2013) under the accession <u>GSE60143</u> and analyzed according to Sielemann et al., (2021): peak sequences were extracted from the TAIR10 reference genome (https://www.arabidopsis.org/) of *A. thaliana* and TFBMs were determined using MEME-ChIP (Machanick and Bailey, 2011). The TFBM with the lowest e-value and less than 21 bases was chosen to avoid long artifacts. AmpDAP-Seq data from *M. polymorpha* was mapped to the reference genome Tak v6.1 (https://marchantia.info/) using Bowtie v. 2.4.2 (Langmead and Salzberg, 2012). Output files were converted to BAM format with SAMtools (Danecek et al., 2021) view and sorted (samtools sort-n). To remove duplicates, the SAMtools commands fixmate, sort and markdup were executed with default parameters. Peaks were called using GEM (Guo et al., 2012) or MACS2 (Zhang et al., 2008) only if GEM could not identify TFBMs. Sequences of 500 bp around the peak summit were submitted to MEME-ChIP for TFBM extraction. Additionally, we accessed TFBMs directly from the open access databases JASPAR (Castro-Mondragon et al., 2022) and PlantTFDB (Jin et al., 2017).

### Family allocation

All TFs were assigned to TAPscan TF families in *A. thaliana*. If a corresponding TF family in another database exists in TAPscan, annotations were converted to TAPscan names. Accurate allocation was validated via annotated Interpro domains (Blum et al., 2021). TFBMs from other species were assigned to the TF families using blastp (Altschul et al., 1990) with the corresponding protein sequence against the *A. thaliana* TAIR10 proteome (https://www.arabidopsis.org/). The protein sequences of the TFs were retrieved from Uniprot or the respective proteome from Phytozome (Goodstein et al., 2012) if available. The best blast hit ranked by e-value and secondly percentage of identity was chosen to determine the nearest *A. thaliana* ortholog, whose TF family annotation was then transferred. The MYB family was manually reduced to contain only MYB3R and R2R3-MYB factors in accordance with annotations from Table 1 in Stracke et al., (2001) and Plant Cistrome database annotations. Other MYB-type TFs were grouped as MYB-related.

Some TFs had multiple TFBMs generated by different methods, requiring the selection of one representative TFBM per protein per species. We preferred ampDAP-Seq data over DAP-Seq data followed by other methods, because DAP-Seq is a high throughput method capturing the most complete set of binding sites on gDNA (O’Malley et al., 2016) and ampDAP additionally requires less input material and binding is solely based on sequence and not influenced by methylation state. We reassigned the JASPAR TFBMs originally retrieved from ReMap to the original method of either ChIP-Seq or DAP-Seq.

### Phylogenetic analyses

Full protein sequences were aligned per family using MUSCLE v3.8.31. Phylogenetic trees were generated with RAxML v7.4.4 with 1,000 bootstraps and the PROTGAMMAJTT matrix (method modified from Guedes Corrêa et al., 2008) for similarity measure. The phylogenetic trees with TFBMs were visualized using the R package motifStack (Ou et al., 2018) (see Figures S1-S29). Protein sequences were scanned for domain annotations from Pfam and Prosite using InterProScan (Jones et al., 2014).

The similarity of TFBMs within a family was assessed using compare_motifs from the R package universalmotif with default parameters. Clusters were generated by cutting a dendrogram resulting from the distances of the TFBMs at 0.5 followed by manual curation (method modified from Jores et al., 2021). Since gapped TFBMs are more difficult to detect (Bailey et al., 2009) we considered sufficiently similar parts of a gapped TFBM as the same cluster. Consensus TFBMs were generated with mergeMotifs (motifStack) and for gapped TFBMs merge_motifs (universalmotif) for each cluster and trimmed with trim_motifs (universalmotif). We defined TFBMs appearing only once in a family as outliers and excluded them from consensus TFBM generation (method modified from Jores et al., 2021). Distinct TFBMs across all TF families were established by TFBM comparison as described before and the clusters generated from an hclust analysis.

We considered TF families with at least four TFs with a known TFBM for our assessment of conservation level within the family. Families with one consensus TFBM, subtracting outliers, and less than 15% outlier TFBMs were considered as a conserved family. Up to four different consensus TFBMs and less than 15% outliers classified families as semi-conserved (see Table S1).

Orthologues proteins were determined using OrthoFinder2 (Emms and Kelly, 2019) for the TFs with a TFBM in a given phylogenetic clade of the tree.

### Expression data

Wild-type RNA-Seq experiments of *A. thaliana* were downloaded from the SRA (Leinonen et al., 2011) and mapped onto the TAIR10 reference genome (as described in Halpape & Wulf et al., 2023). Pearson correlation was calculated within conserved families and within groups binding to the same TFBM in semi-conserved families and visualized using the corrplot R package. Expression levels in the leaf and hypocotyl of *A. thaliana* from Klepikova et al., (2016) were averaged across the two replicates and log2 transformed.

## Data availability

Raw ampDAP-Seq data for *M. polymorpha* subsp. *ruderalis* BoGa is available under the Bioproject <u>PRJNA1007631</u> on the NCBI SRA. All *A. thaliana* (amp)DAP-Seq binding sites on nuclear encoded gene promoters are deposited under https://doi.org/10.4119/unibi/2982196. Code to analyze binding motifs and the expression of TFs binding a specific motif and all binding motifs in MEME-format are available from GitHub (https://github.com/sanjaze/meta-analysis_TFBMs).

## Supporting information

allSupplementalFigures

allSupplementalTables

## Acknowledgements

This study was funded by the German Research Foundation (Deutsche Forschungsgemeinschaft; DFG) through the Sonderforschungsbereich 175 (SFB-TR175): “The Green Hub, Central Coordinator of Acclimation in Plants”, and through “Evolutionary network analysis based on the transcriptome atlas of *Marchantia polymorpha*” FundingID: BR4617/1-1. We gratefully acknowledge support by the BMBF-funded de.NBI Cloud within the German Network for Bioinformatics Infrastructure, and the CeBiTec compute cluster for computational resources.

