## Supplementary material for "Transcription factors operate on a limited vocabulary of binding motifs in *Arabidopsis thaliana*": allSupplementalFigures

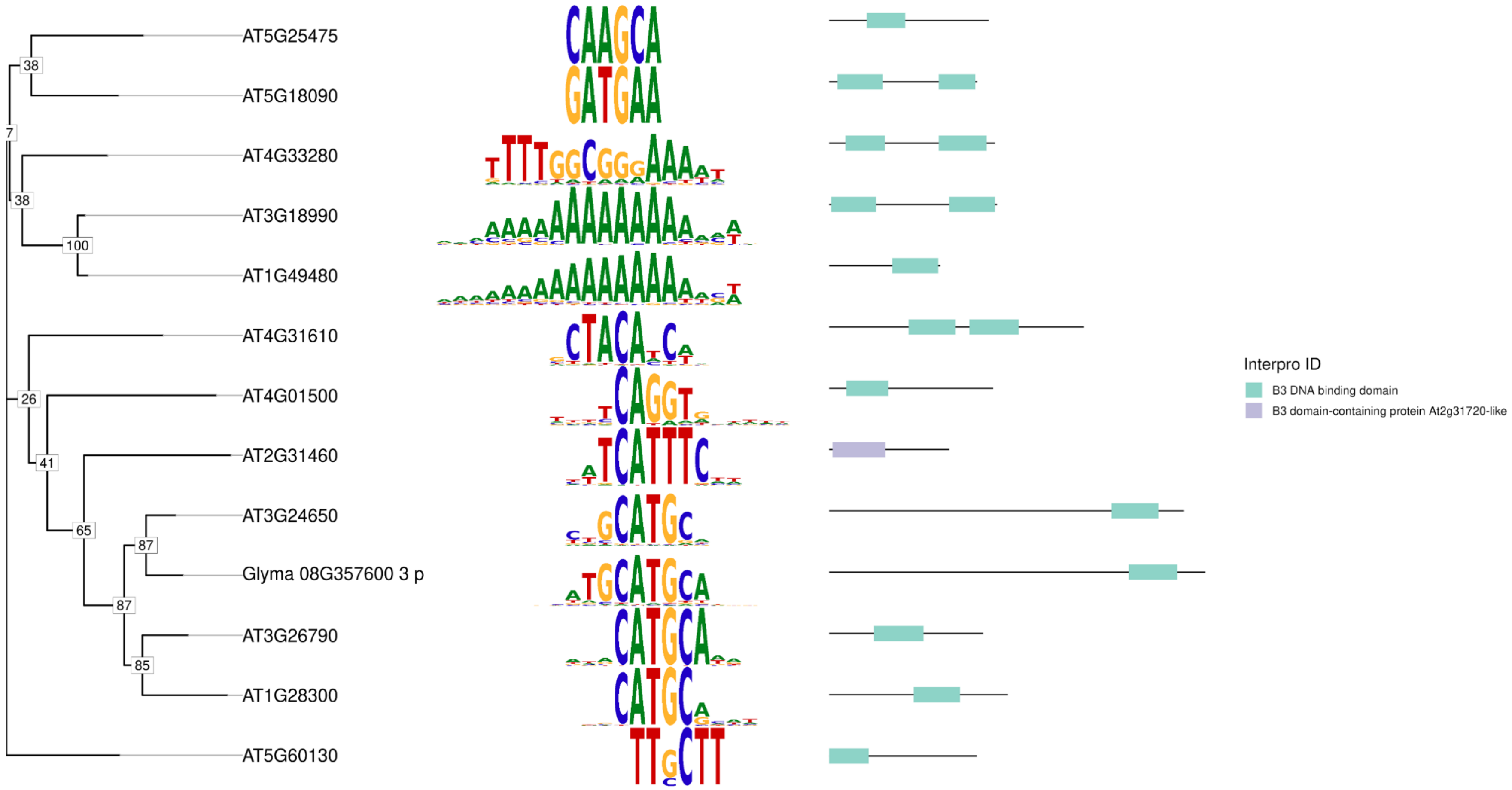


**Figure S1 Phylogenetic tree of the ABI3/VP1 family TFs with TFBMs**

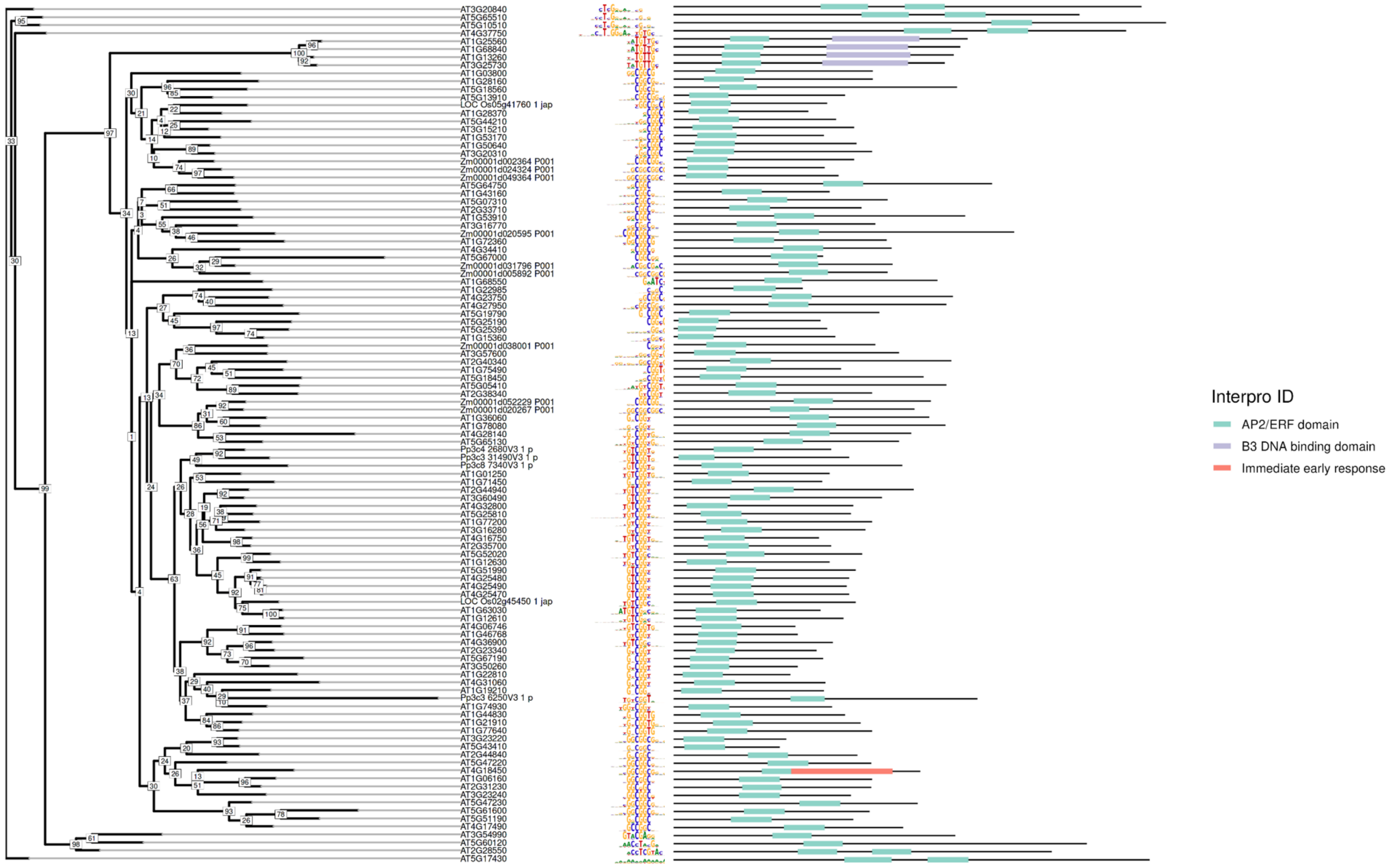


**Figure S2 Phylogenetic tree of the AP2/EREBP family TFs with TFBMs**

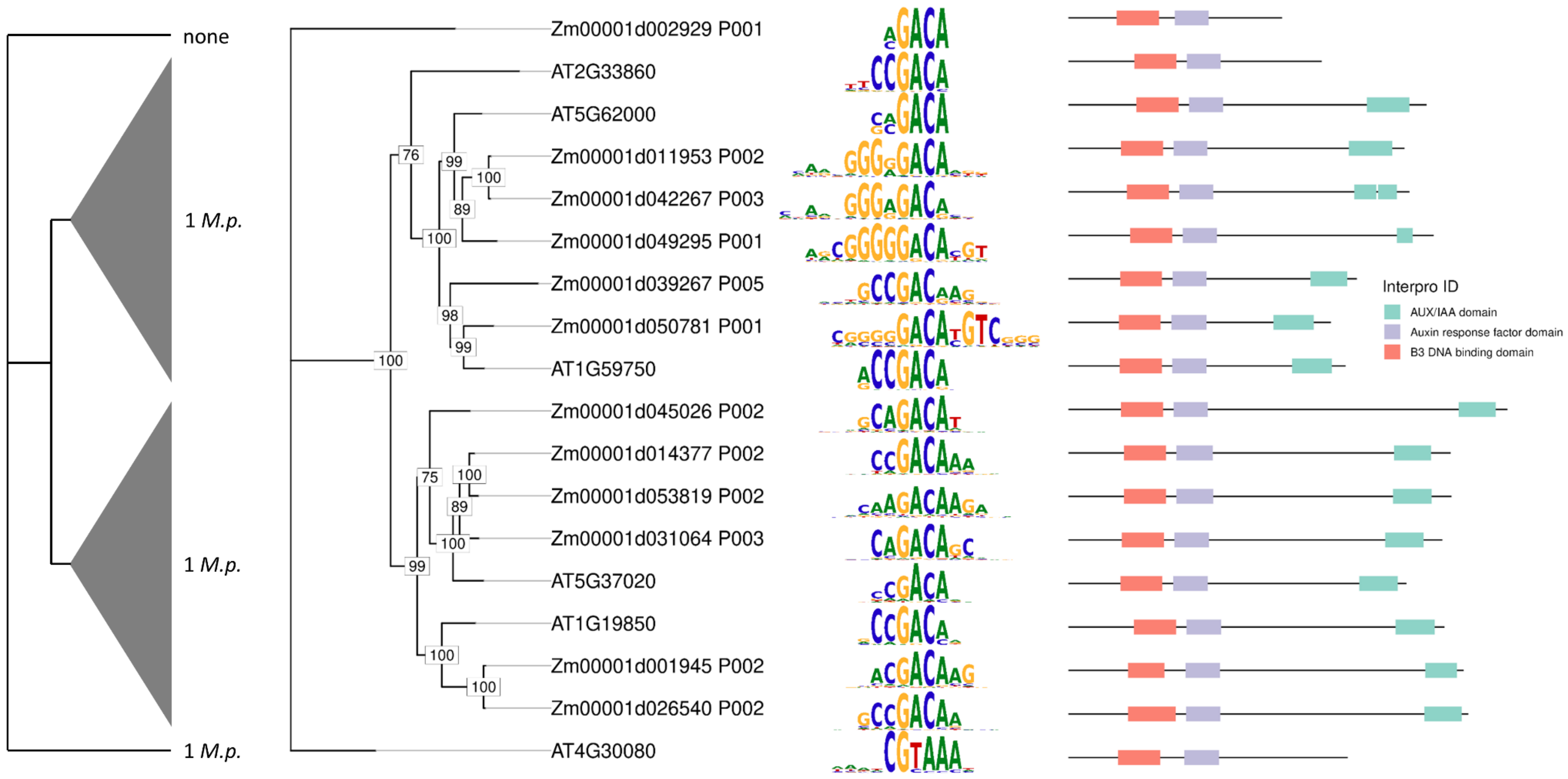


**Figure S3 Phylogenetic tree of the auxin response factor (ARF) family TFs with TFBMs**

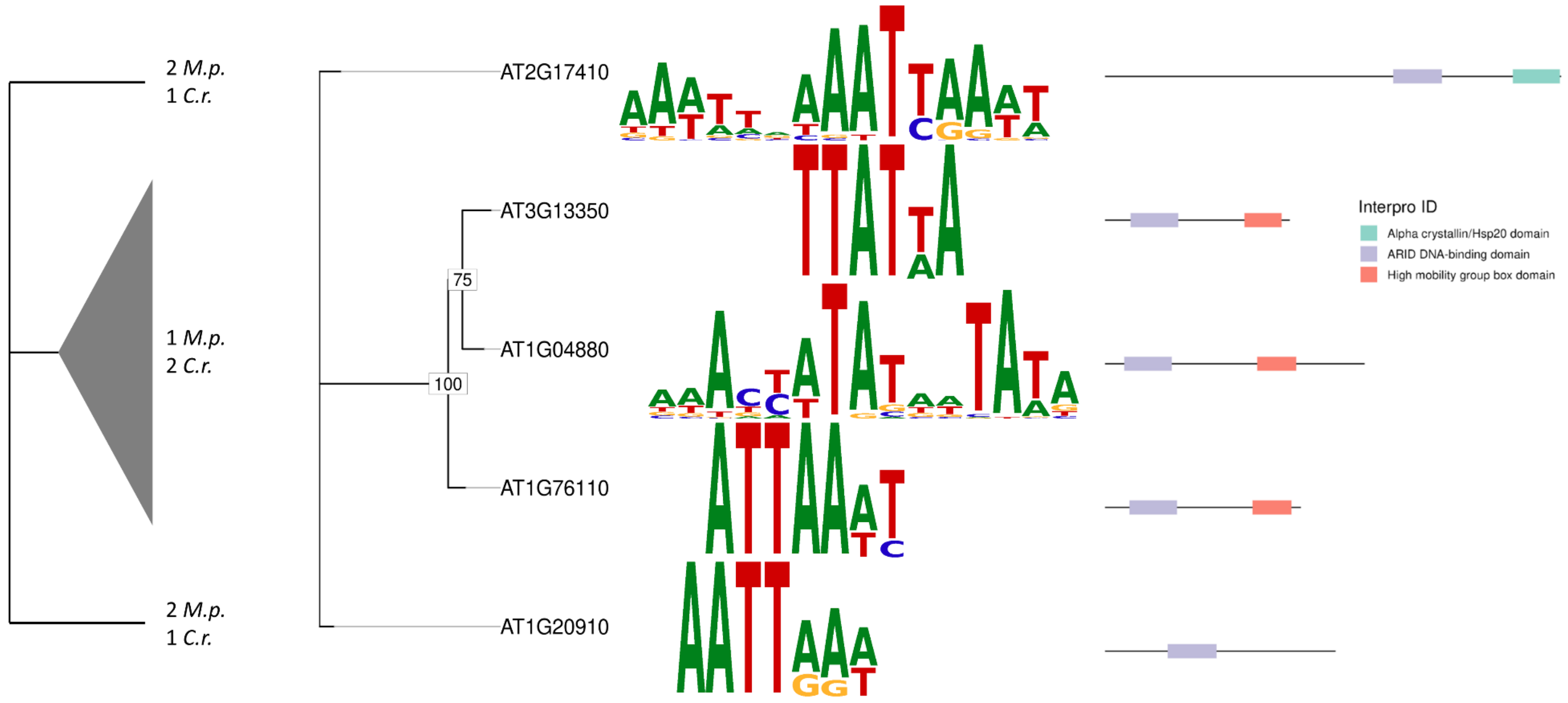


**Figure S4 Phylogenetic tree of the ARID family TFs with TFBMs**

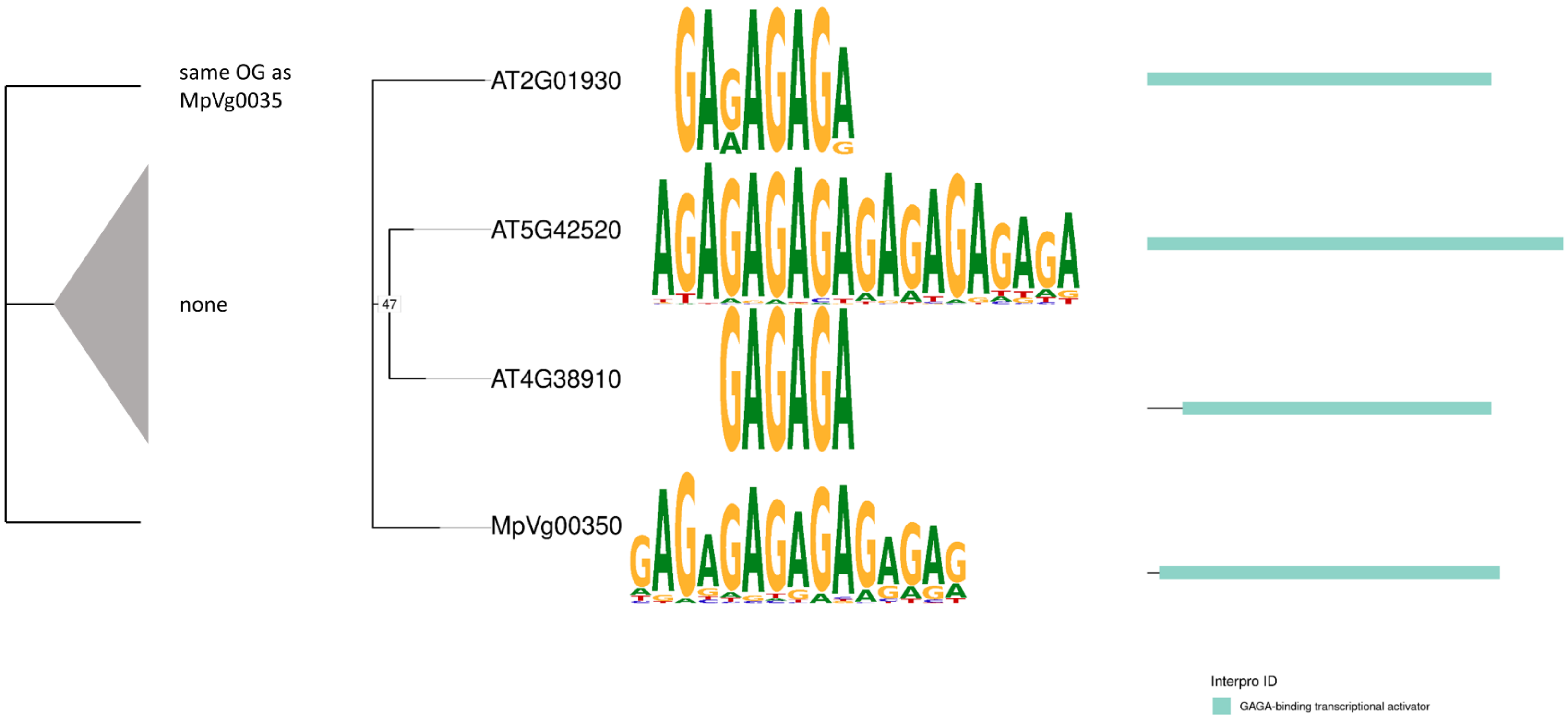


**Figure S5 Phylogenetic tree of the BBR/BPC family TFs with TFBMs**

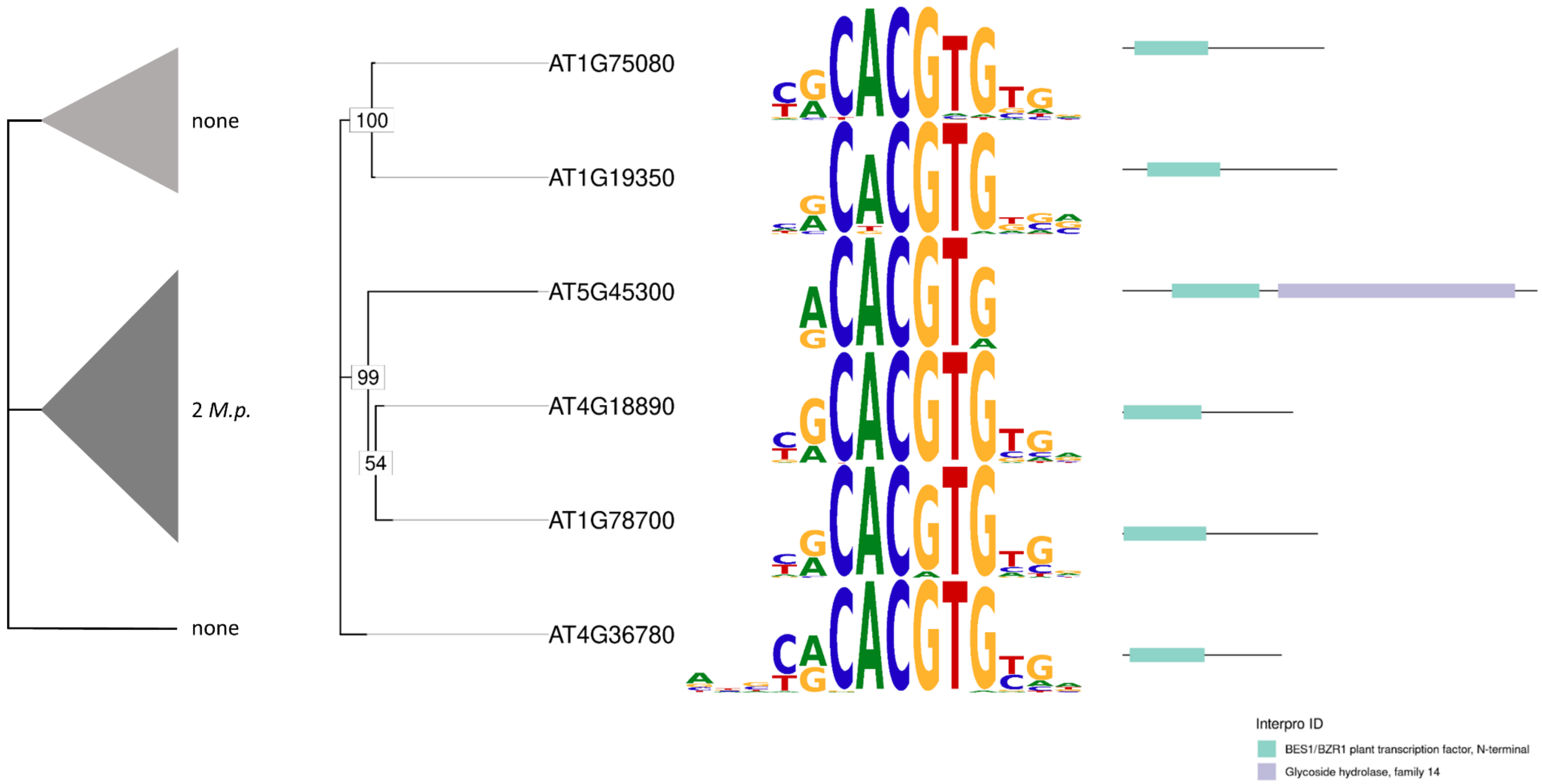


**Figure S6 Phylogenetic tree of the BES1 family TFs with TFBMs**

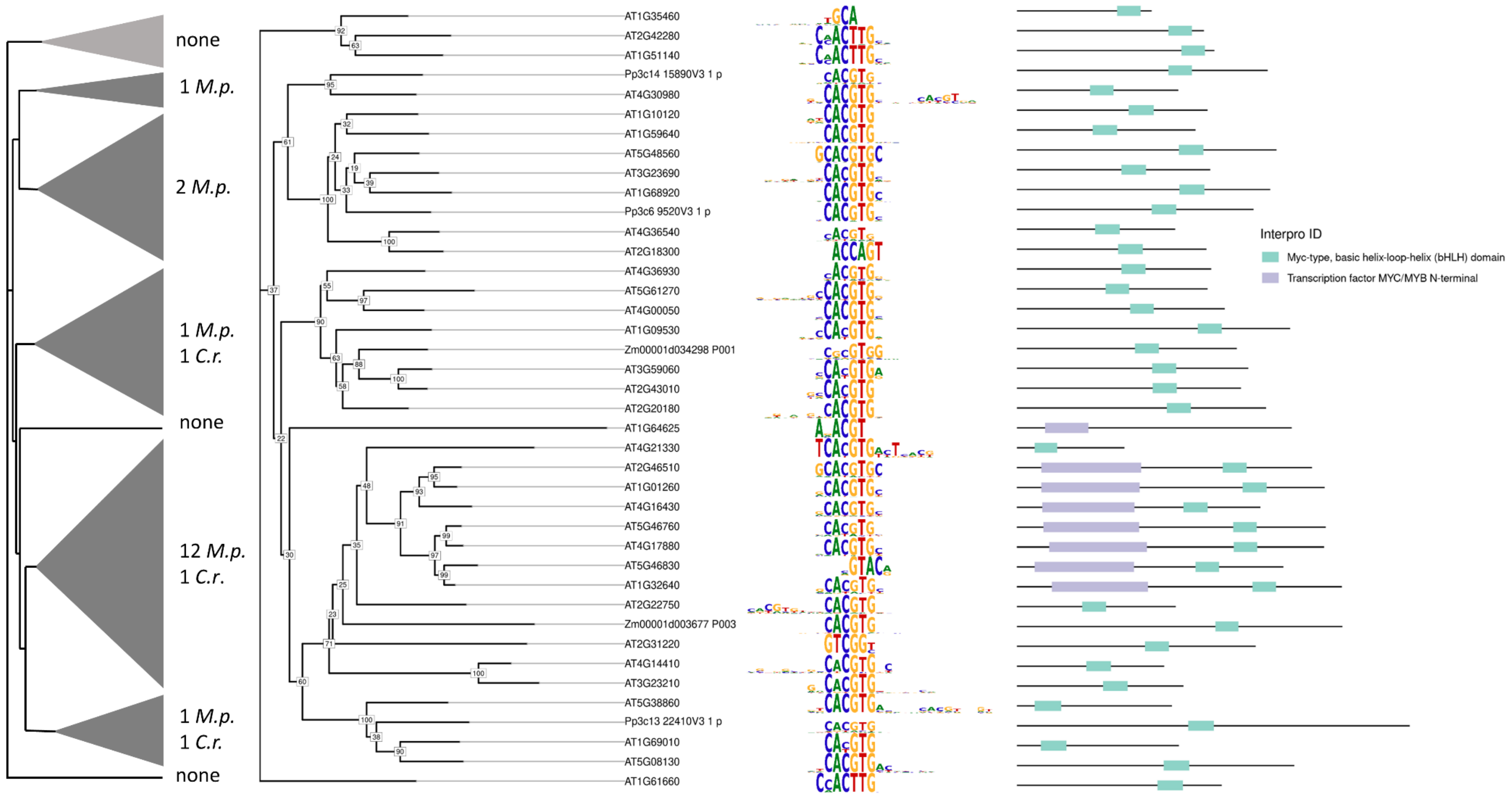


**Figure S7 Phylogenetic tree of the bHLH family TFs with TFBMs**

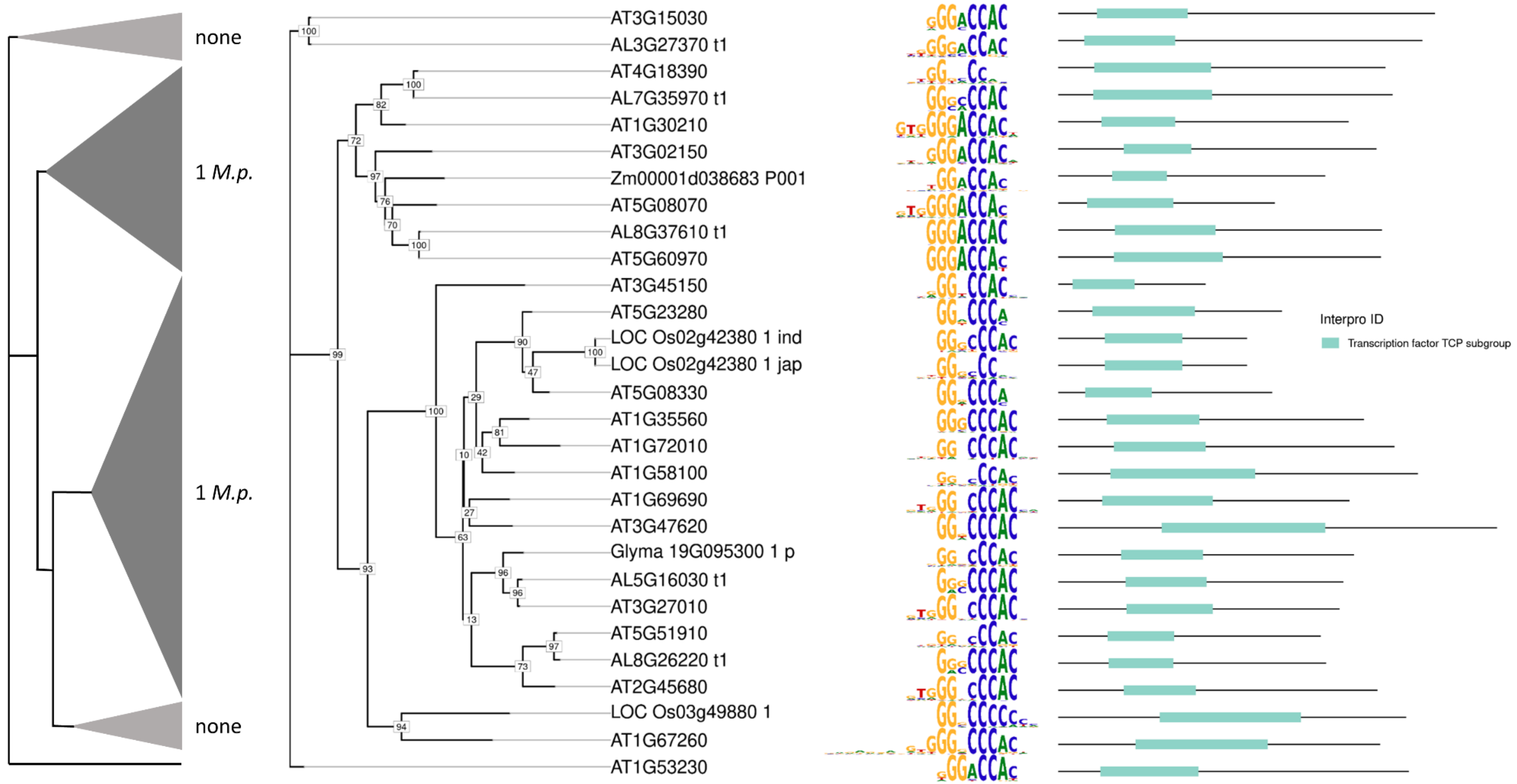


**Figure S8 Phylogenetic tree of the bHLH TCP family TFs with TFBMs**

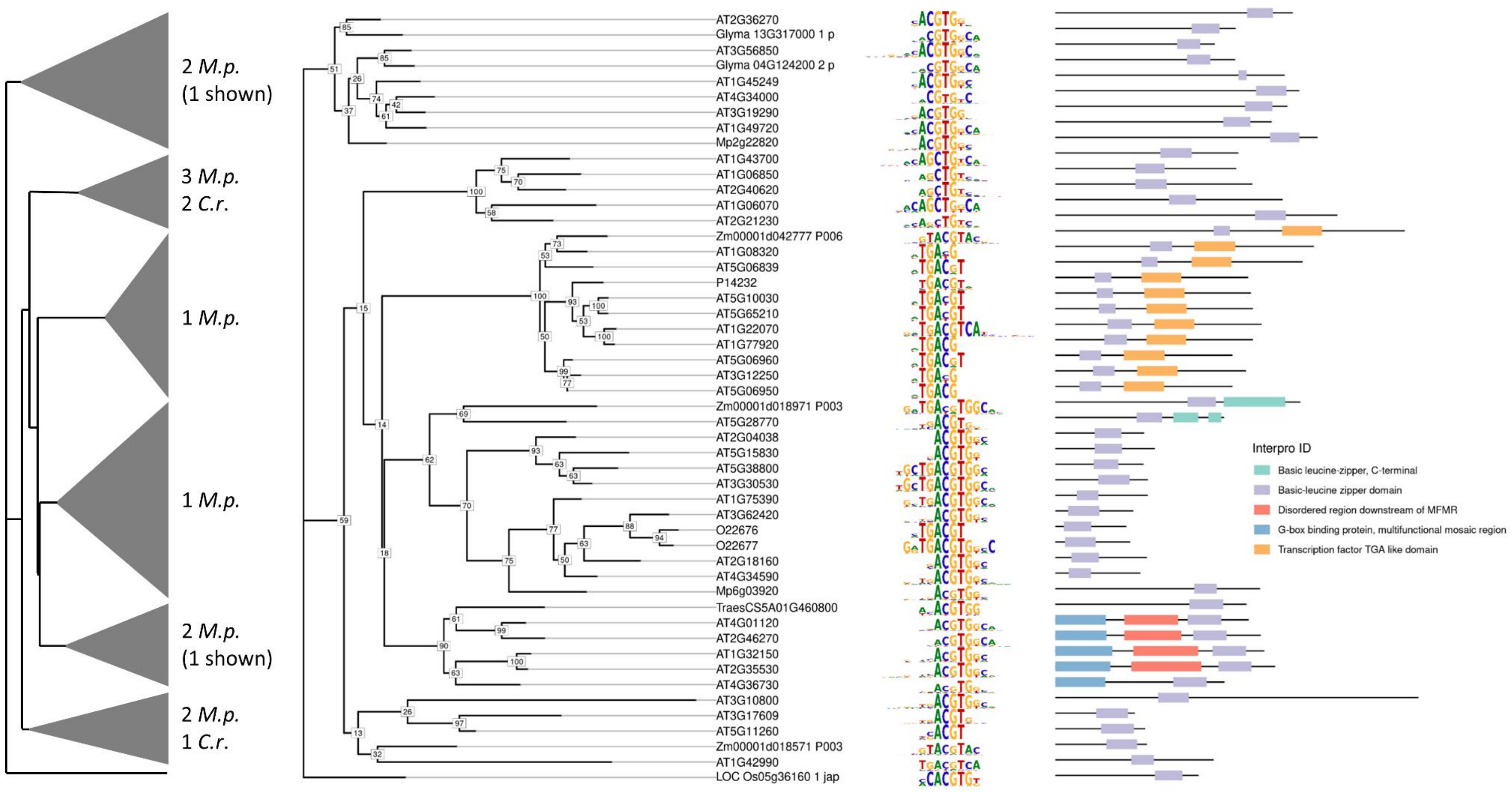


**Figure S9 Phylogenetic tree of the bZIP family TFs with TFBMs**

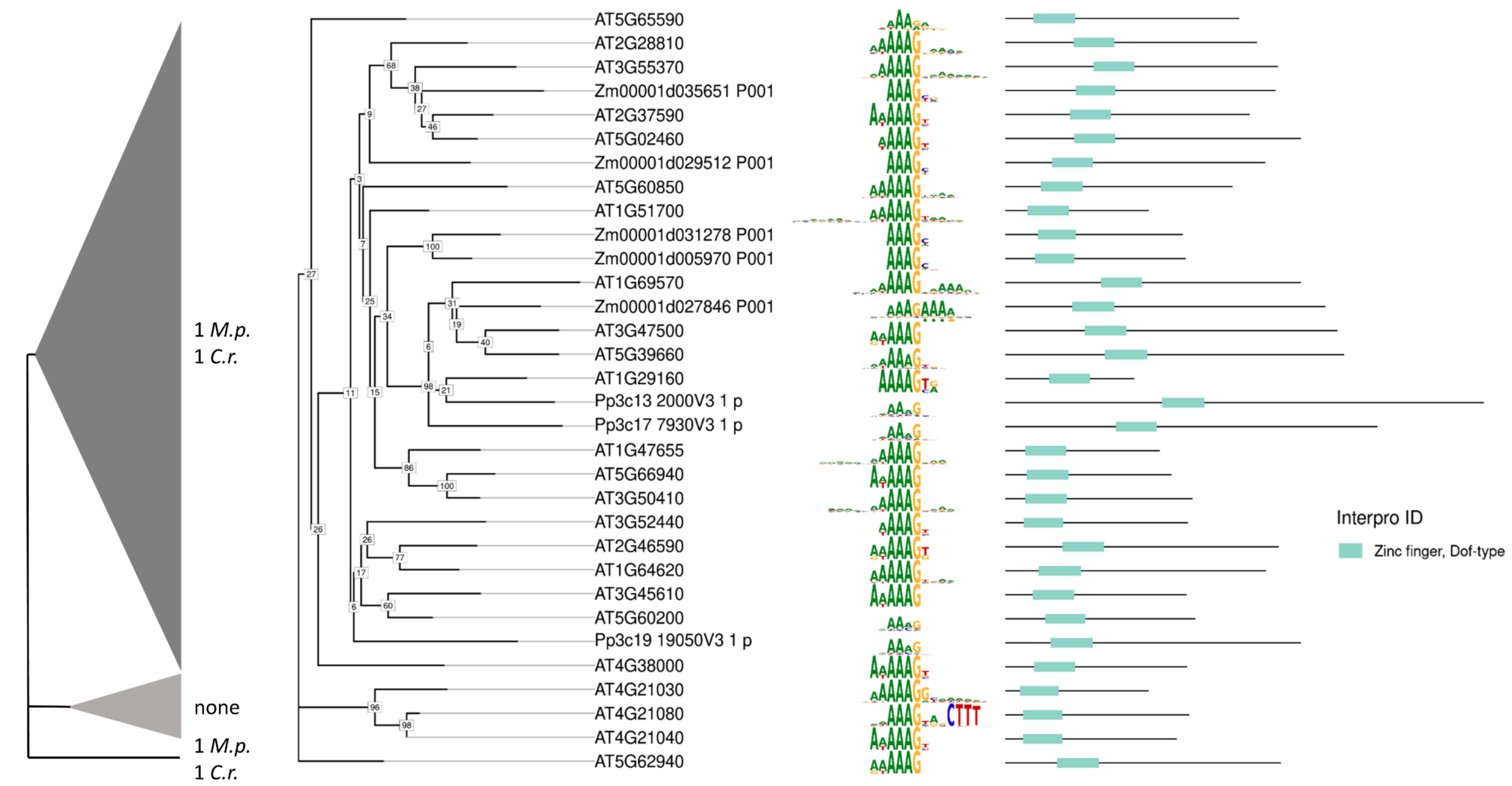


**Figure S10 Phylogenetic tree of C2C2 Dof family TFs with TFBMs**

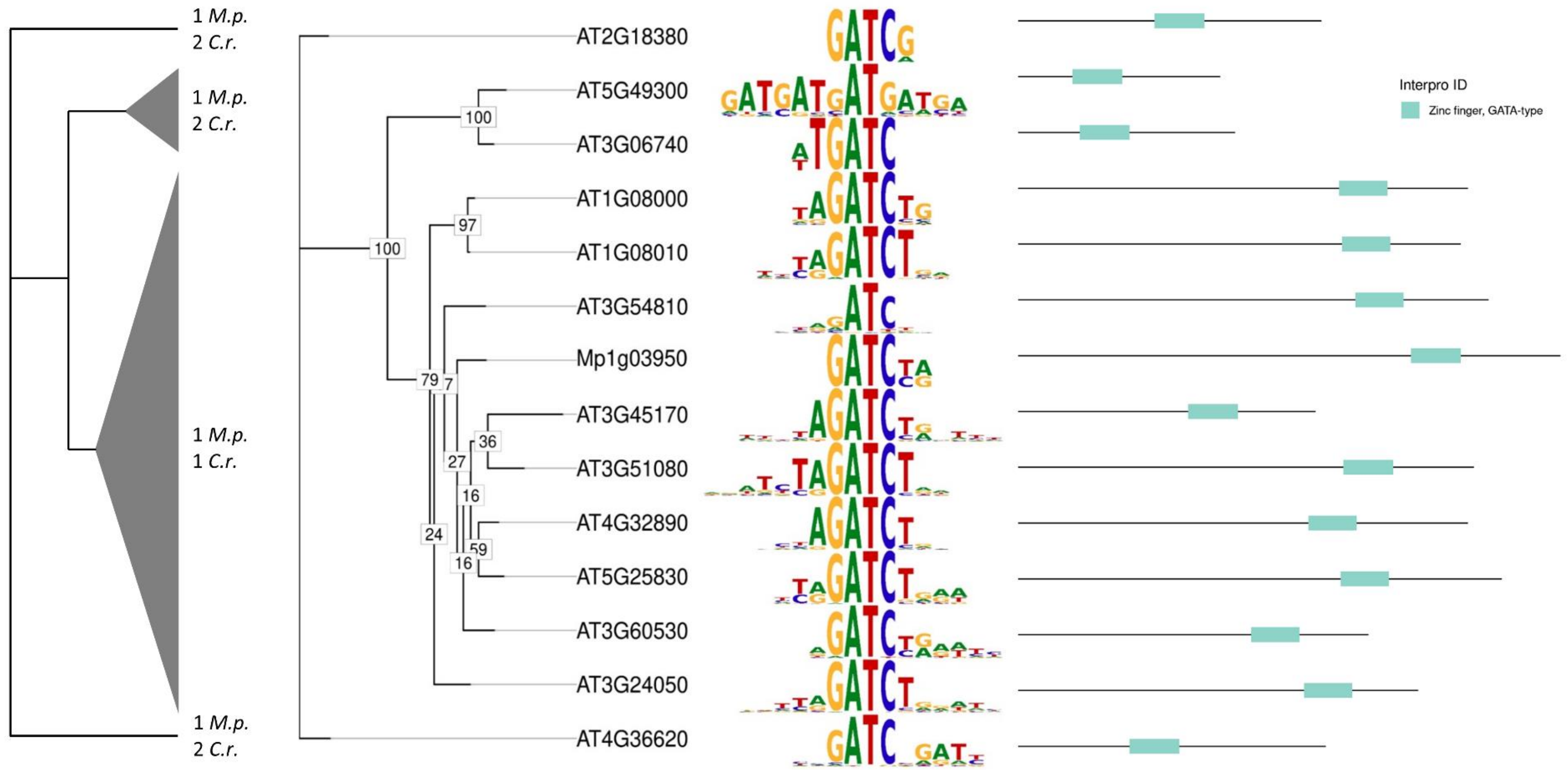


**Figure S11 Phylogenetic tree of the C2C2 GATA family TFs with TFBMs**

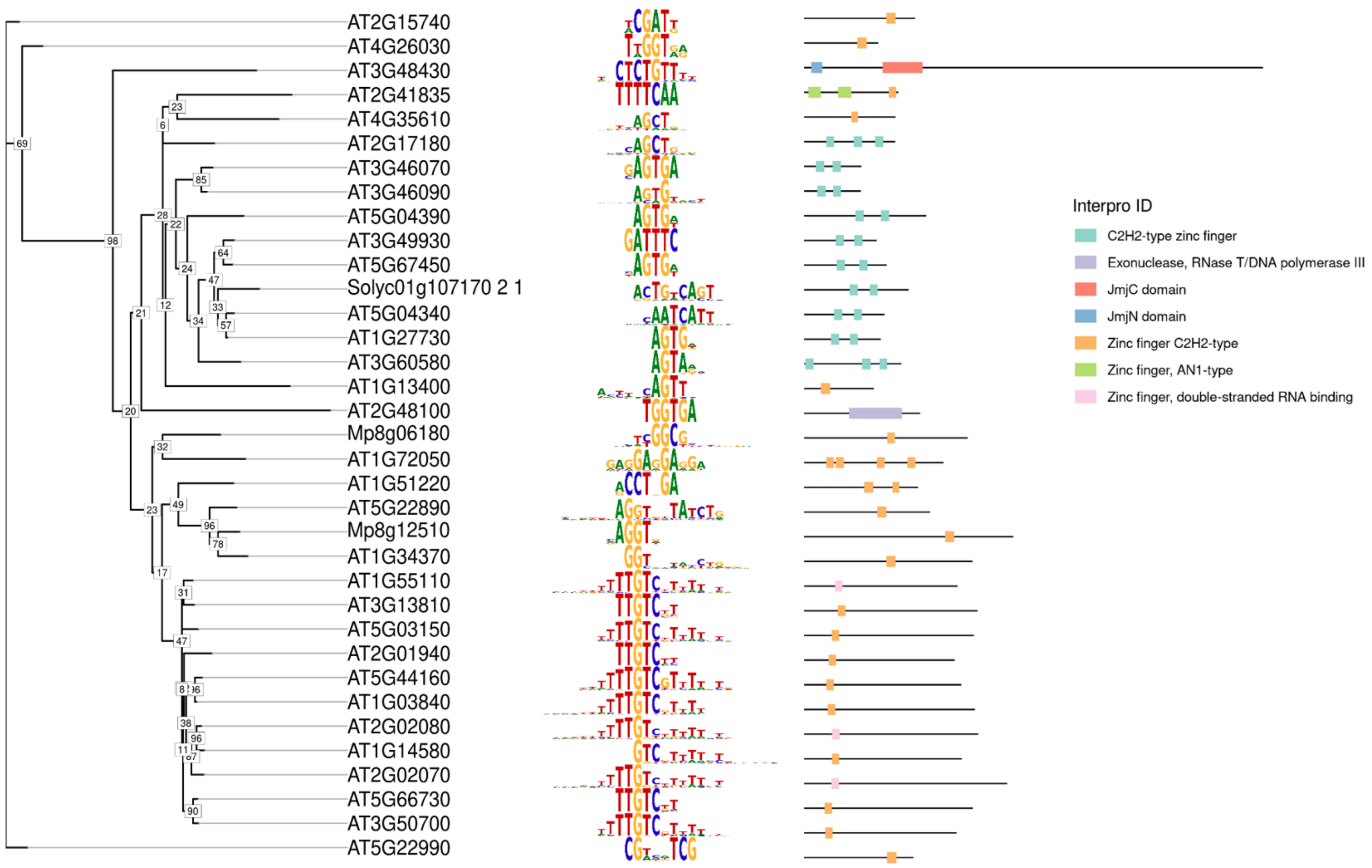


**Figure S12 Phylogenetic tree of the C2H2 family TFs with TFBMs**

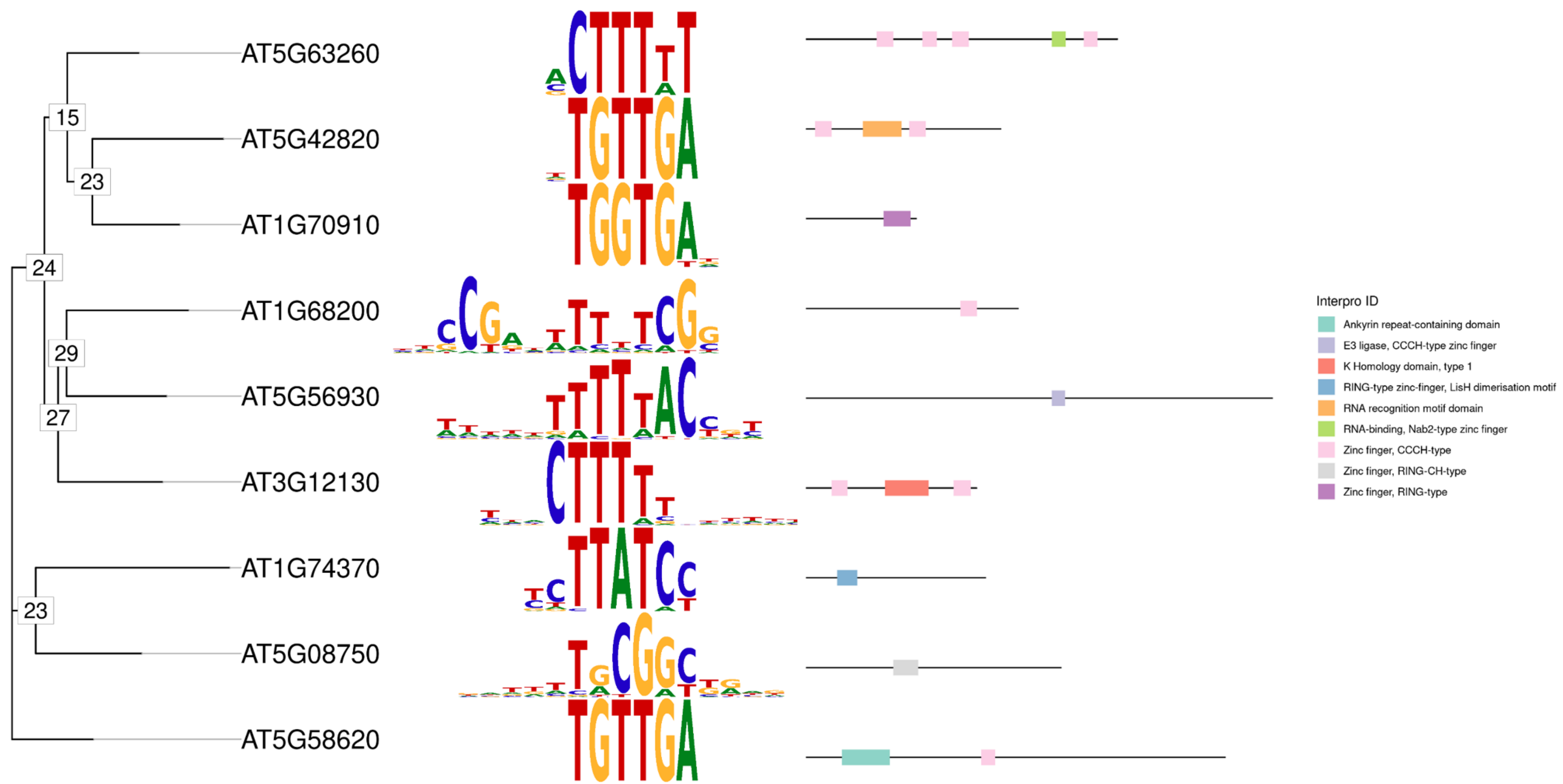


**Figure S13 Phylogenetic tree of the C3H family TFs with TFBMs**

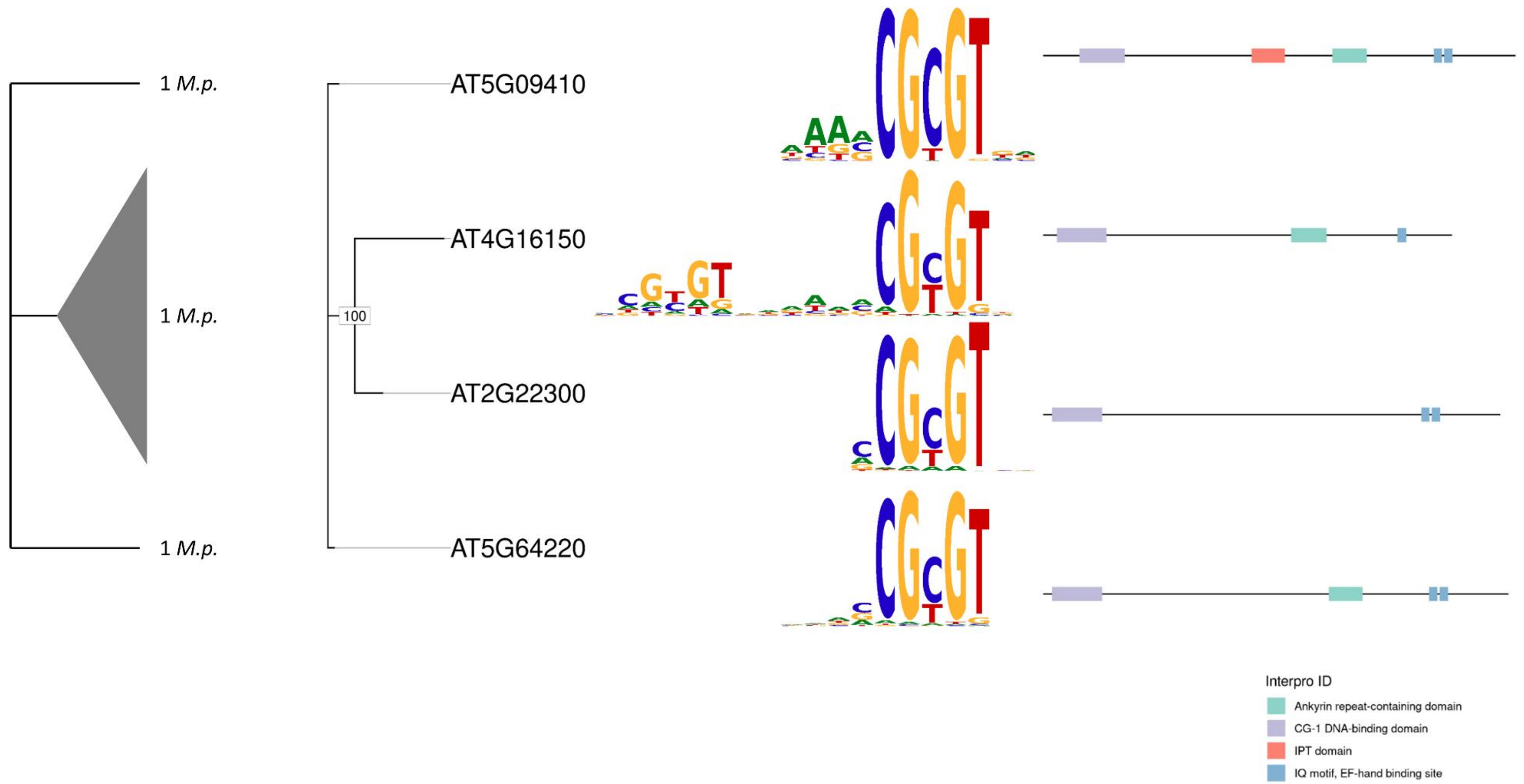


**Figure S14 Phylogenetic tree of the CAMTA family TFs with TFBMs**

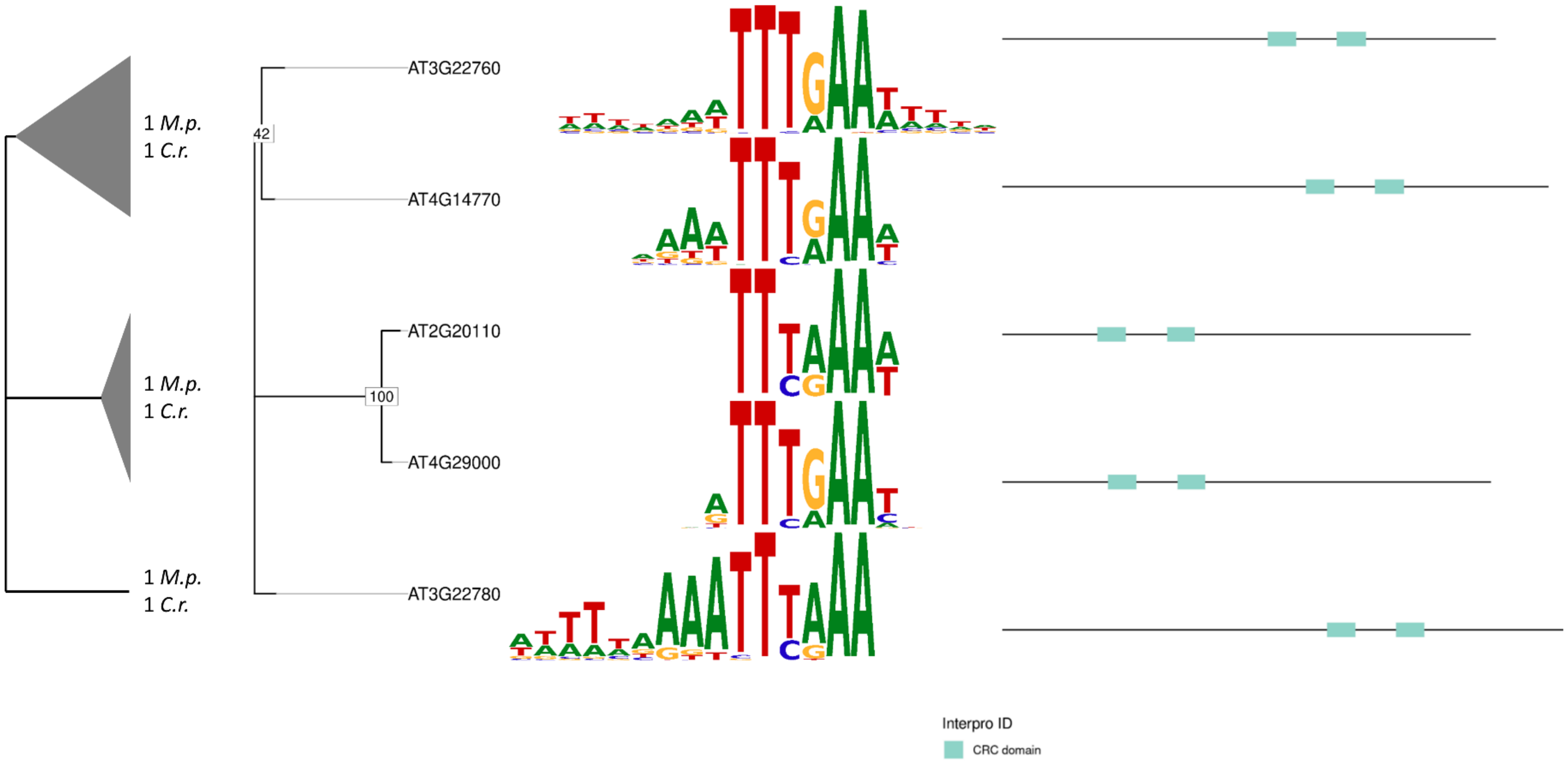


29

30 **Figure S15 Phylogenetic tree of the CPP family TFs with TFBMs**

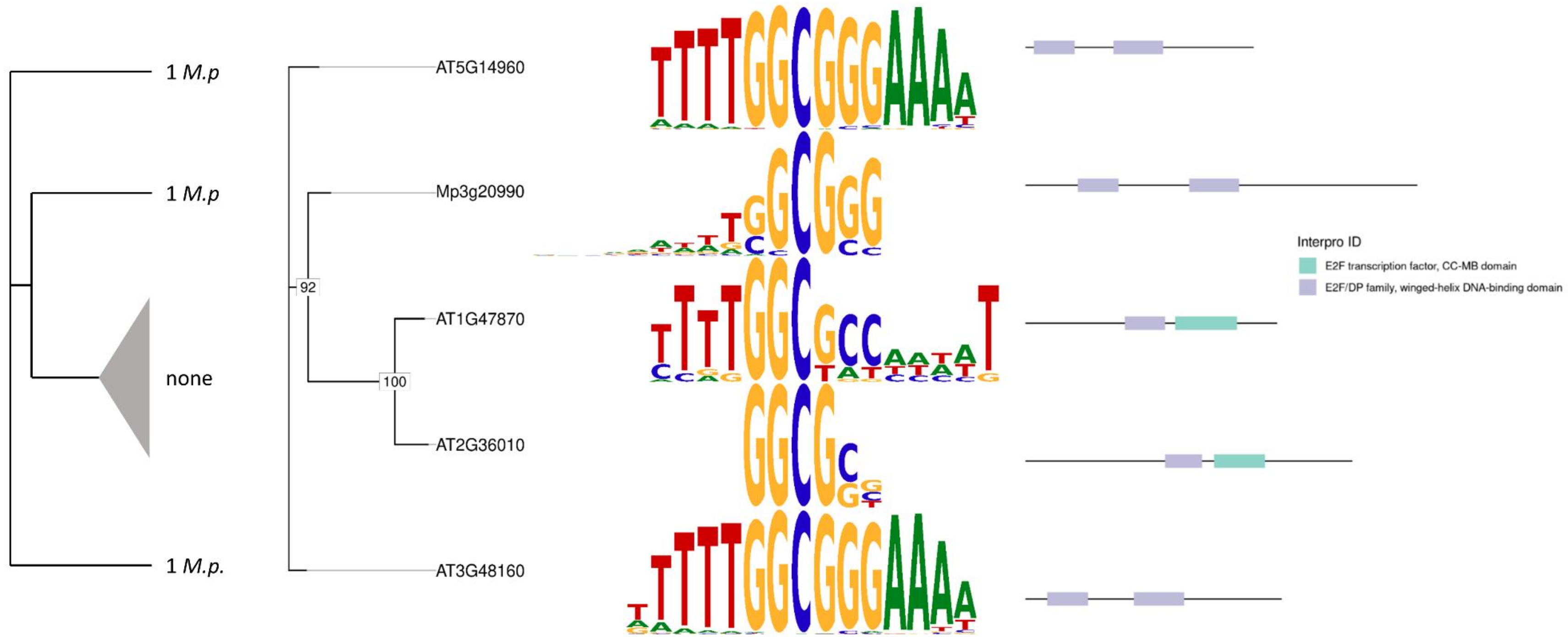


**Figure S16 Phylogenetic tree of the E2F/DP family TFs with TFBMs**

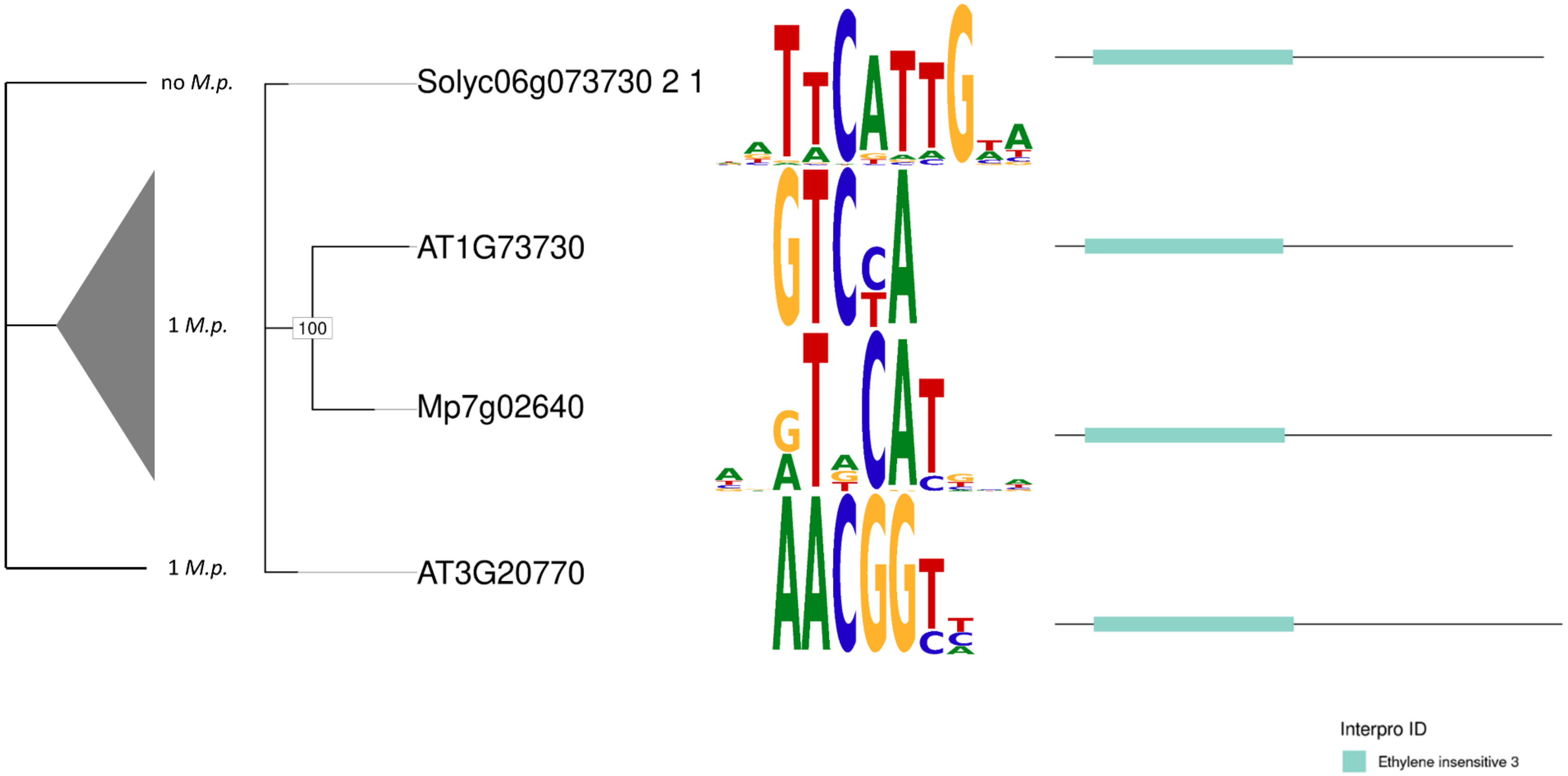


**Figure S17 Phylogenetic tree of the EIL family TFs with TFBMs**

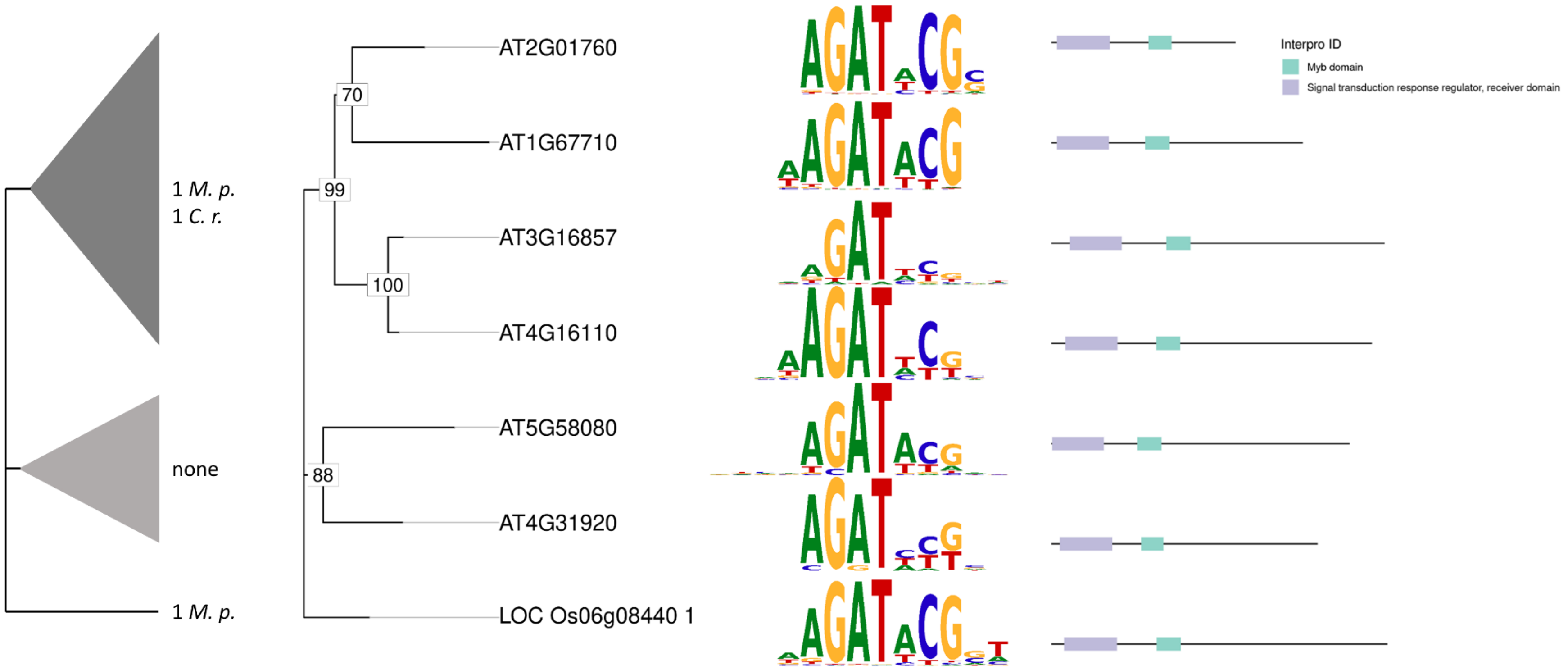


**Figure S18 Phylogenetic tree of the GARP ARR-B family TFs with TFBMs**

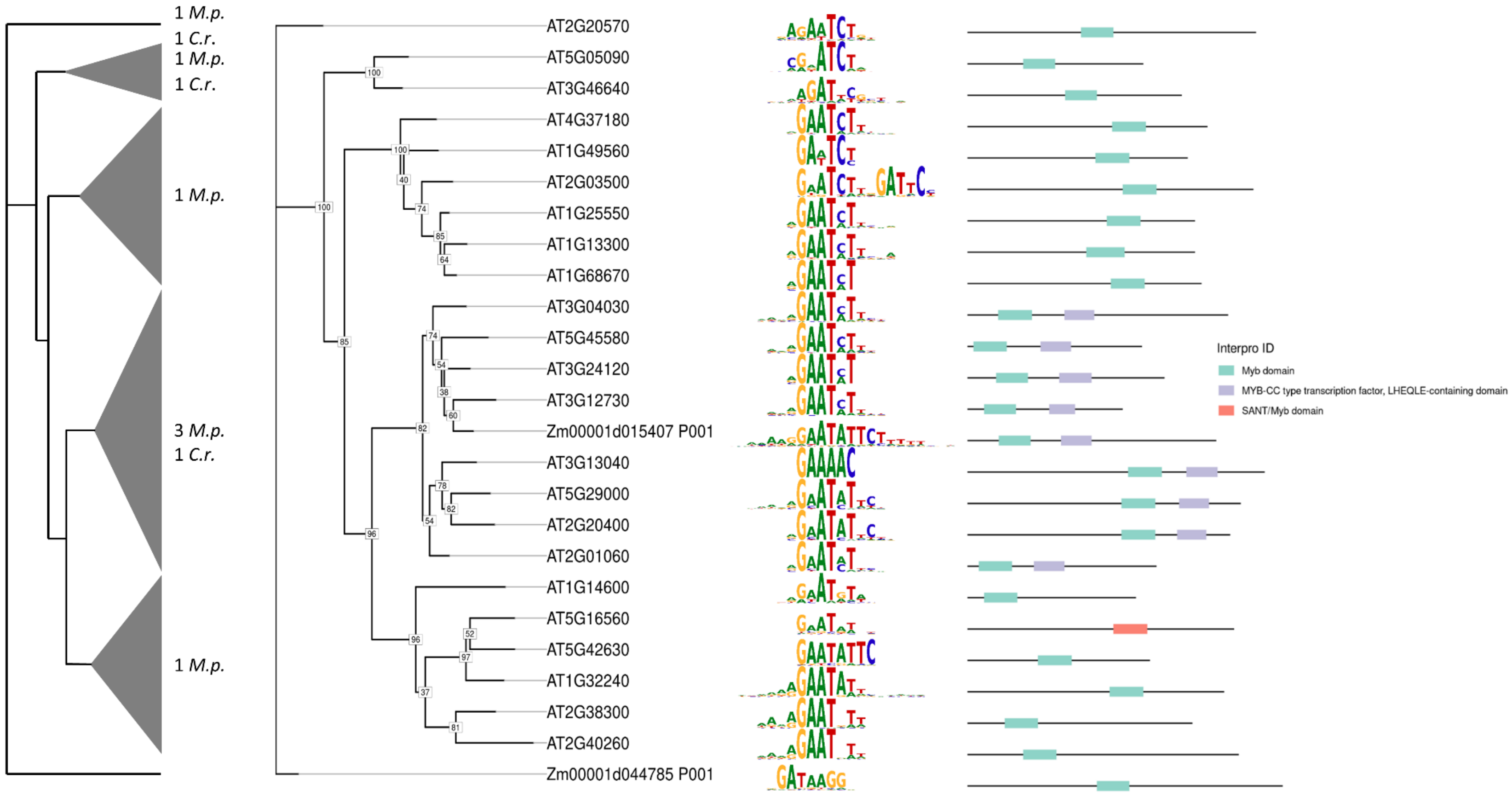


**Figure S19 Phylogenetic tree of the GARP G2-like family TFs with TFBMs**

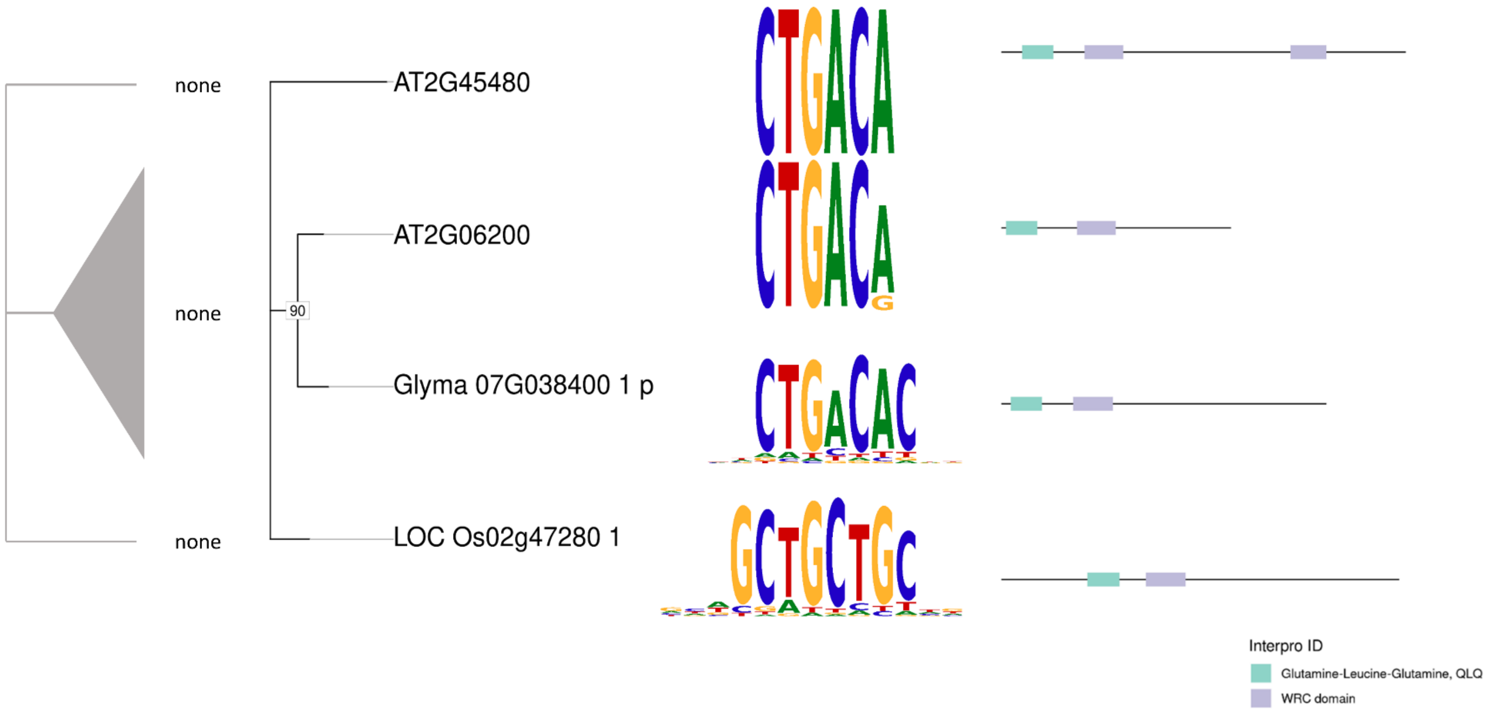


**Figure S20 Phylogenetic tree of the GRF family TFs with TFBMs**

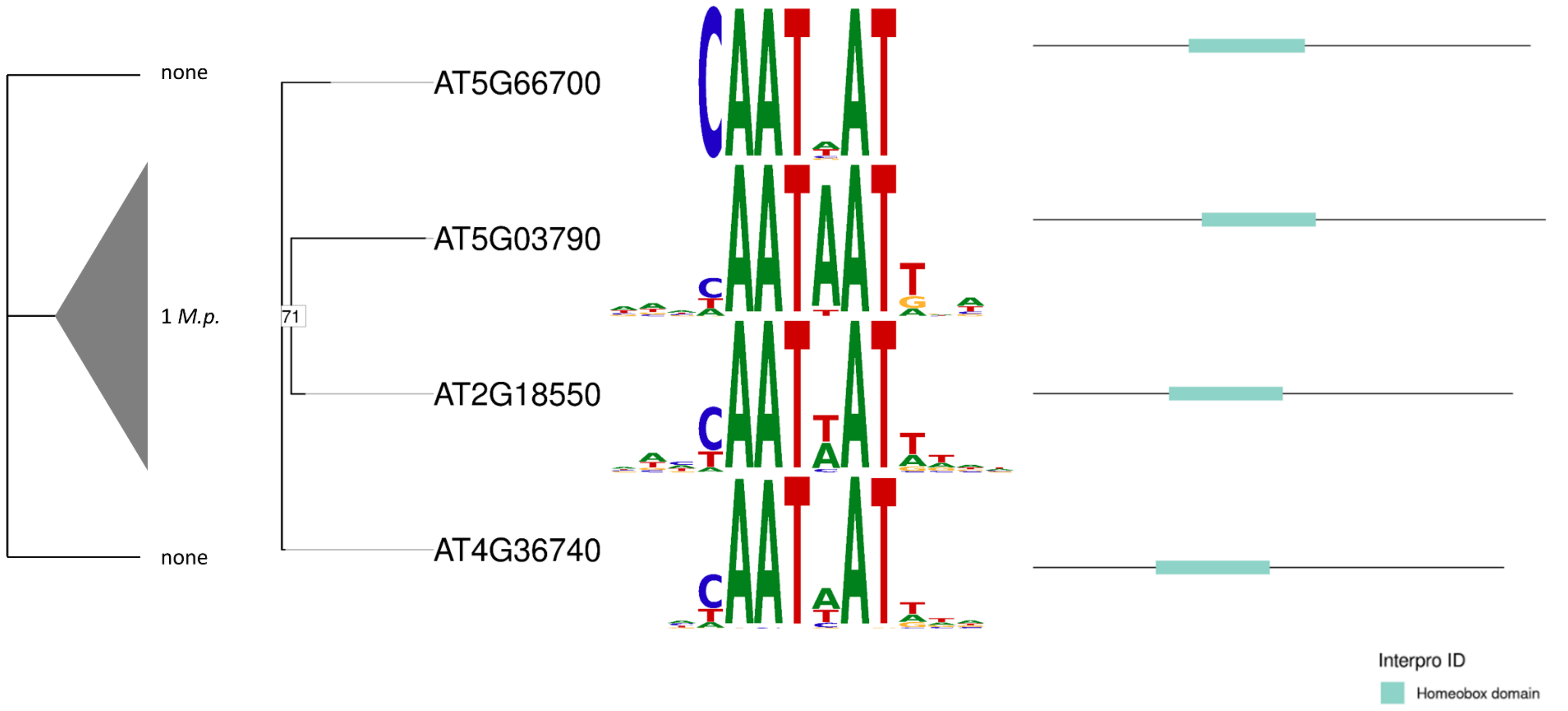


**Figure S21 Phylogenetic tree of the homeodomain (HD) family TFs with TFBMs**

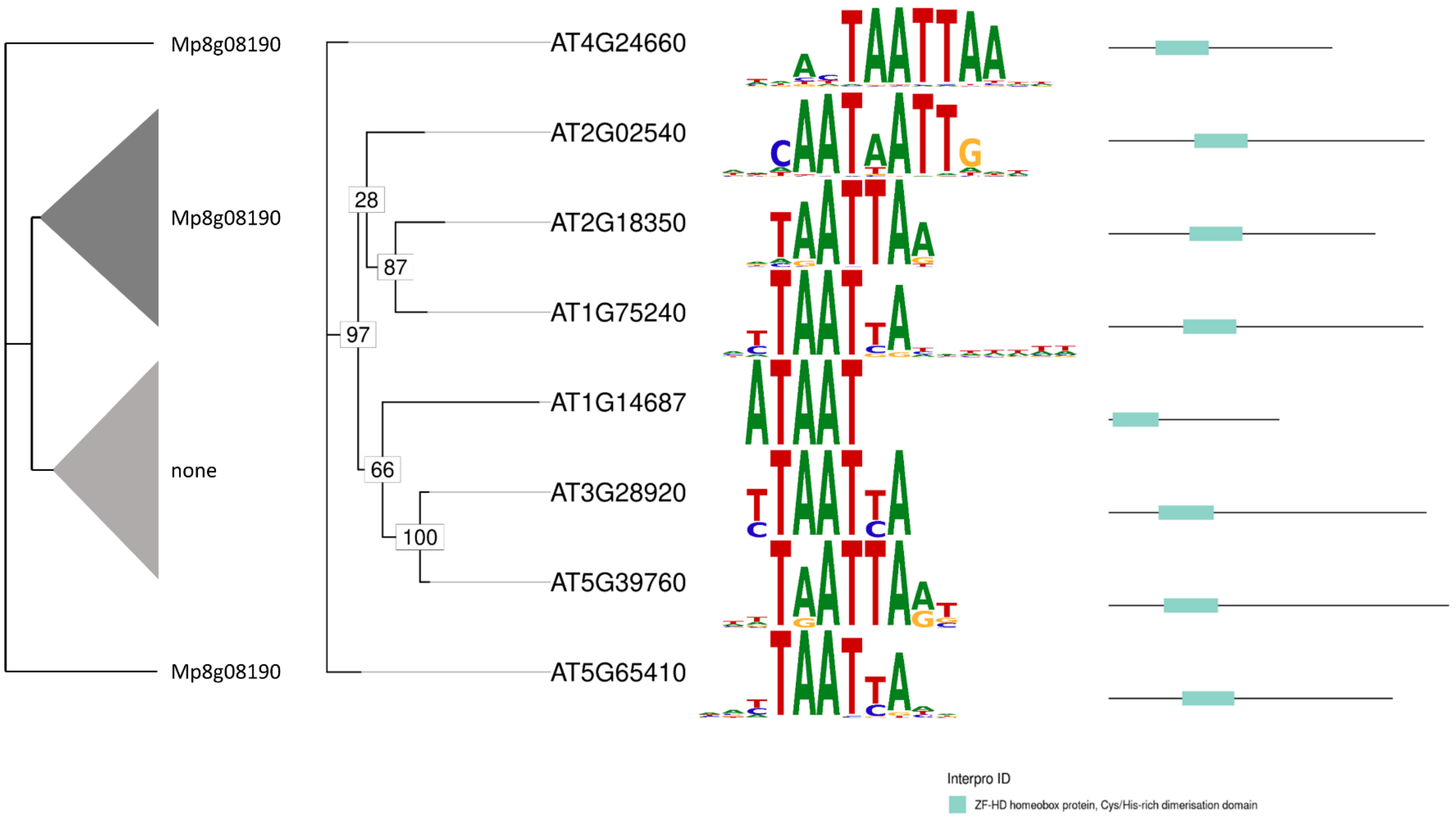


**Figure S22 Phylogenetic tree of the HD PLINC family TFs with TFBMs**

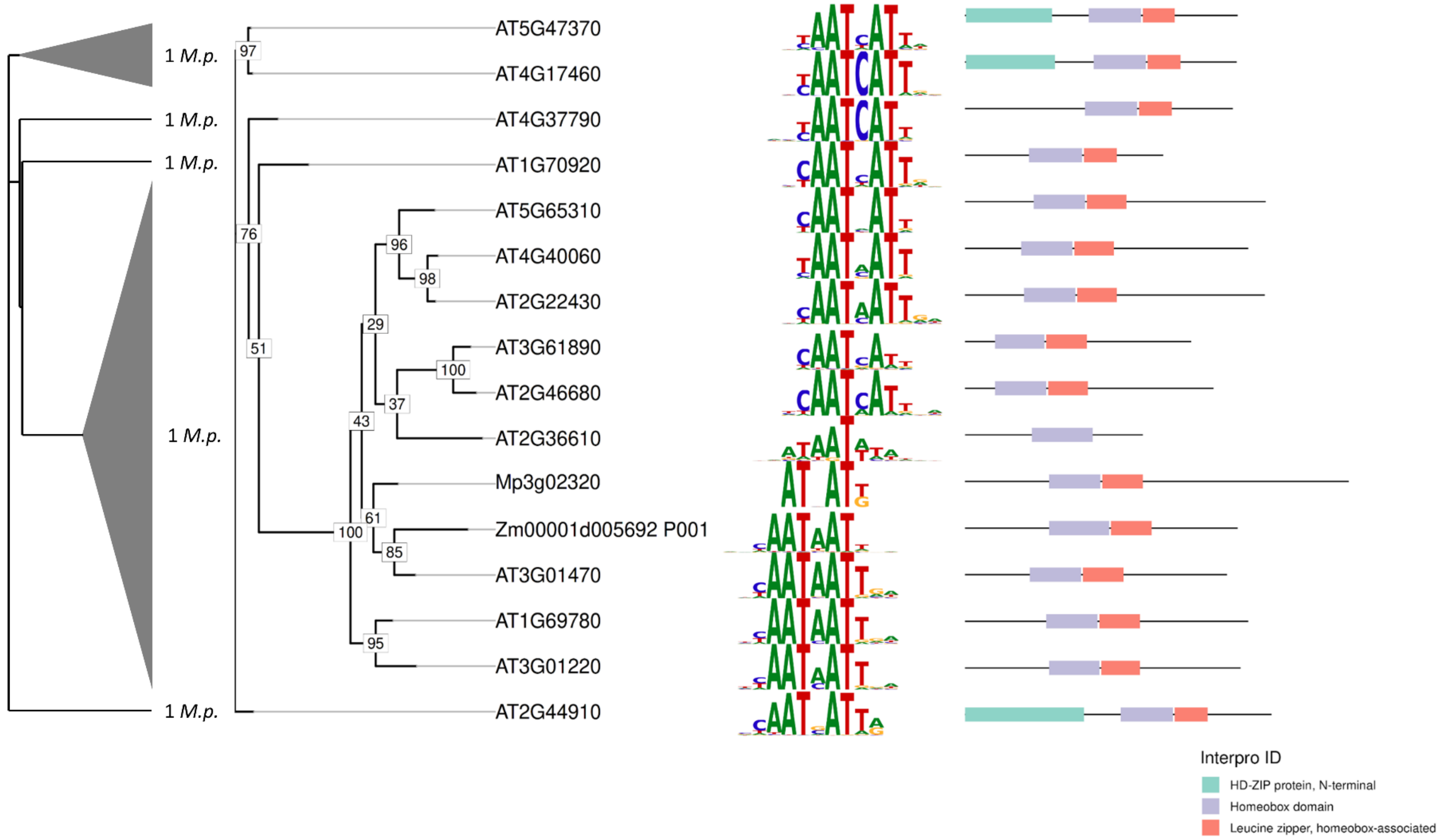


**Figure S23 Phylogenetic tree of the HD-Zip I-II family TFs with TFBMs**

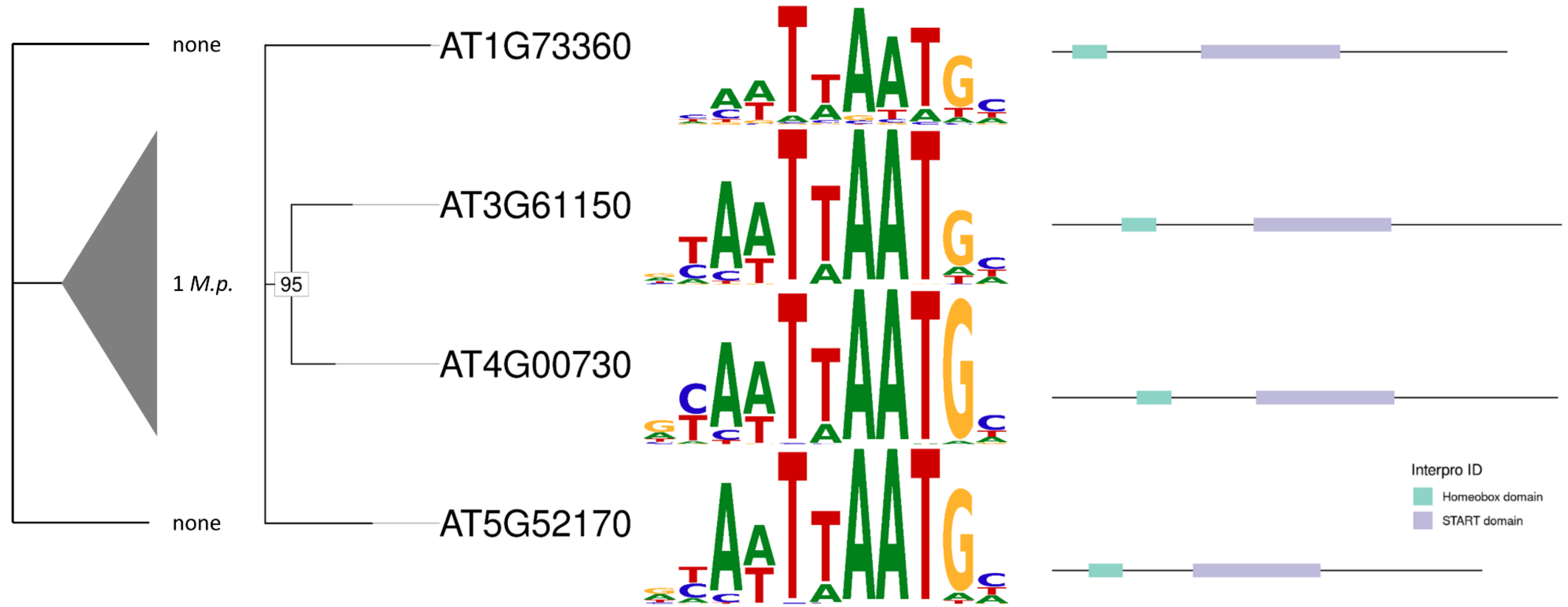


**Figure S24 Phylogenetic tree of the HD-Zip IV family TFs with TFBMs**

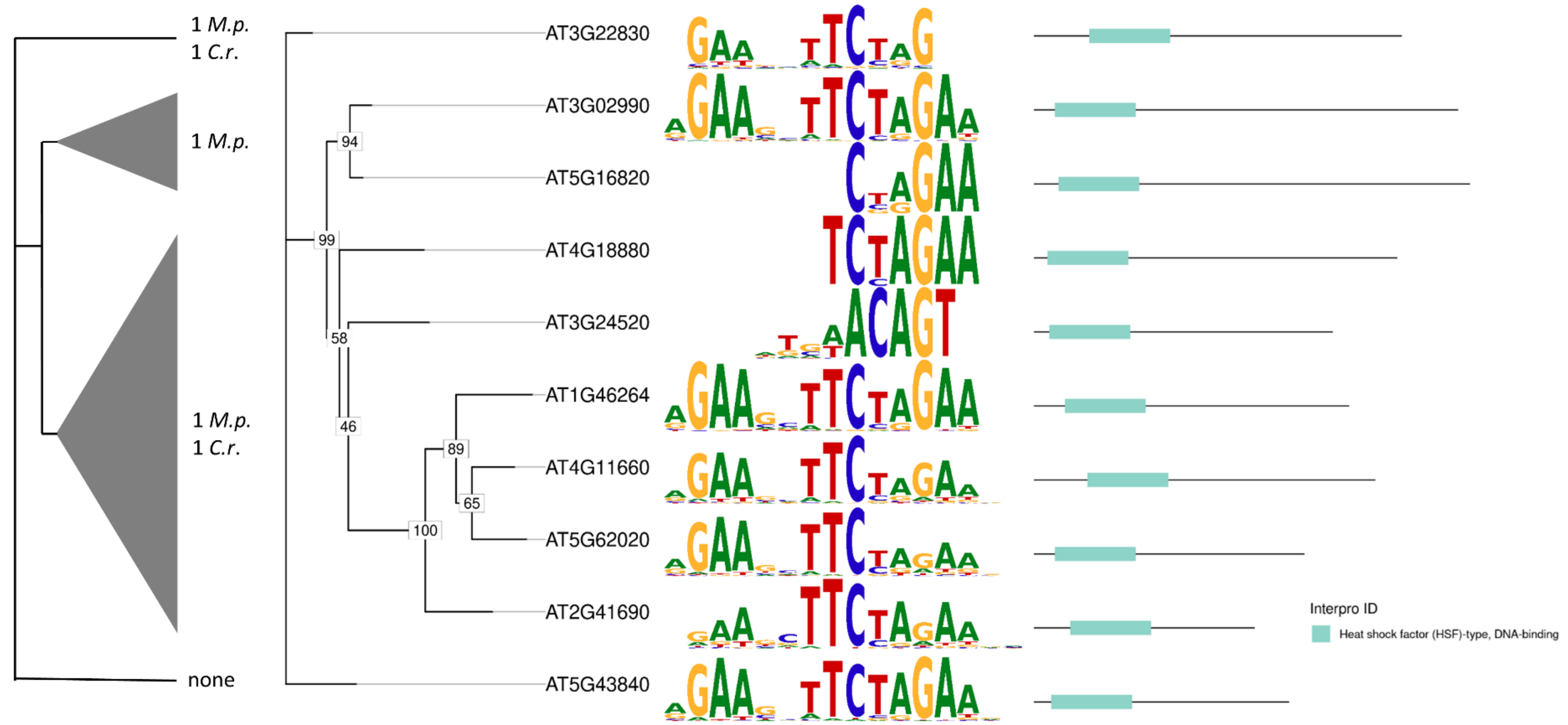


**Figure S25 Phylogenetic tree of the HSF family TFs with TFBMs**

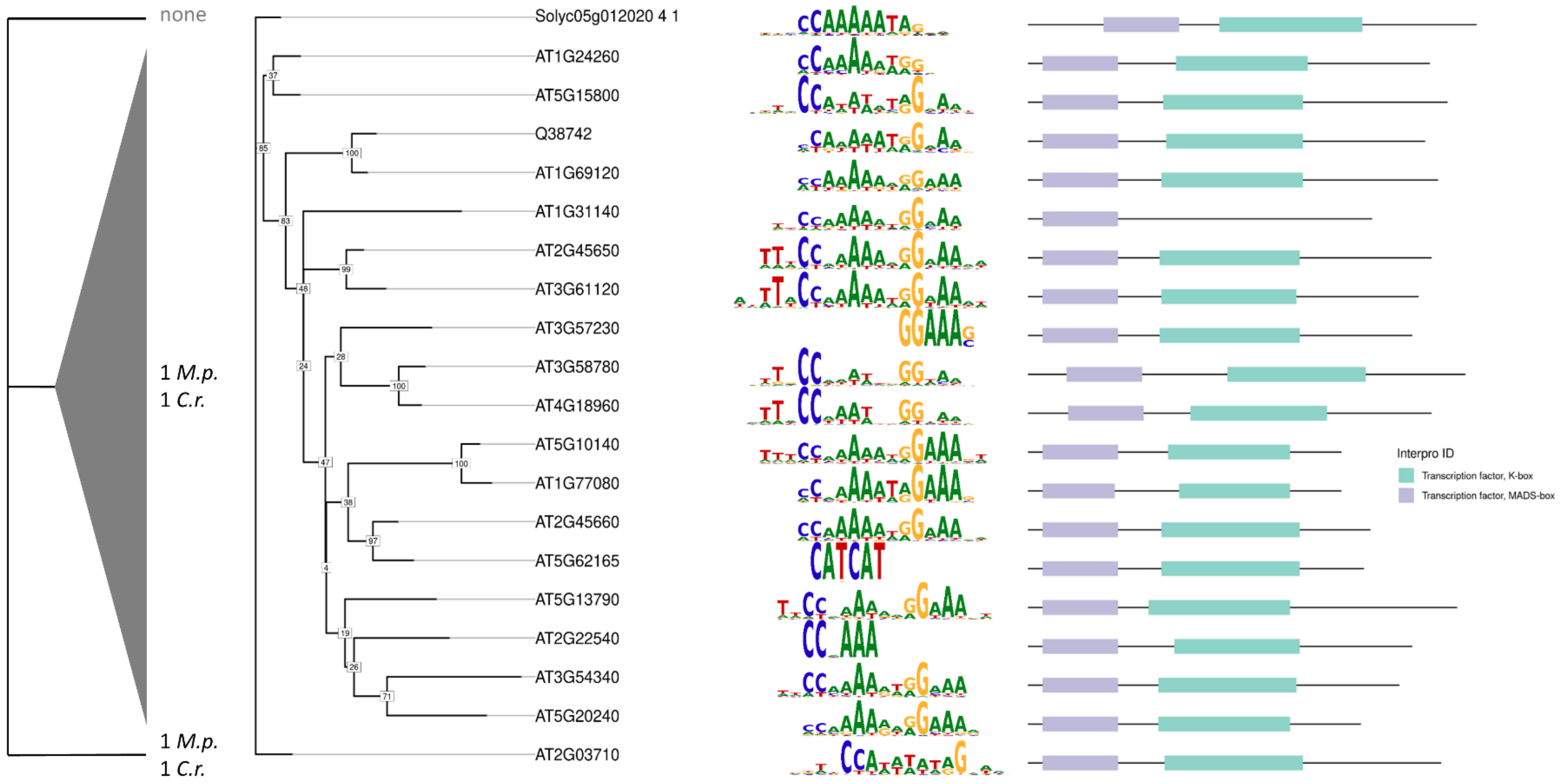


**Figure S26 Phylogenetic tree of the MADS MIKC family TFs with TFBMs**

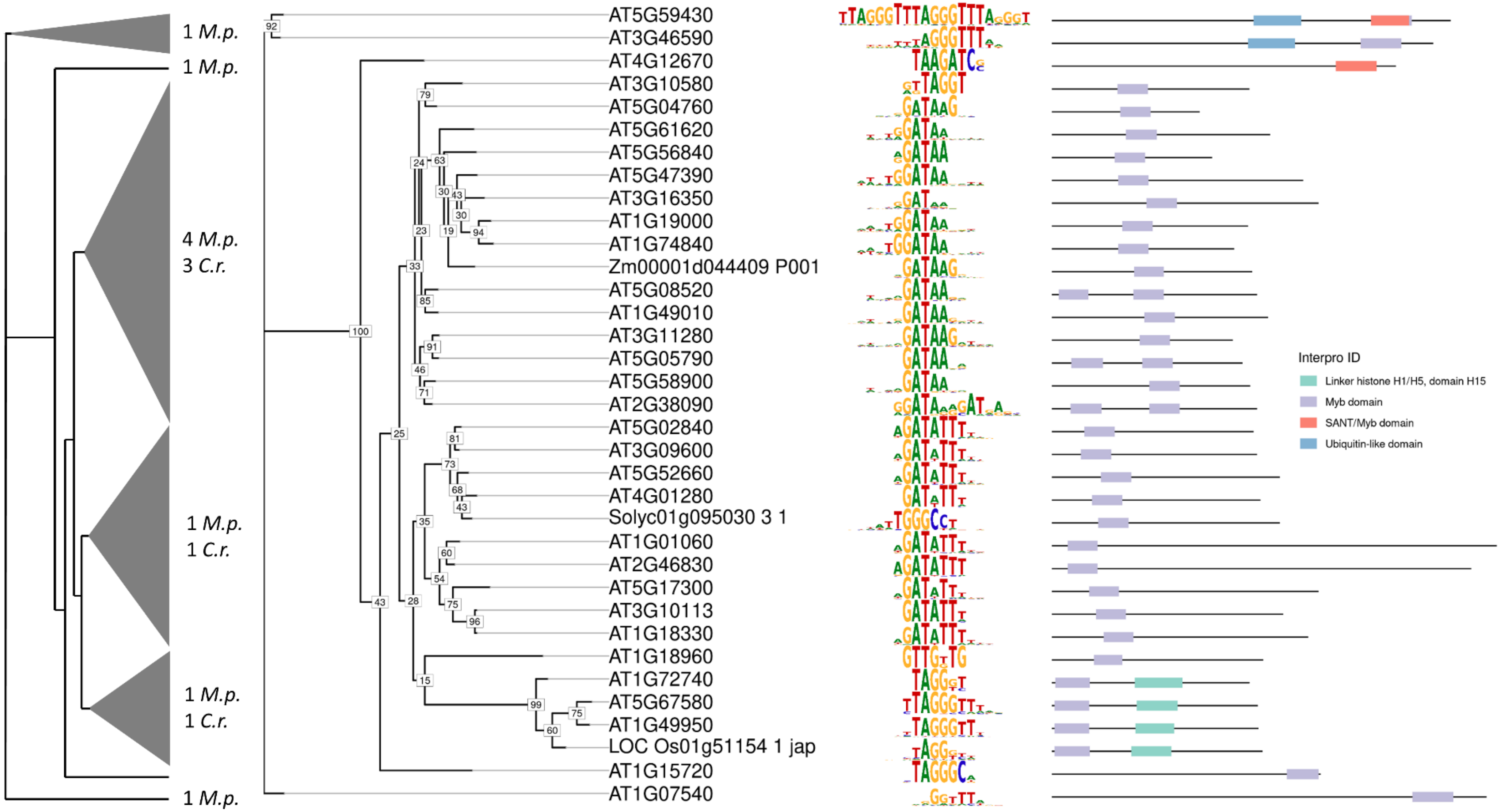


**Figure S27 Phylogenetic tree of the MYB-related family TFs with TFBMs**

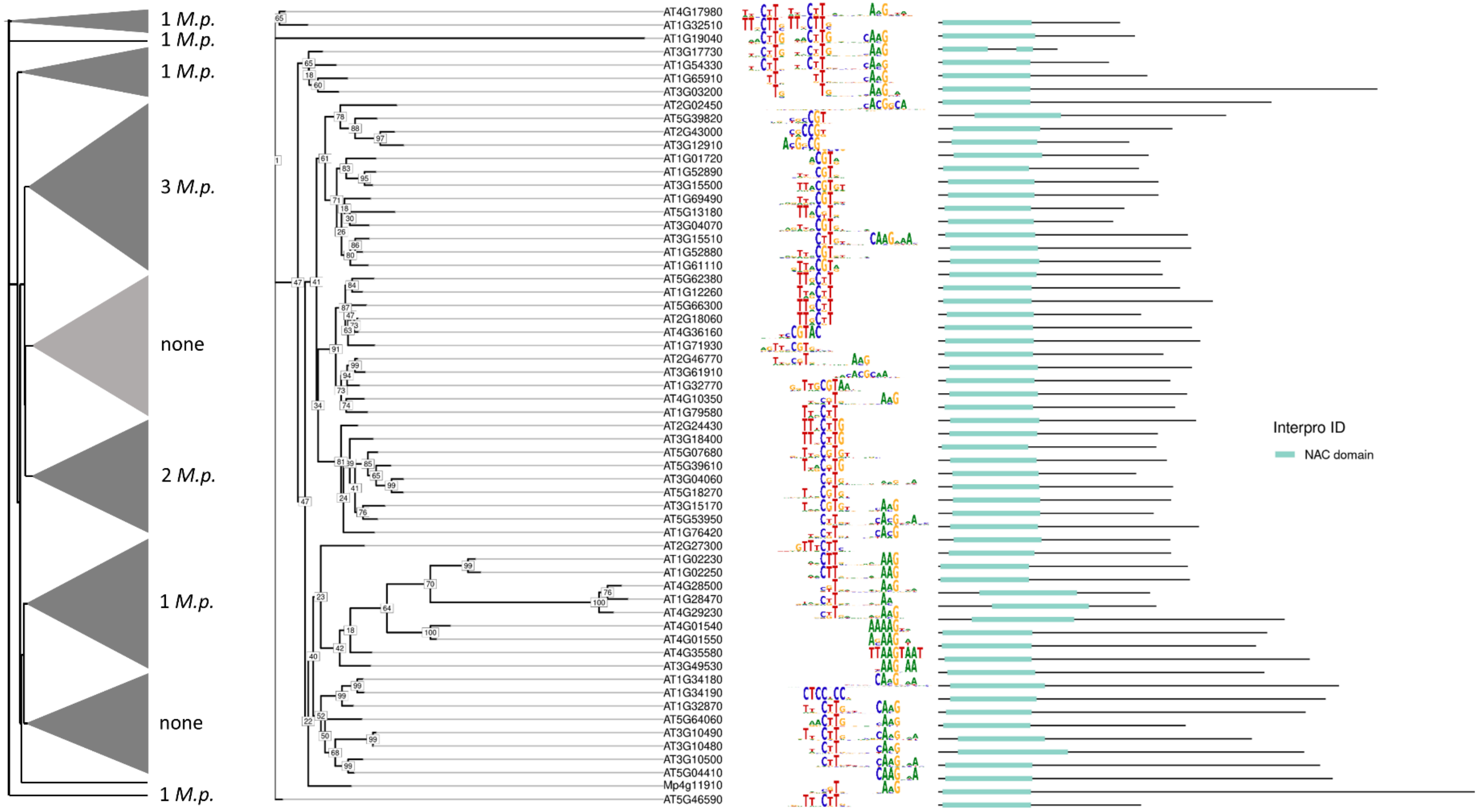


**Figure S28 Phylogenetic tree of the NAC family TFs with TFBMs**

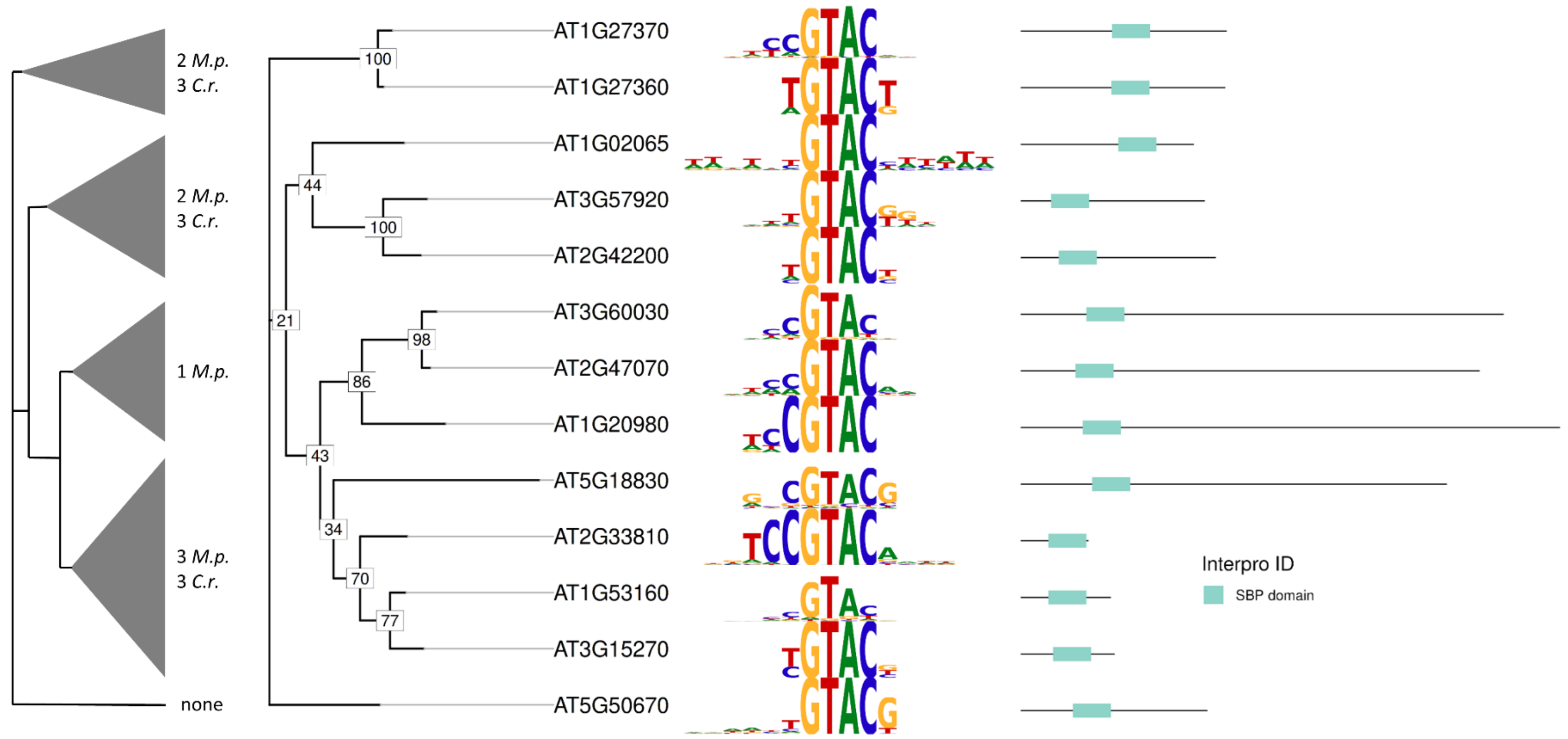


58 **Figure S29 Phylogenetic tree of the SBP family TFs with TFBMs**

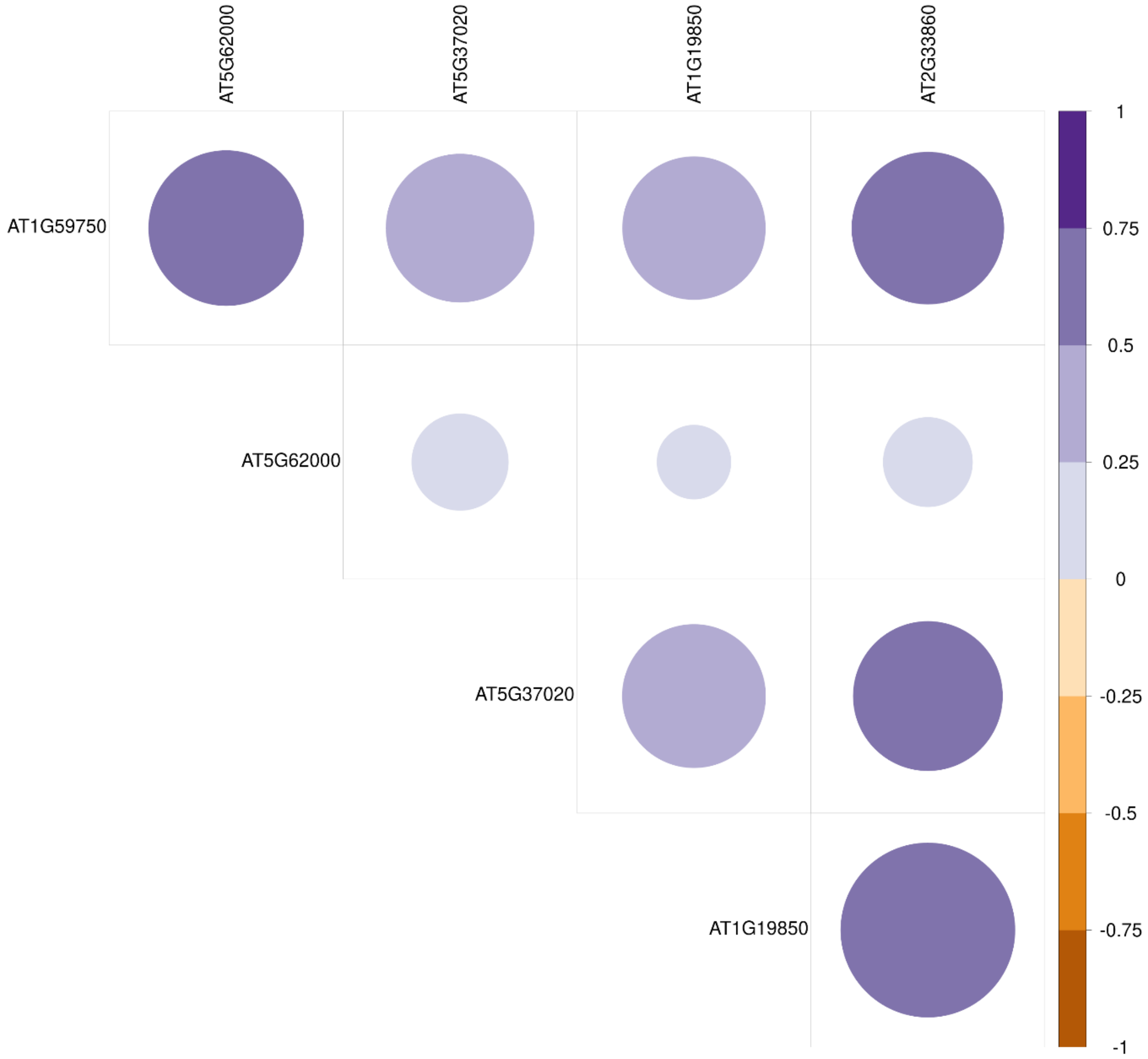


**Figure S30 Correlation matrix of ARF family TF expression in *A. thaliana***

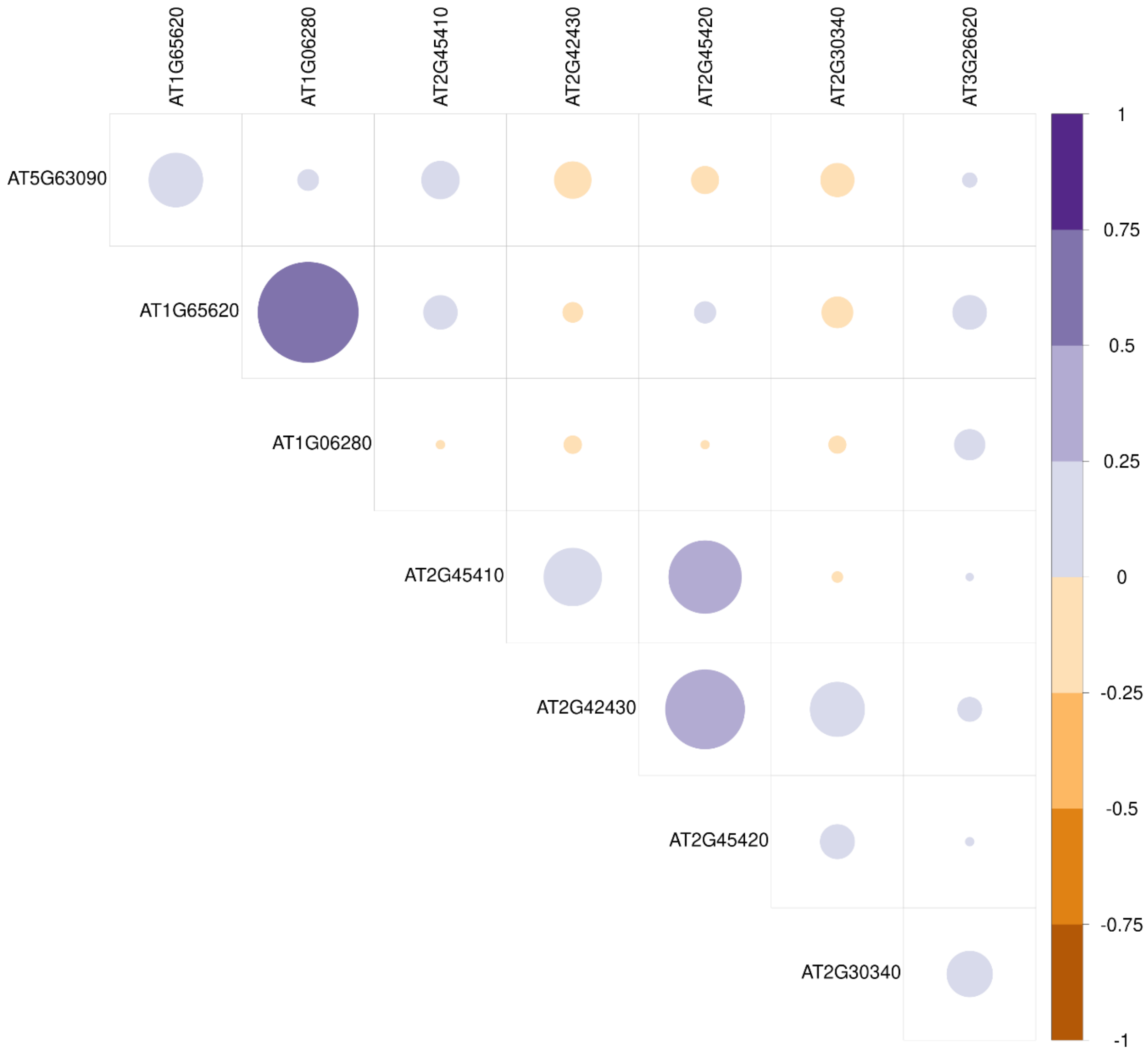


61

62 **Figure S31 Correlation matrix of AS2/LOB family TF expression in *A. thaliana***

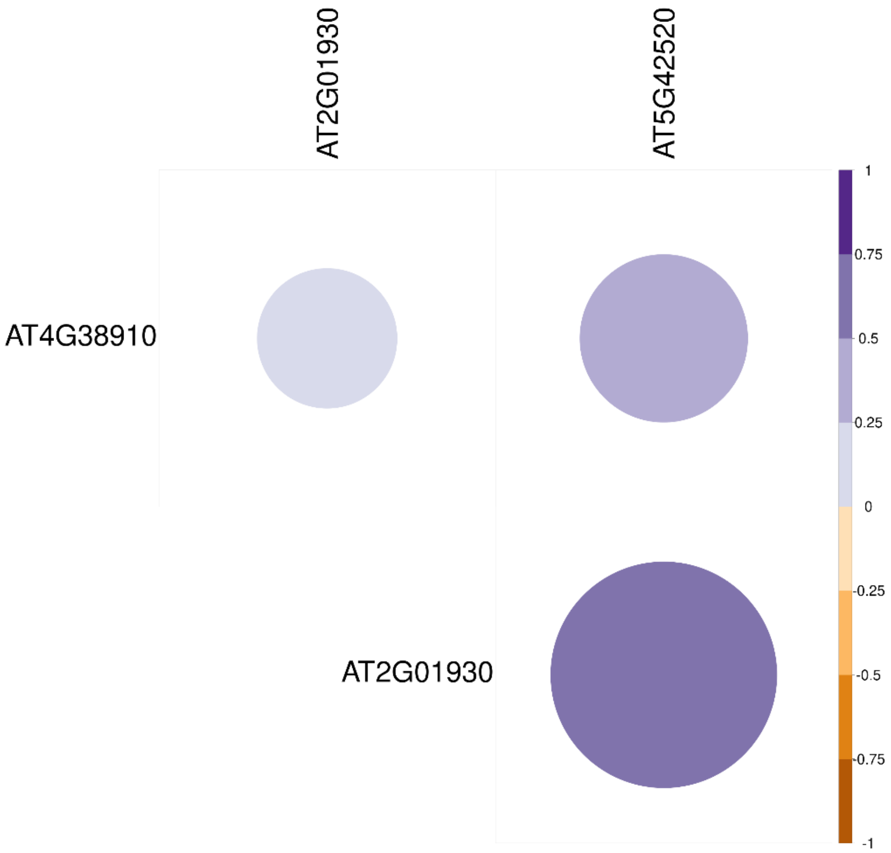


**Figure S32 Correlation matrix of BBR/BPC family TF expression in *A. thaliana***

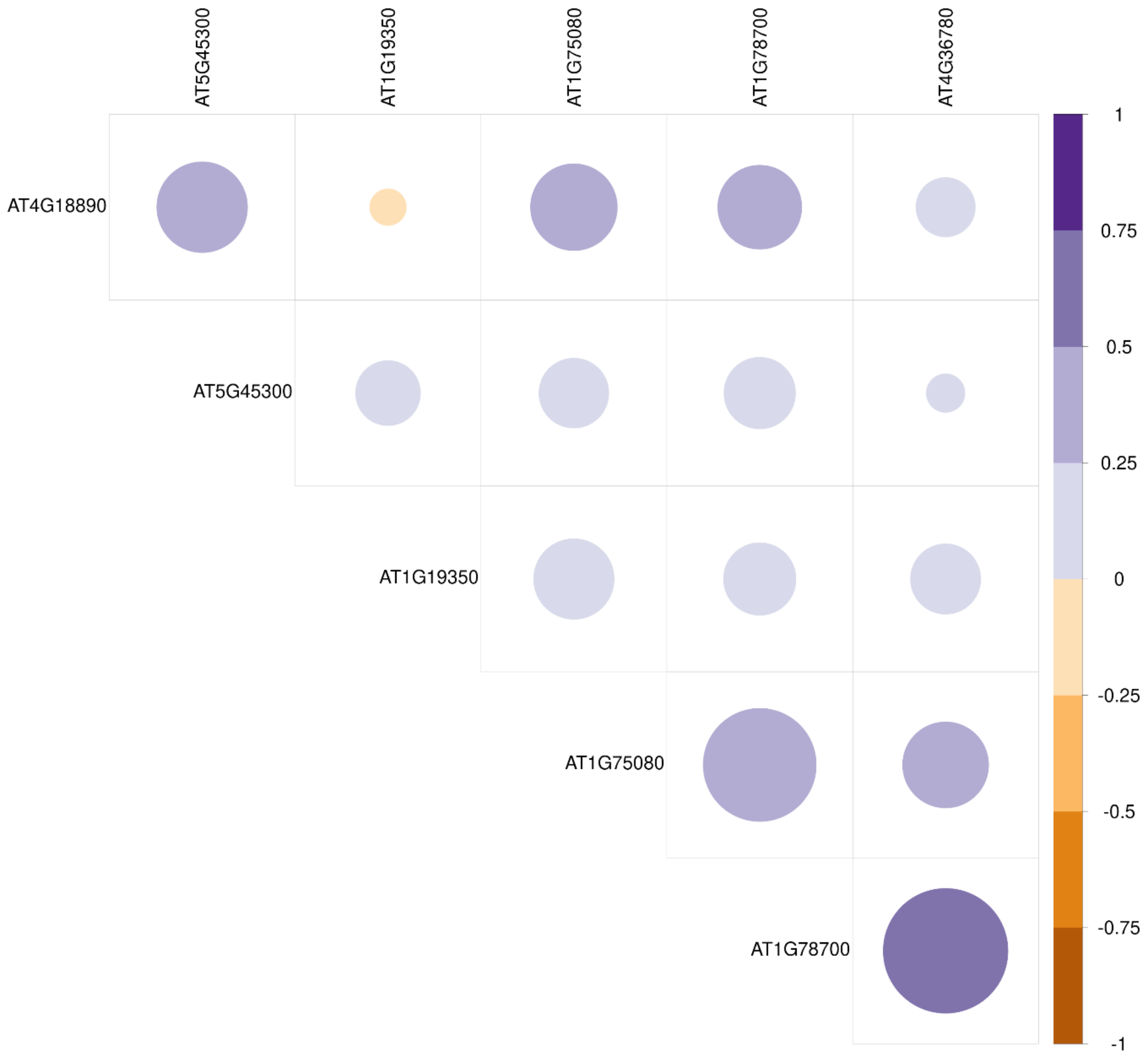


**Figure S33 Correlation matrix of BES1 family TF expression in *A. thaliana***

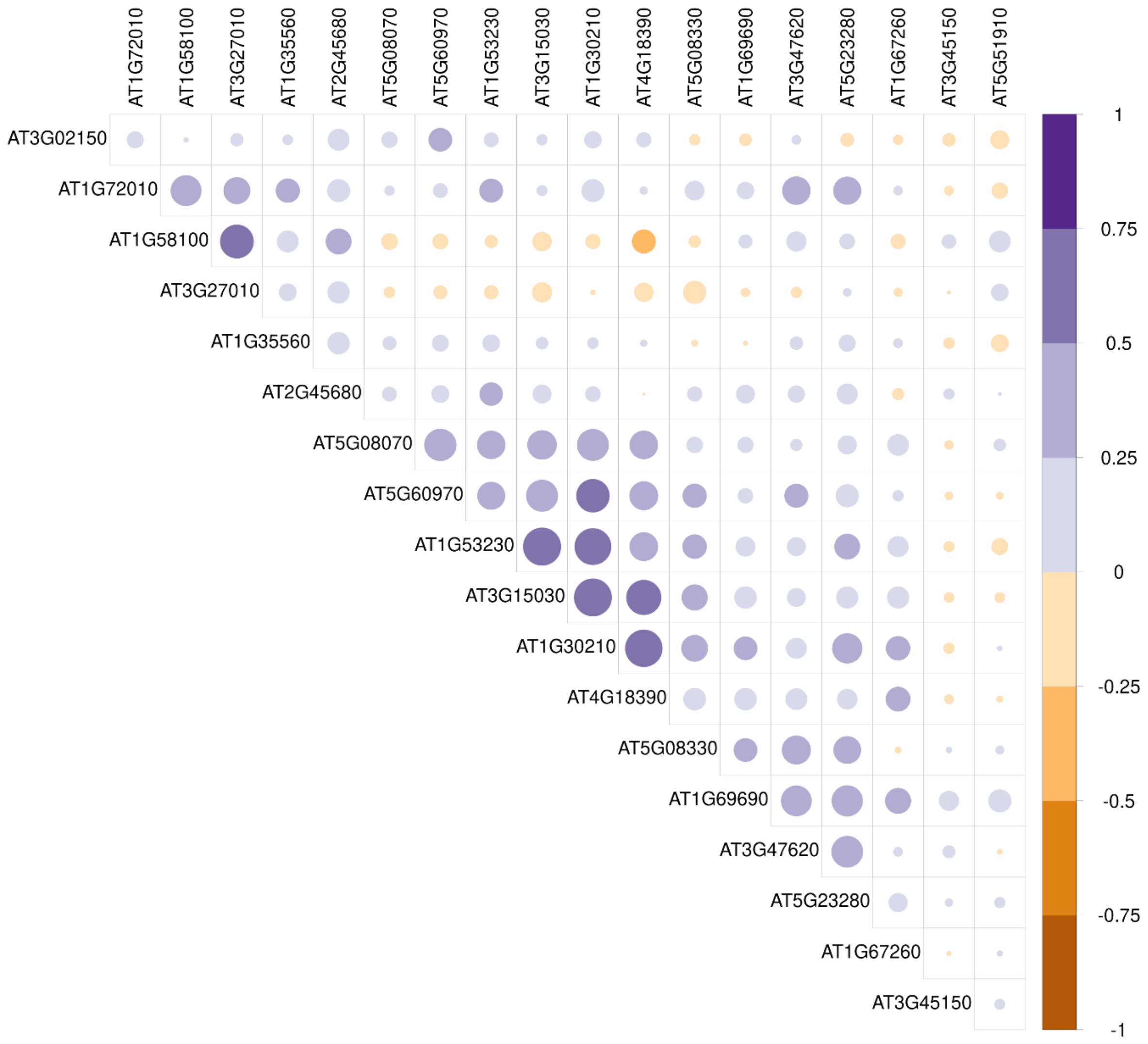


67

68 **Figure S34 Correlation matrix of bHLH TCP family TF expression in *A. thaliana***

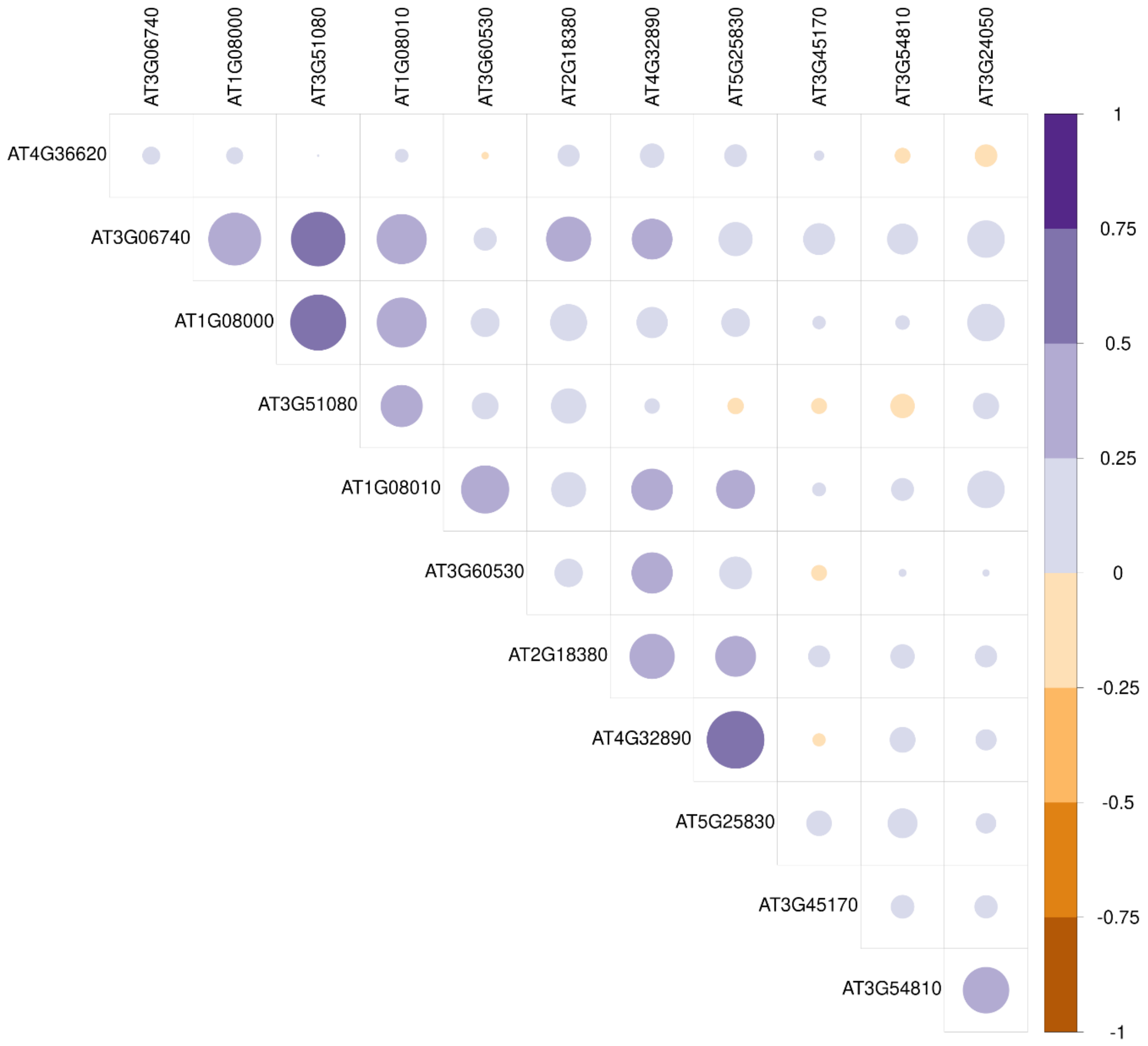


**Figure S35 Correlation matrix of C2C2 GATA family TF expression in *A. thaliana***

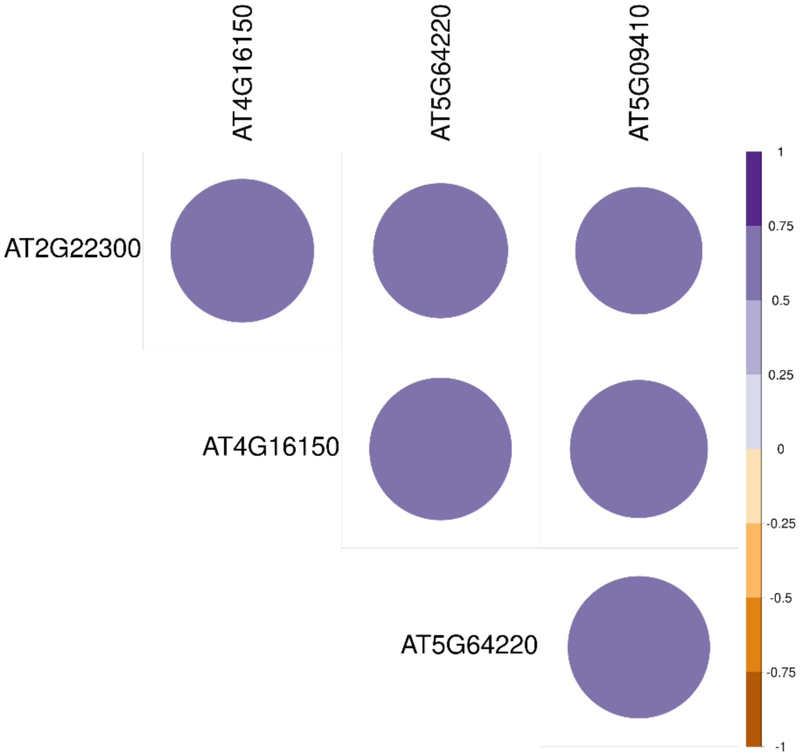


**Figure S36 Correlation matrix of CAMTA family TF expression in *A. thaliana***

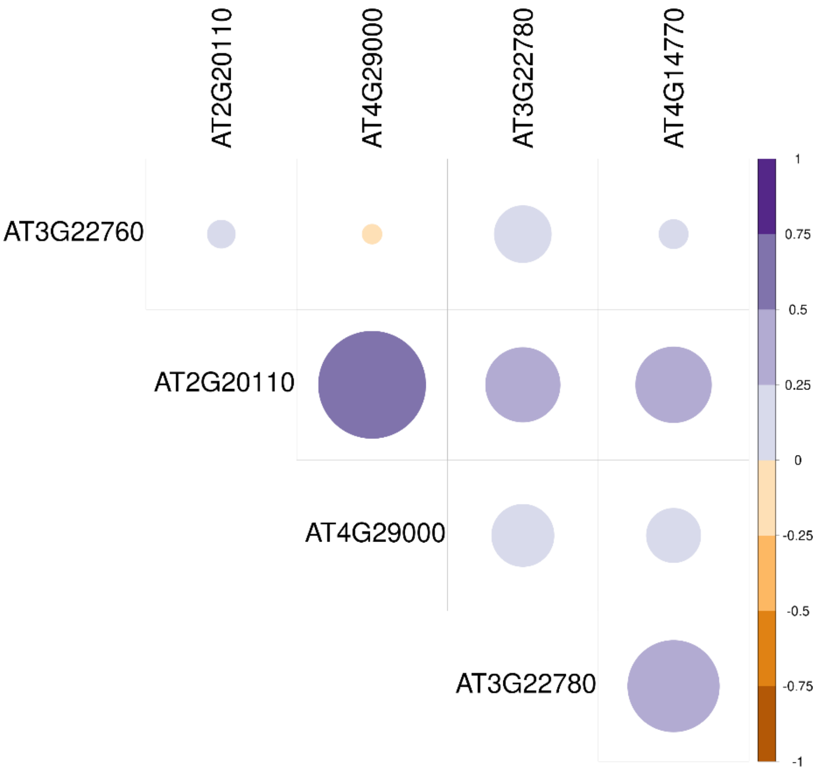


73

74 **Figure S37 Correlation matrix of CPP family TF expression in *A. thaliana***

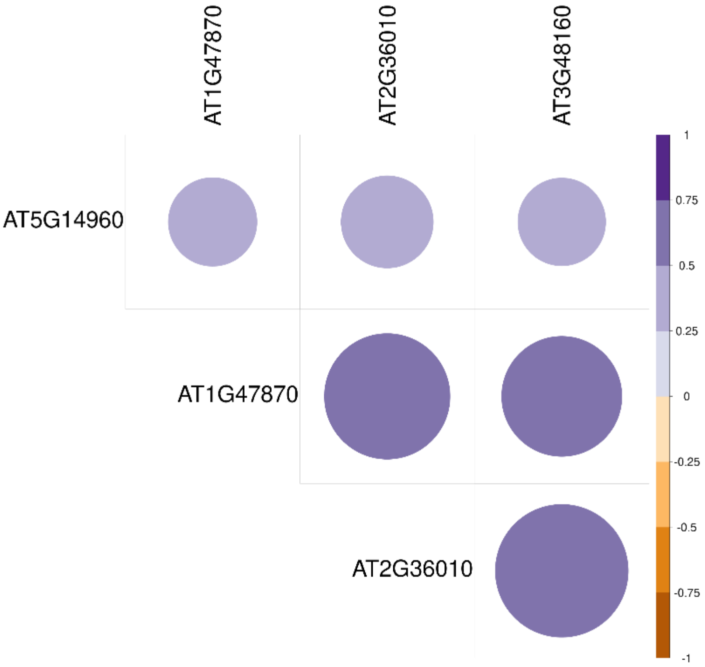


**Figure S38 Correlation matrix of E2F/DP family TF expression in *A. thaliana***

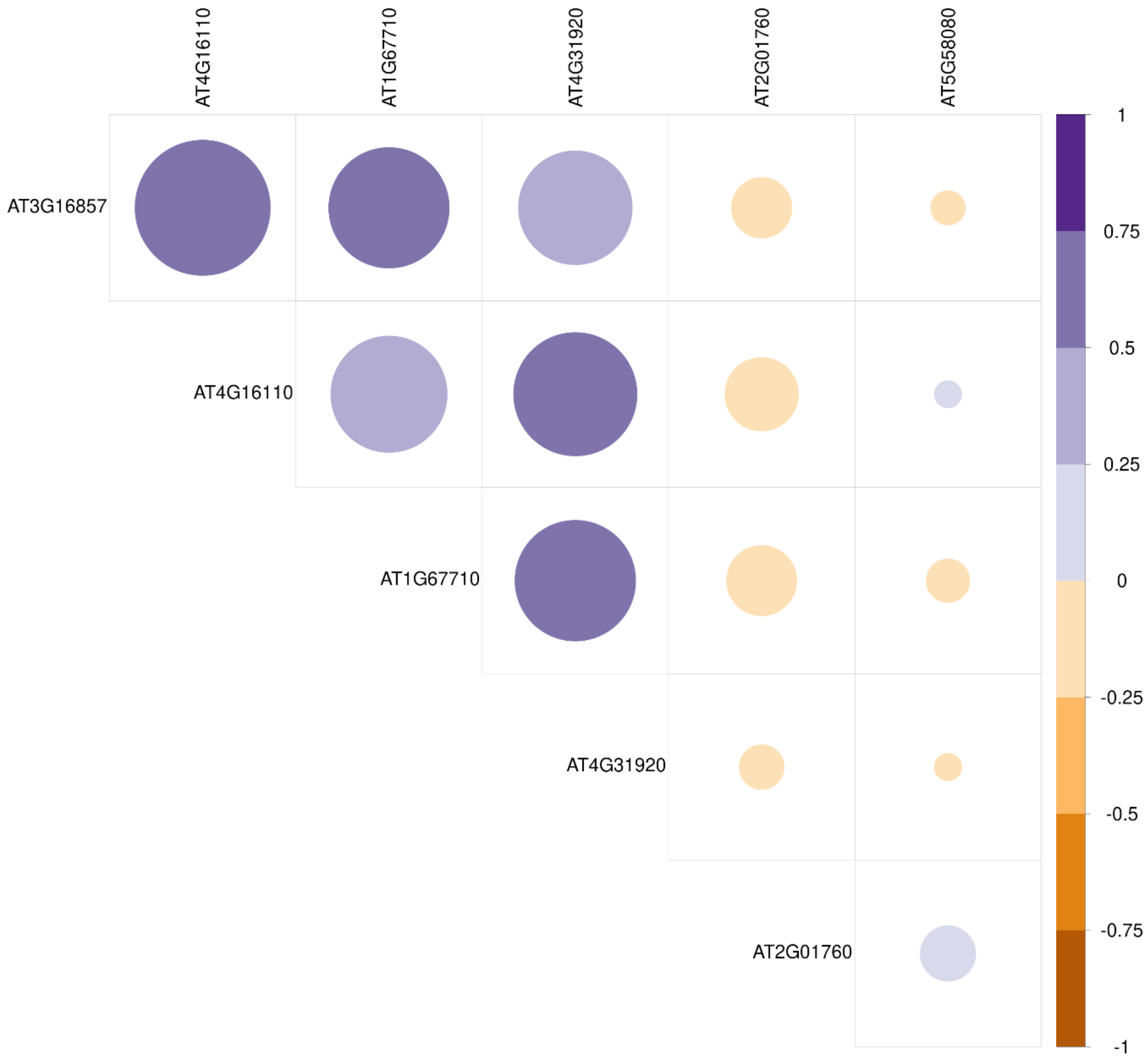


**Figure S39 Correlation matrix of GARP ARR-B family TF expression in *A. thaliana***

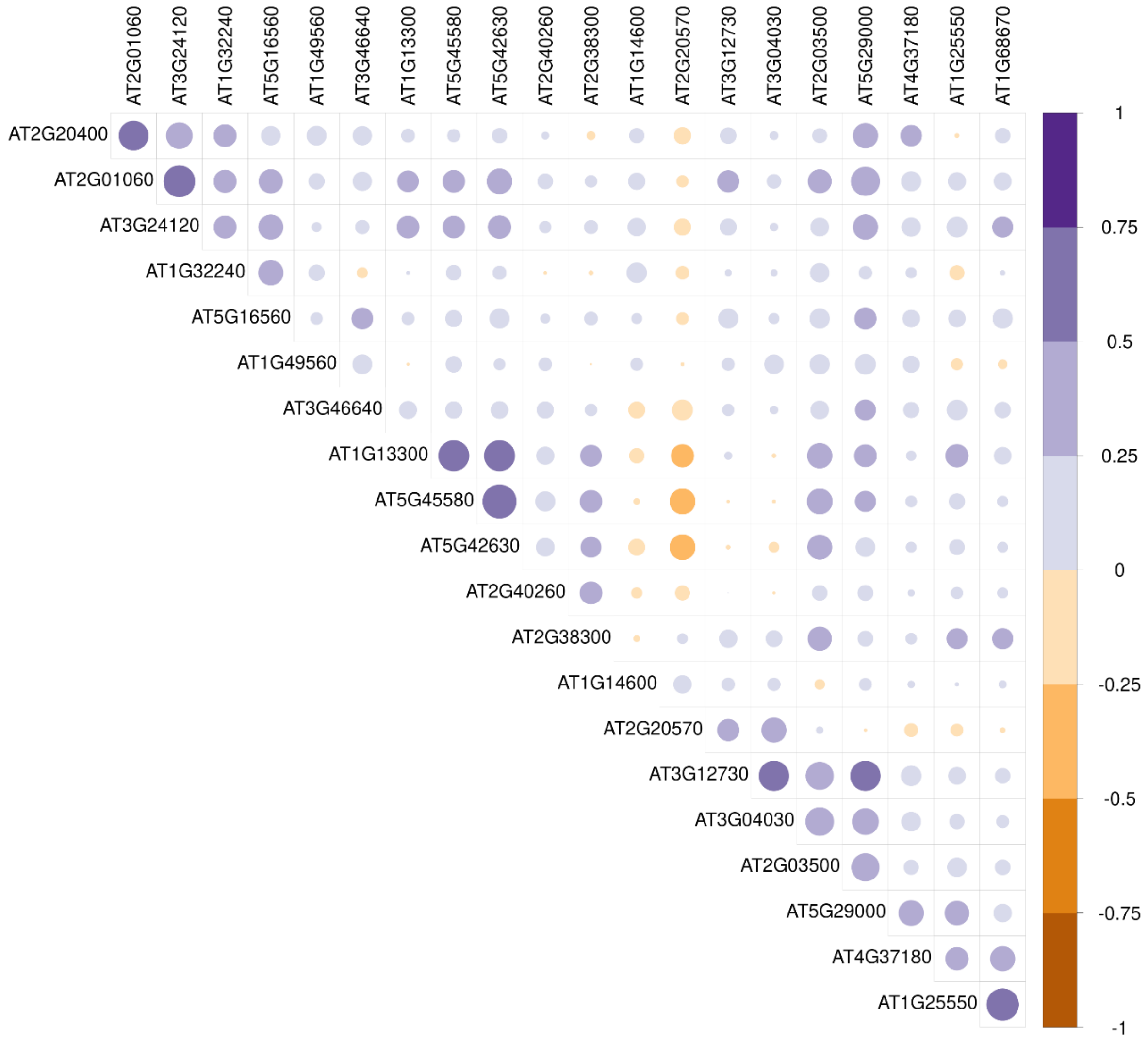


79

80 **Figure S40 Correlation matrix of ARF family TF expression in *A. thaliana***

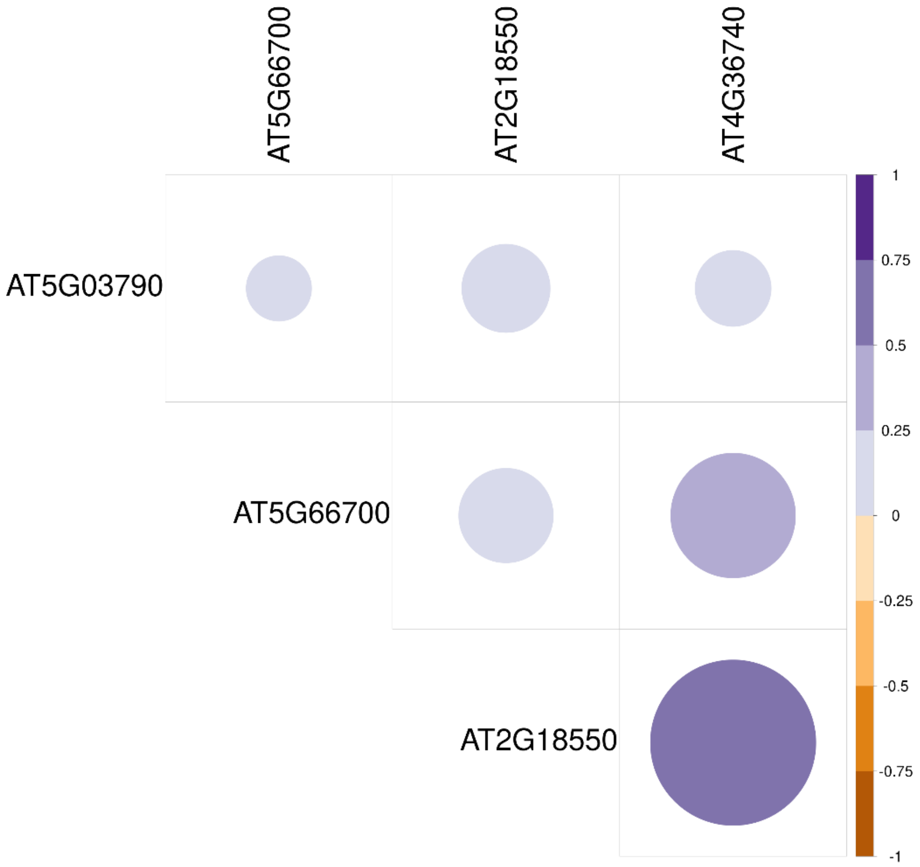


81
82 **Figure S41 Correlation matrix of HD family TF expression in *A. thaliana***

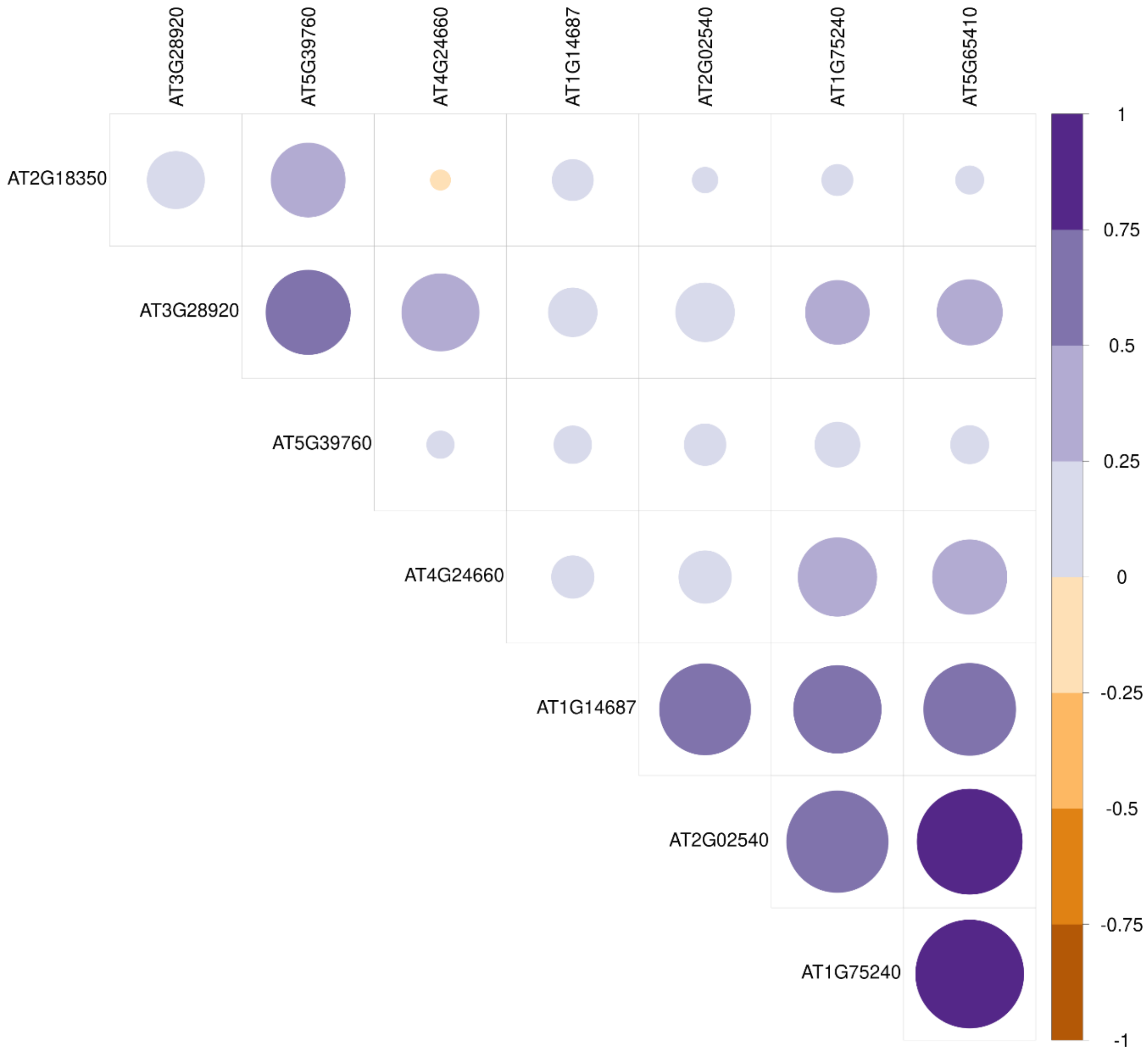


**Figure S42 Correlation matrix of HD PLINC family TF expression in *A. thaliana***

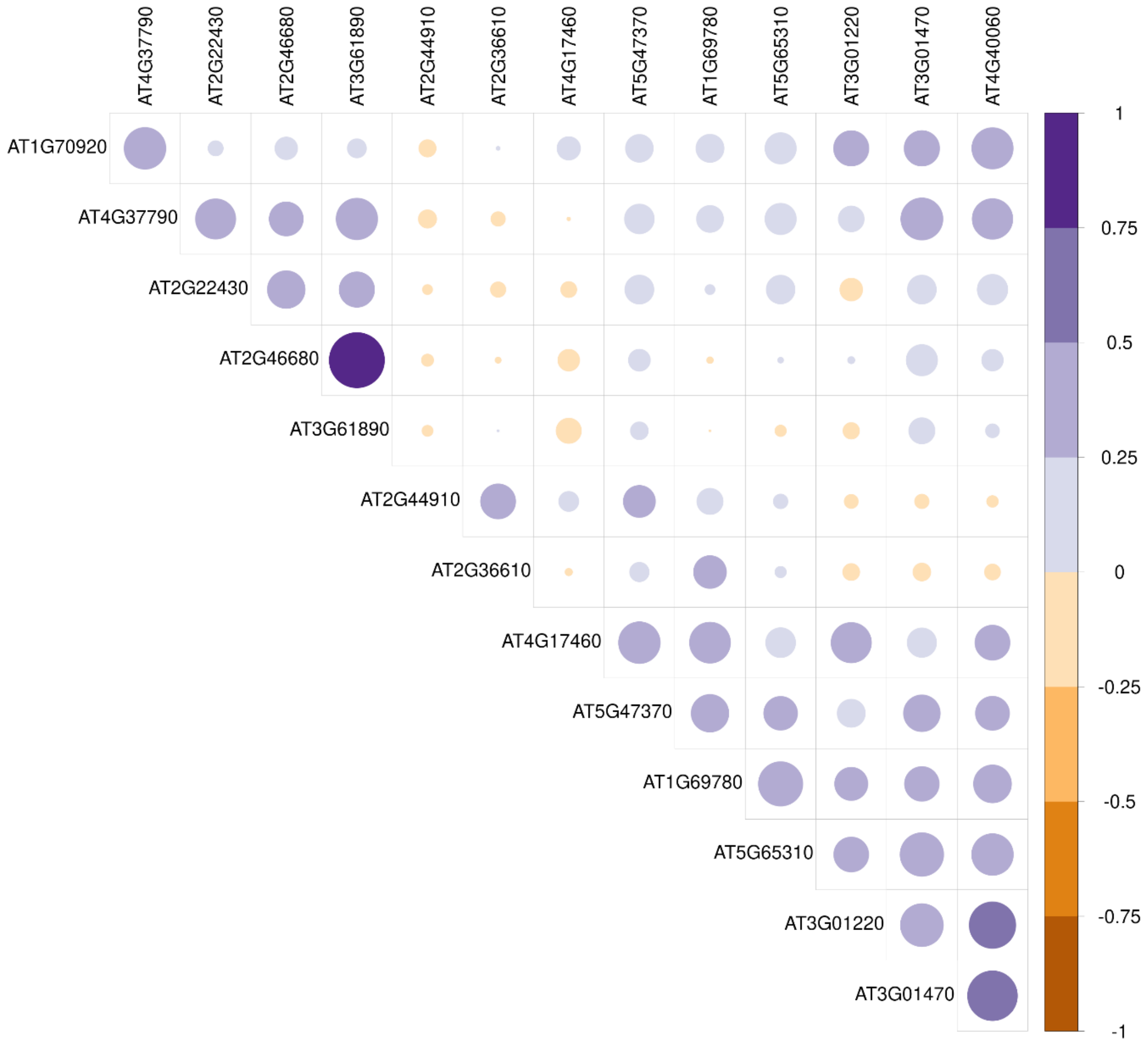


**Figure S43 Correlation matrix of HD-Zip I-II family TF expression in *A. thaliana***

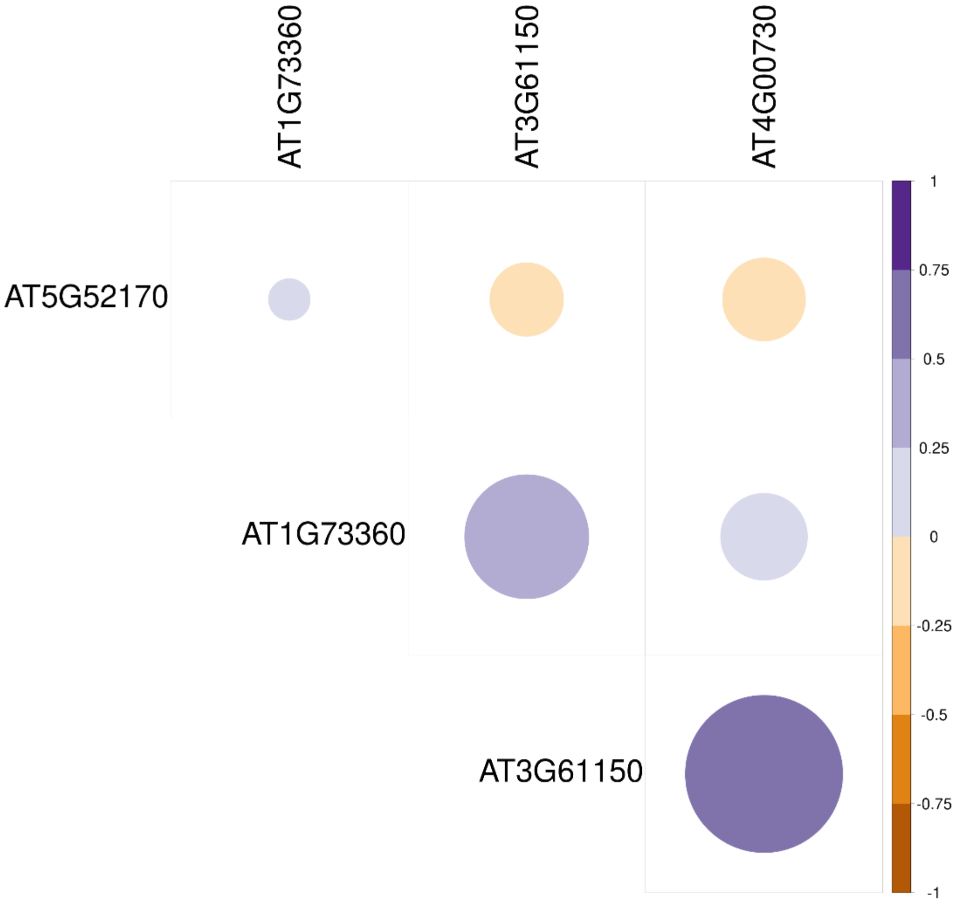


**Figure S44 Correlation matrix of HD-Zip IV family TF expression in *A. thaliana***

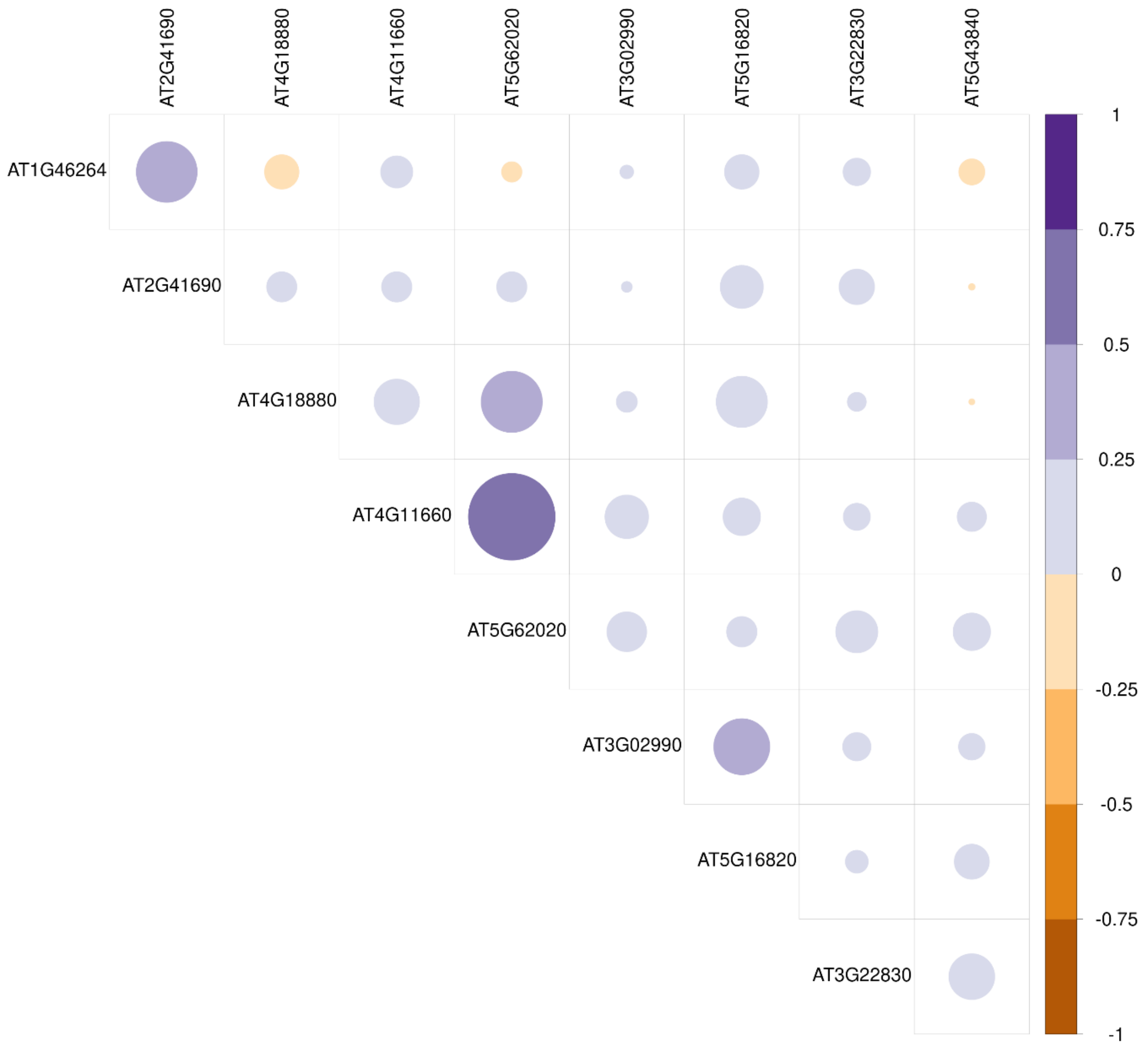


**Figure S45 Correlation matrix of HSF family TF expression in *A. thaliana***

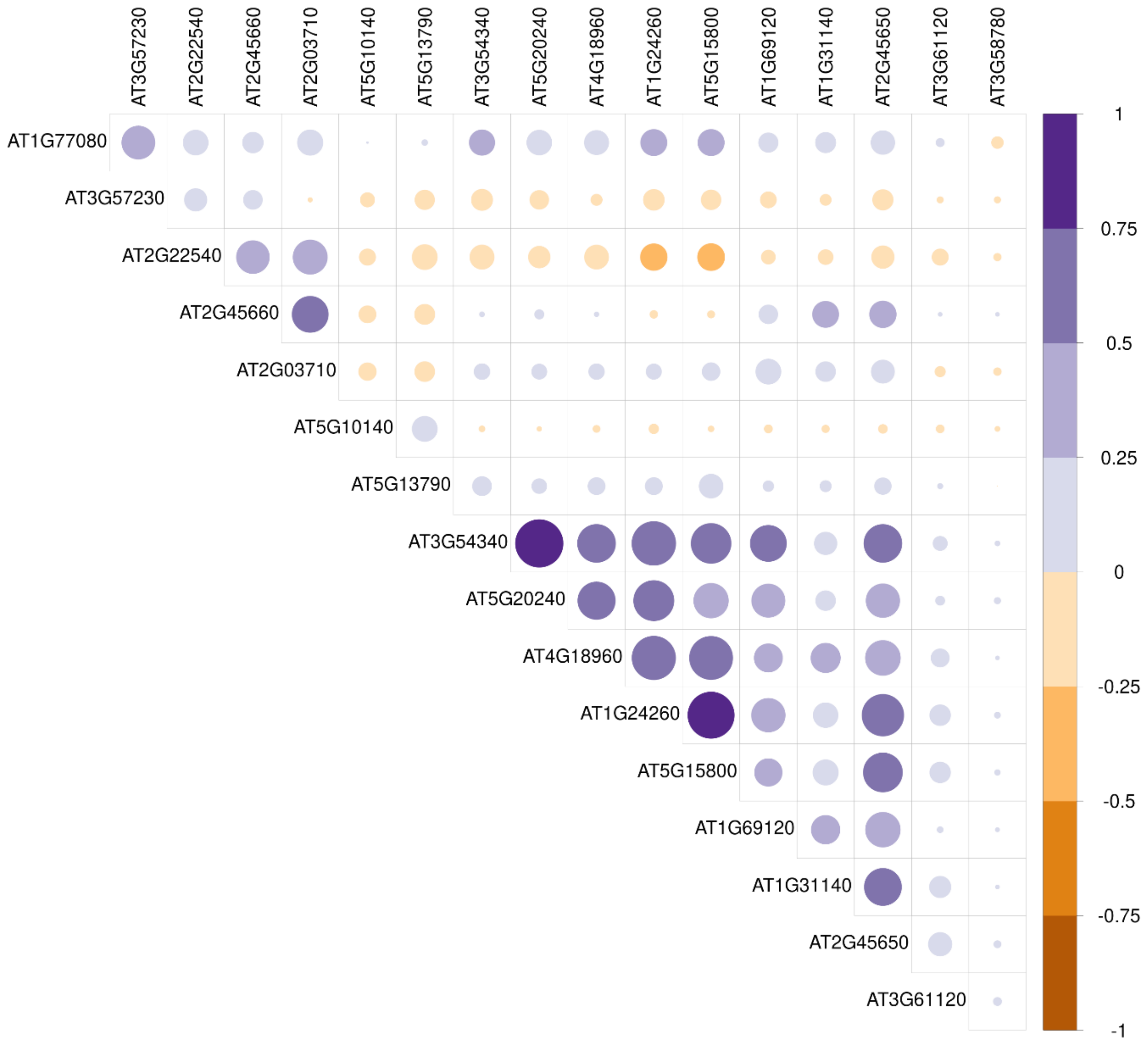


**Figure S46 Correlation matrix of MADS MIKC family TF expression in *A. thaliana***

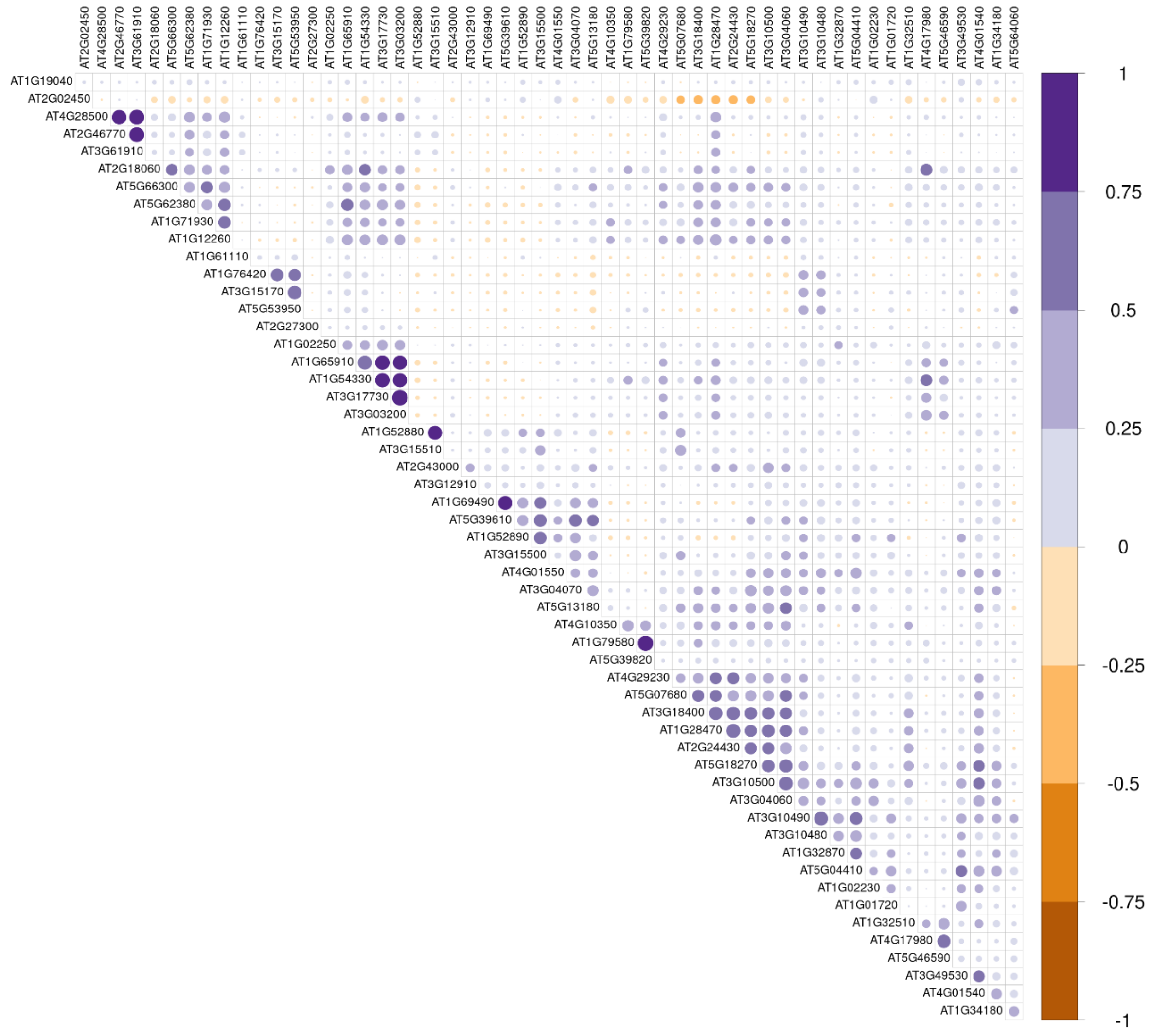


93

94 **Figure S47 Correlation matrix of NAC family TF expression in *A. thaliana***

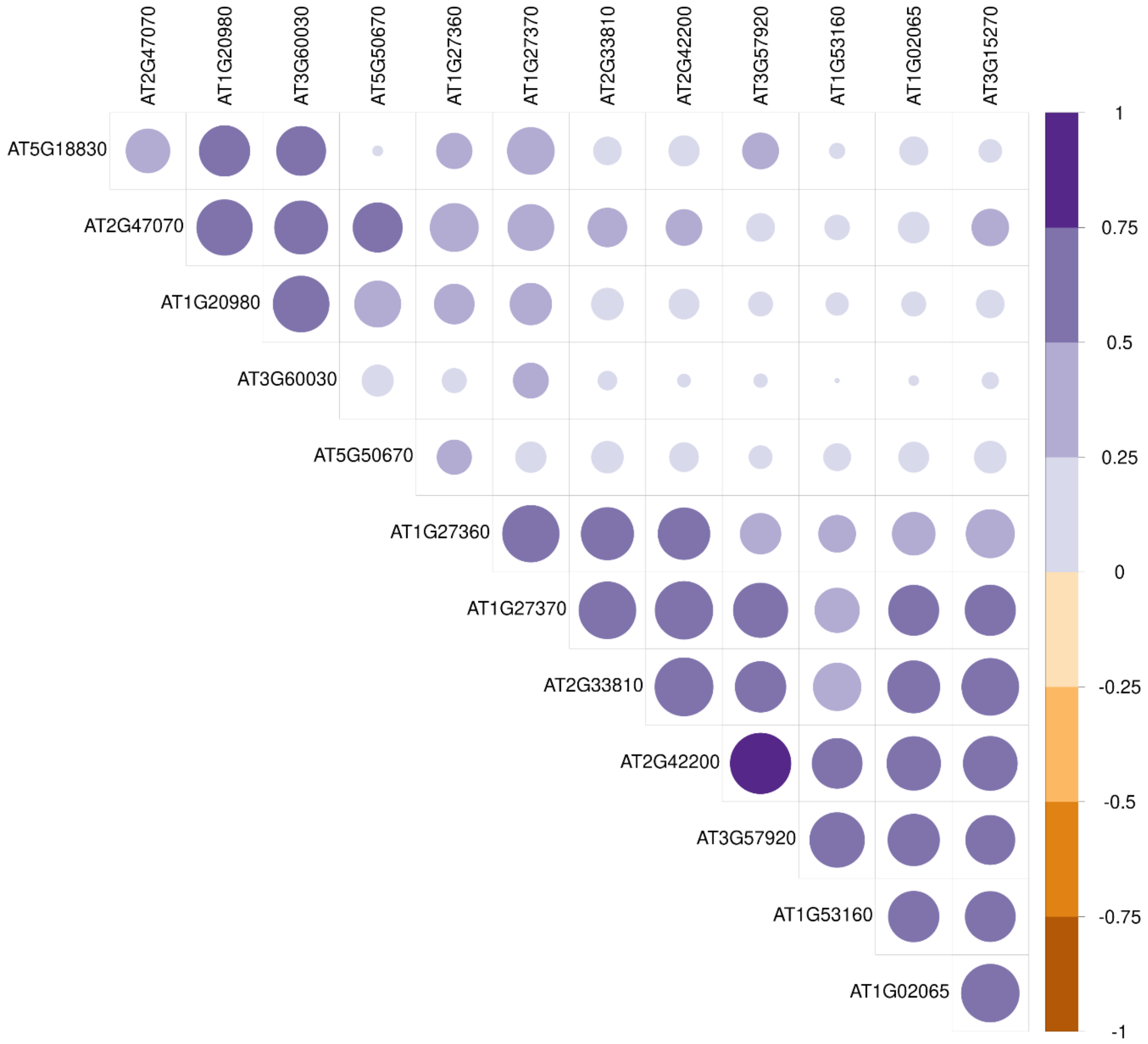


**Figure S48 Correlation matrix of SBP family TF expression in *A. thaliana***

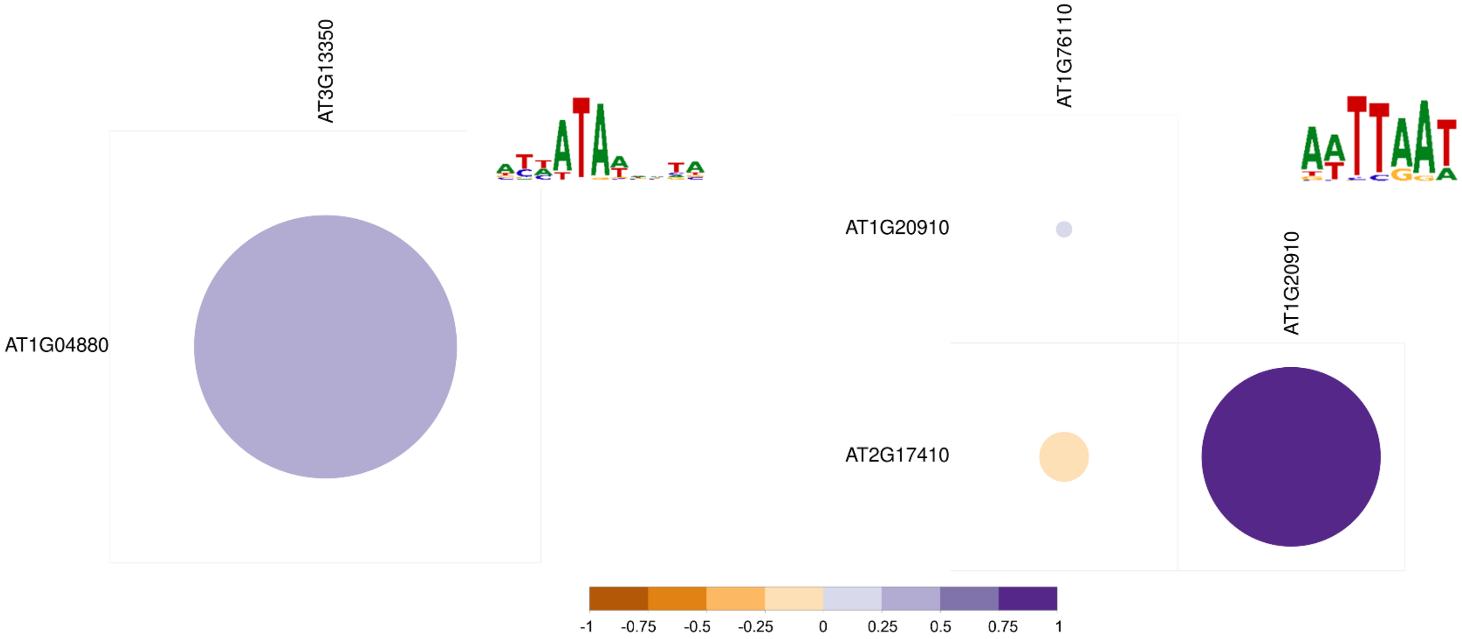


**Figure S49 Correlation matrices of ARID TF expression within groups binding the same TFBM in *A. thaliana***

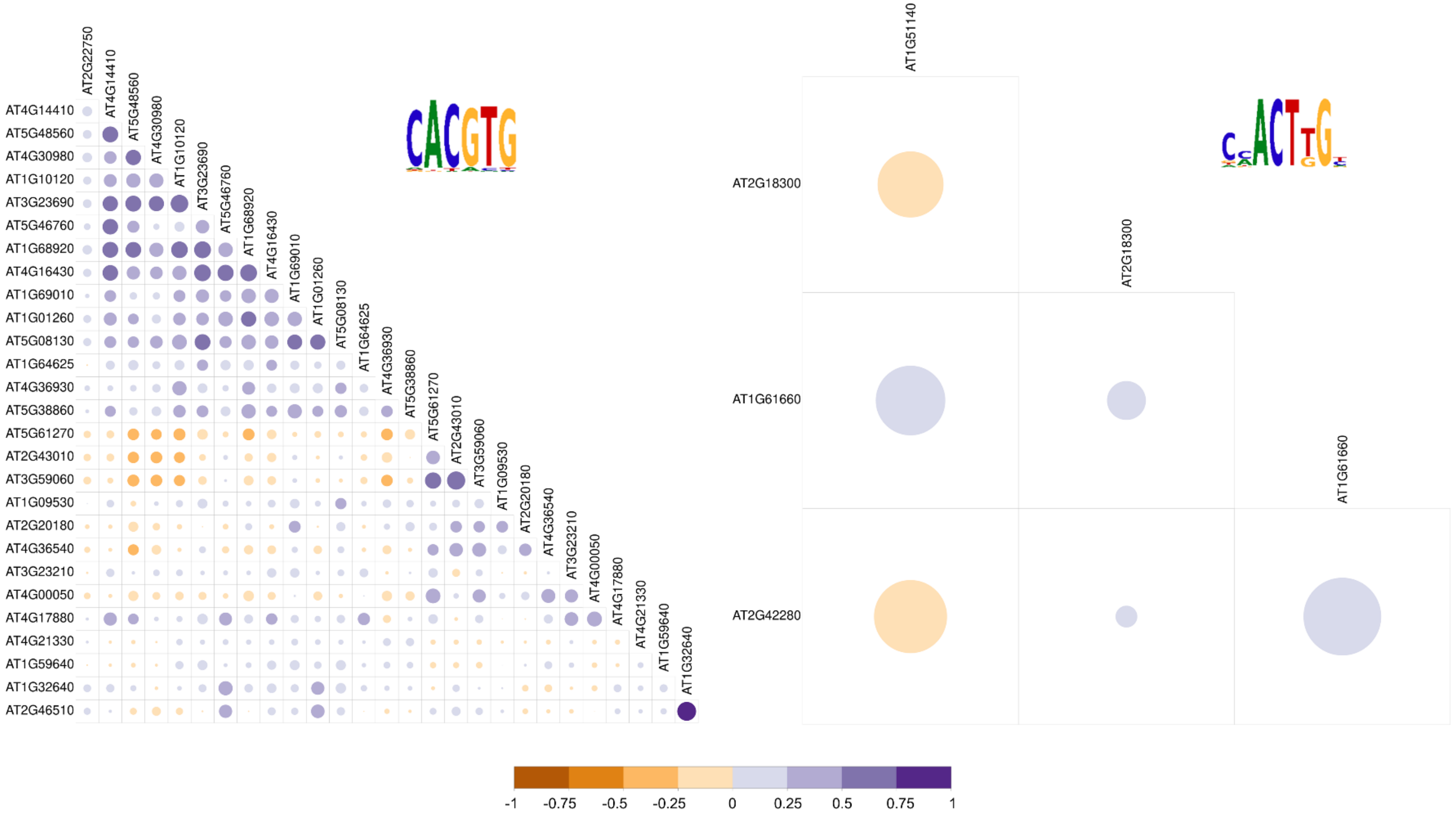


**Figure S50 Correlation matrices of bHLH TF expression within groups binding the same TFBM in *A. thaliana***

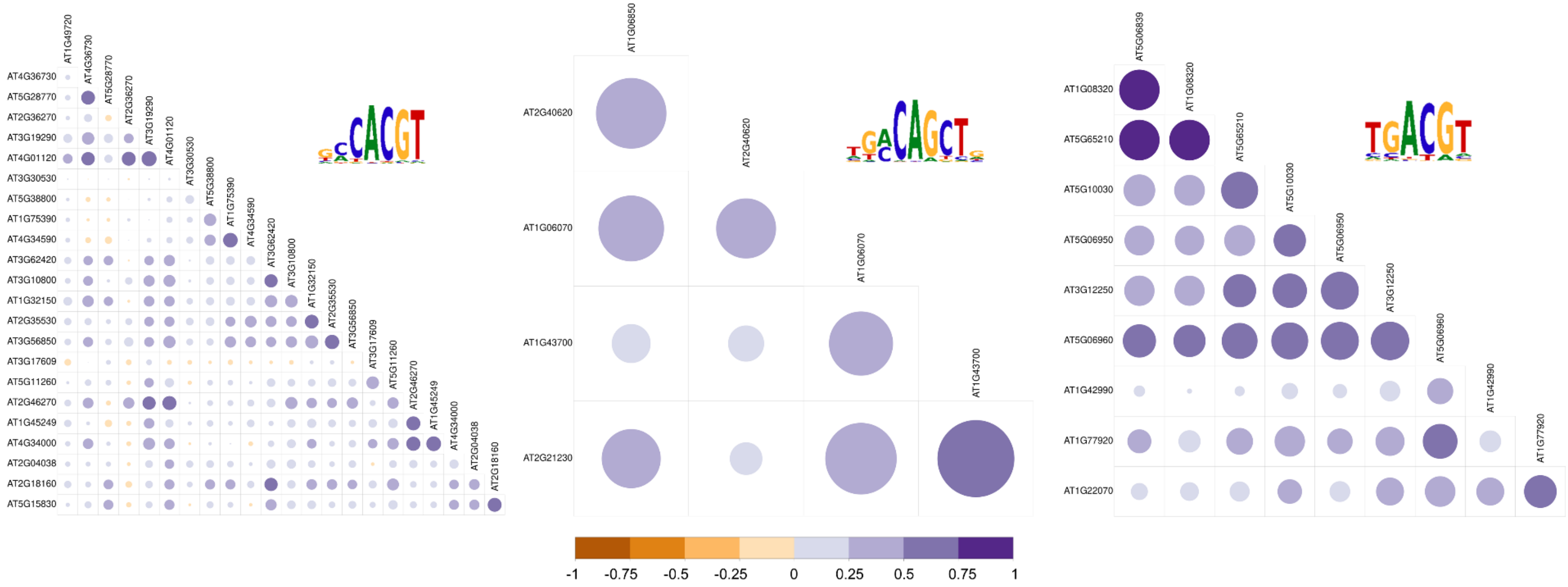


**Figure S51 Correlation matrices of bZIP TF expression within TFs binding the same TFBM in *A. thaliana***

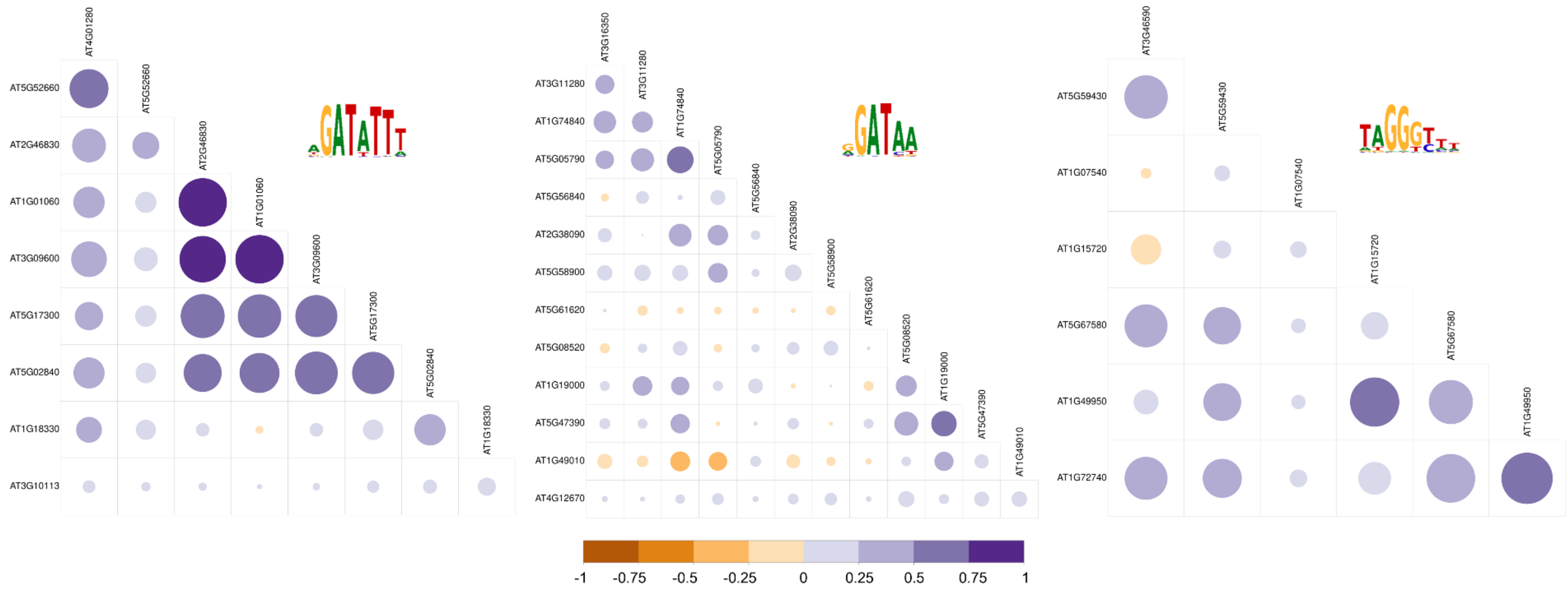


**Figure S52 Correlation matrices of MYB-related TF expression within TFs binding the same TFBM in *A. thaliana***

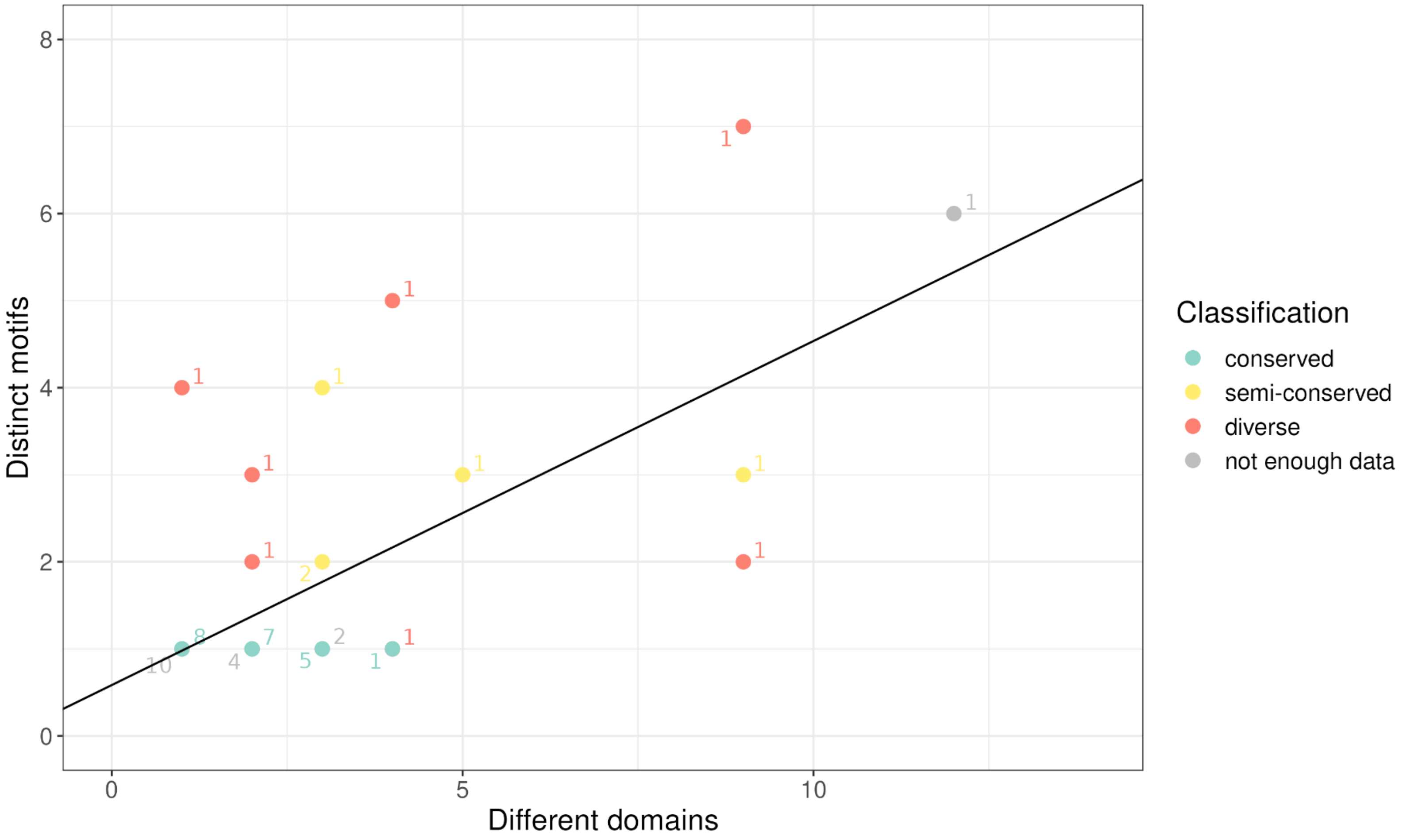


**Figure S53 Correlation of number of different domains annotated in a family versus number of distinct motifs.**

Linear regression with Pearson coefficient. Number of data points at each position labelled. Pearson correlation of 0.69.
